# FedPyDESeq2: a federated framework for bulk RNA-seq differential expression analysis

**DOI:** 10.1101/2024.12.06.627138

**Authors:** Boris Muzellec, Ulysse Marteau-Ferey, Tanguy Marchand

## Abstract

Large-scale transcriptomic studies are often limited by data silos and risks of privacy leakage, which may lead to missed clinical insights. Meta-analysis methods may be used to aggregate local results, but they induce lower statistical power and are particularly sensitive to heterogeneous settings. A recent paradigm in distributed computing, federated learning (FL) is a means of fitting models from siloed data, while ensuring that private data does not leave its storage facilities. Here, we introduce `FedPyDESeq2`, a software for differential expression analysis (DEA) on siloed bulk RNA-seq. Building on FL tools, `FedPyDESeq2` implements the `DESeq2` pipeline for DEA on siloed datasets in a privacy-enhancing manner. We benchmark `FedPyDESeq2` on datasets from The Cancer Genome Atlas corresponding to 8 different indications, split by geographical origin. `FedPyDESeq2` achieves near-identical results on siloed data compared with `PyDESeq2` on pooled data, and significantly outperforms meta-analysis baselines.

---

Differential expression analysis (DEA) is the primary application of RNA-seq, whether at bulk or single-cell resolution [1]. From sequenced transcriptomic profiles, DEA aims to identify differentially expressed genes (DEGs) between two conditions (e.g., treated and control samples), using statistical software such as `edgeR` [2], `limma` [3] or `DESeq2` [4] and its recently proposed python alternative, `PyDESeq2` [5].

To obtain relevant results, sample size is a critical factor. Several studies [6, 7] have come to the conclusion that replicate numbers have a larger impact than library sizes on the power of DE analyses. This is especially the case in the context of high sample variability, as pointed out in [1]: “for highly diverse samples, such as clinical tissue from cancer patient tumours, many more replicates are likely to be required in order to pinpoint changes with confidence.”

Putting aside the costs of RNA sequencing, one of the major obstacles to the existence of larger cohorts is the fact that genomic data is very sensitive, and its exchanges are tightly regulated by privacy-preserving legal constraints. As a result, DEA is often limited by the quantity of RNA-seq data held by a single institution. When data cannot be shared between institutions, meta-analysis [8] may be employed to aggregate the local results of independent analyses performed in each data center. However, this comes at the cost of statistical power, especially in the presence of data heterogeneity [9].

Recently, federated learning (FL) [10, 11] has emerged as a promising technology to face the challenges of siloed data. FL is a distributed computing paradigm originating from the machine learning (ML) community, which allows clients (or centers) to build ML models as if the data were pooled together while keeping control of their data. Schematically, it consists in repeating the following steps, under the supervision of a central server:

1. Clients perform computations on their private data.
2. Clients transfer their local models to the server.
3. The server aggregates models to form a single global model.
4. The server shares the model back to clients.

Recently, several works have applied FL to biomedical tasks to overcome the privacy constraints inherent in medical data. For example, FedECA [12] introduces a FL inverse probability of treatment weighting (IPTW) method as a means to implement external control arms. FL has also been employed for drug discovery [13], or to predict histological response to neoadjuvant chemotherapy in triple-negative breast cancer [14]. Finally, Flimma [15] aims to reproduce the limma-voom [3] DEA workflow in an FL setting. To the best of our knowledge, Flimma is the only federated framework for DEA so far, which contrasts with the diversity of available DEA software. In particular, widely popular tools such as `edgeR` [2] and `DESeq2` [4] were still lacking an FL version at the time this paper was being written.

In this paper, we introduce `FedPyDESeq2`, a software for federated DEA which aims to reproduce the results of `DESeq2` [4] in the presence of data silos. `FedPyDESeq2` is based on `Substra` [16], an open-source FL software hosted by the Linux foundation for artificial intelligence and data, and on `PyDESeq2` [5], a python re-implementation of the original R `DESeq2` package [4]. The goal of `FedPyDESeq2` is to output, from siloed bulk RNA-seq data, results that are as close as possible to those which would have been obtained if `PyDESeq2` had been applied after pooling the data together. To do so, `FedPyDESeq2` follows the `DESeq2` pipeline, implementing an equivalent version of each step, while ensuring that no individual data point leaves its center.

We evaluate `FedPyDESeq2` on bulk RNA-seq datasets from The Cancer Genome Atlas (TCGA, https://www.cancer.gov/tcga) corresponding to 8 different indications. To generate an FL setting, we split the data according to two different methodologies. First, following [17], we recover realistic federated datasets by splitting the data according to their sampling source sites, which we group into 7 regions. Second, to further benchmark the robustness of our approach, we generate a suite of synthetic splits of increasing heterogeneity. We then run `FedPyDESeq2` on a federated network in which each split is stored in an independent machine. In both settings, our experiments show that `FedPyDESeq2` yields results that are extremely close to those of `PyDESeq2` on pooled data. Finally, we compare `FedPyDESeq2` with various meta-analysis methods, which produce global results by aggregating local DEA results from each center. Our experiments show that using a federated method instead of meta-analysis leads to improved statistical power and sensitivity.

`FedPyDESeq2` is available at https://github.com/owkin/fedpydeseq2 under an MIT license. The scripts to reproduce our experiments are available at https://github.com/owkin/fedpydeseq2-paper-experiments.

## Results

### Overview of FedPyDESeq2

`FedPydDESeq2` is a python software for differential expression analysis on siloed bulk RNA-seq data. `FedPydDESeq2` operates on a network consisting of a set of centers, each with their own private RNA-seq data and sample covariates, and a central server to orchestrate them. For each gene, `FedPyDESeq2` derives a log-fold change (LFC) parameter and a p-value. Computations are performed without any exchange of raw individual RNA-seq or covariate data over the network.

`FedPyDESeq2` is built on `Substra` [16], an open-source FL framework which has already been applied in federated biomedical studies [13, 14]. `Substra` can be deployed in real-world federated settings, but also supports a simulated mode which can be run on a single machine.

From a bird’s eye view, `FedPyDEseq2` consists of a succession of two types of routines (see Fig. 1):

**Fig. 1.**
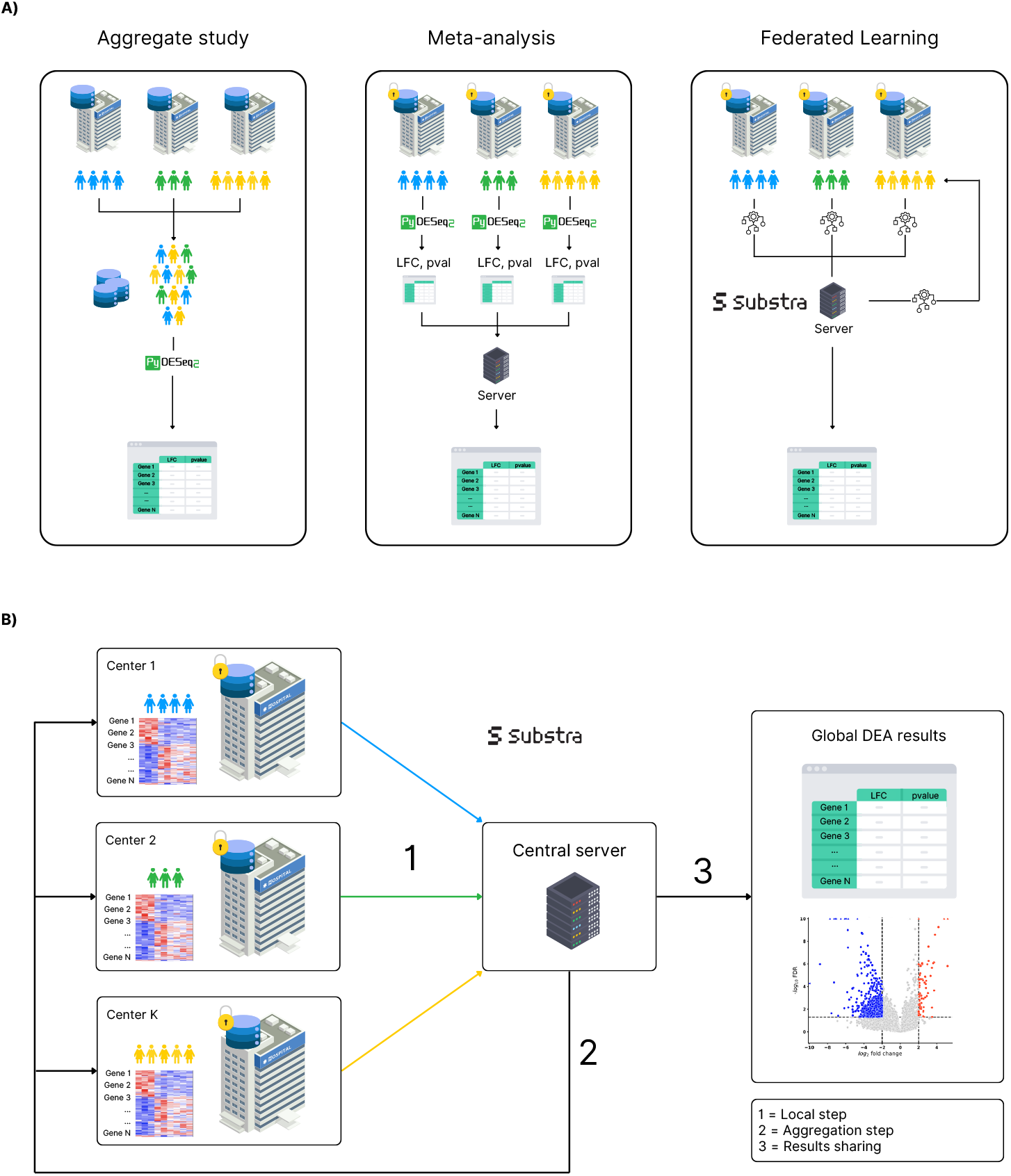
**(A)** Comparison of aggregate analysis (left), meta-analysis (middle) and federated learning (right) methods for DEA on siloed RNA-seq data. While aggregate analysis is the preferred method, it requires pooling data from all centers in a single processing center, which is often unfeasible. Meta-analysis circumvents this issue by running a full DEA pipeline locally on each center, sharing the resulting log-fold changes and p-values to a server for aggregation, at the loss of statistical power. FL, on the other hand, aims to reproduce the algorithm of the aggregate study by distributing the computation over a network, without private data leaving their premises. **(B)** `FedPyDESeq2` alternates between (1) local steps run independently in each center and (2) `aggregation steps` run centrally by the server. This is done for multiple rounds over a federated network orchestrated by `Substra`, before (3) the server outputs final results. To ensure privacy, centers may only share aggregate data.

1. `Local steps`, run independently by each center, during which centers may use their own private data in addition to quantities shared by the central server. The results are then stored locally in each center, and part of them may be transferred to the central server in the form of `shared states`;
2. `Aggregation steps`, run on the central server, using the `shared states` sent by the centers. The output of aggregation steps is then shared back to the centers.

The succession of `local steps` and `aggregation steps` is statically fixed in advance and communicated to the centers and the server by `Substra`. Formally, `FedPyDESeq2` follows the same mathematical procedure as `DESeq2` [4]. Raw counts are modeled as independent negative binomial distributions. Dispersion parameters are first estimated independently for each gene by fitting a generalized linear model (GLM). They are then shrunk towards a global trend curve, to pool information among genes. Next, dispersions are used to fit gene-wise log-fold changes (LFC), which encode the differences in expression levels between conditions. Outliers are then detected and removed using Cook’s distances [18]. Finally, p-values are computed from LFCs and dispersions using Wald tests, and adjusted to control the false discovery rate.

`FedPyDESeq2` is structured along the same architecture as `DESeq2`, but implements each step in a federated manner, which requires a few adjustments. `FedPyDESeq2` fits dispersions parameters using grid search instead of `l-bfgs-b` [19], which is less amenable to federation. To fit LFCs, we implement a federated version of the iteratively reweighted least squares algorithm [IRLS, see, e.g., 20]. A final challenge lies in the computation of Cook’s distances for outlier detection [18], for which we design a federated quantile computation procedure.

To prevent data leakage, `FedPyDESeq2` adheres to the following privacy guideline: data at the sample or patient level, such as individual RNA transcripts, design covariates or size factors, are considered private and are never shared. On the other hand, gene-level information that is part of the output of the algorithm (such as dispersions, LFCs or p-values) is freely exchanged between the centers and the server.

The features currently available in `FedPyDESeq2` cover the majority of DEA cases. `FedPyDESeq2` supports multi-factor designs for both categorical and numerical covariates, the refitting of outliers based on Cook’s distances, p-value computation using Wald tests, and p-value adjustment using the Benjamini-Hochberg procedure and independent filtering (see [4] for details). In particular, `FedPyDESeq2` makes it possible to include center information as design covariates and to compare cohorts that are perfectly partitioned between data silos, which is completely impossible for meta-analysis methods. This enables users to alleviate batch effects between centers or to compare the effects of different treatment protocols performed in a given set of hospitals.

We refer to the Methods section and the supplementary material for more details on `DESeq2`, `PyDESeq2` and `FedPyDESeq2`, including a detailed account of the quantities shared between the centers and the server during the `FedPyDESeq2` workflow.

### FedPyDESeq2 yields near-identical results on siloed RNA-seq data compared with PyDESeq2 on pooled data

The goal of `FedPyDESeq2` is to obtain LFCs and p-values that are as close as possible to those that would be output by `PyDESeq2` on the same data, if it were pooled together in a single database. We evaluate this by comparing the outputs of both methods on RNA-seq datasets from The Cancer Genome Atlas (TCGA, https://www.cancer.gov/tcga) corresponding to 8 different indications.

In a first set of experiments, we simulate a realistic federated scenario by splitting datasets according to the geographical origin of each sample (available in the metadata), resulting in 7 sub-datasets for each indication. For each indication, we run `FedPyDESeq2` on a network of 8 machines, comprising a server and 7 clients (one for each sub-dataset), and compare the results with those obtained by running `PyDESeq2` on the original (undivided) data on a single machine. This experiment is repeated with three different designs, using covariates corresponding to clinical data from TCGA: (i) a single-factor design, (ii) a multi-factor design with categorical factors, and (iii) a multi-factor design which includes a numerical factor. The results for design (i) are summarized in Fig. 2.

**Fig. 2.**
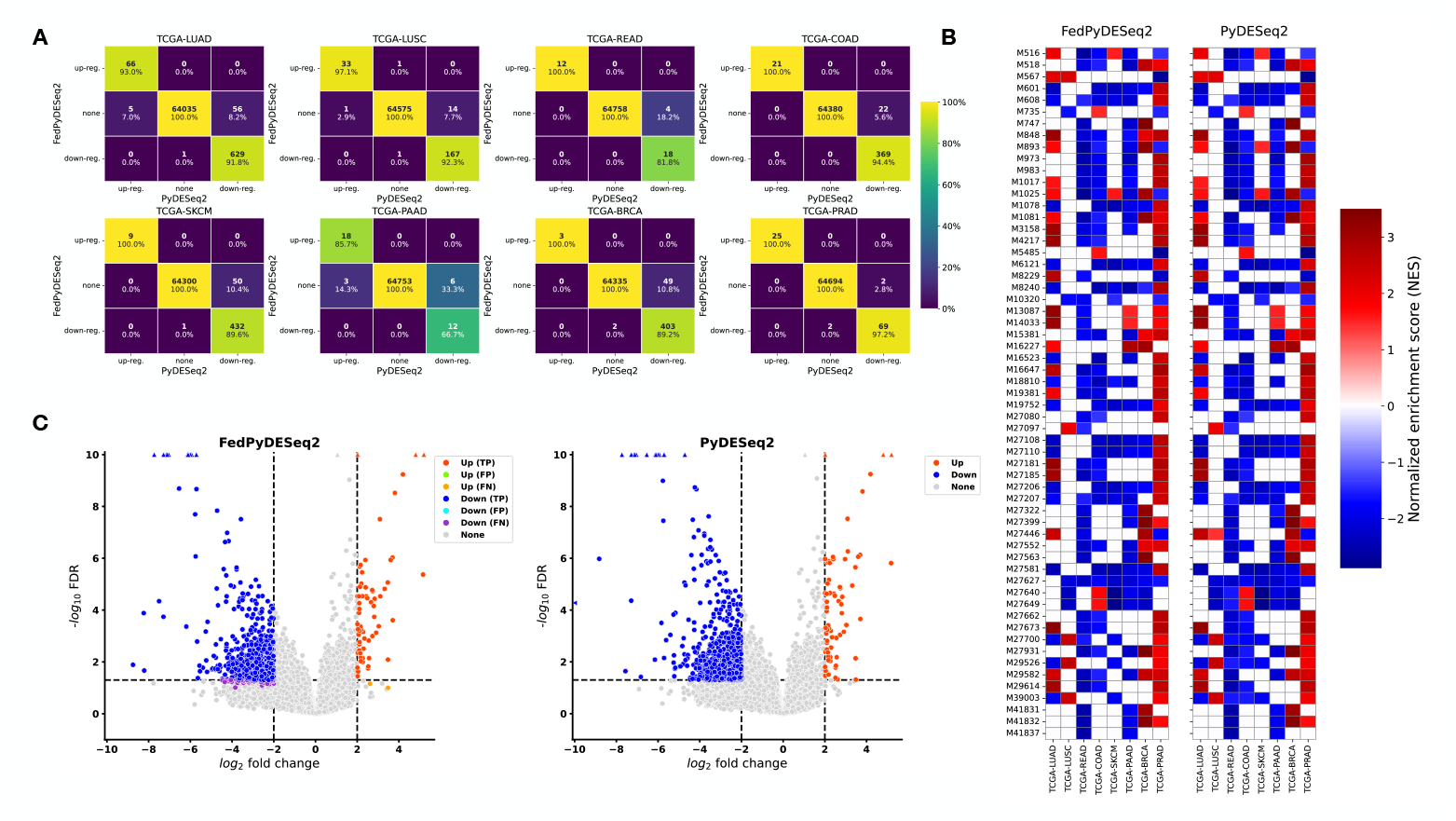
Results of the single-factor design experiment (`Advanced` vs `Non-advanced` tumoral stage) with a geographical split. **(A)** DEGs (with padj ≤ 0.05 and |LFC| ≥ 2) according to `FedPyDESeq2` (from siloed data) and `PyDESeq2` (from pooled data). Percentages are expressed w.r.t. column totals. **(B)** Significantly enriched pathways (padj ≤ 0.05) obtained with the `fgsea` package, using Wald statistics as gene-ranking metric. Only pathways that are in the top 10 (according to the adjusted p-value) for at least one indication are represented. If for a given indication, a pathway is not significantly enriched, the corresponding square is left blank. **(C)** Volcano plots of DEGs in TCGA-LUAD (left: `FedPyDESeq2`, siloed data; right: `PyDESeq2`, pooled data). Differences in DEGs are marked as false positives/negatives (FP/FN) on the `FedPyDESeq2` plot.

Across all 8 indications, `FedPyDESeq2` retrieves the vast majority of `PyDESeq2` DEGs. Looking closer, it appears that the differences are mainly due to genes identified as DE by `PyDESeq2`, but not `FedPyDESeq2`. Overall, `FedPyDESeq2` tends to be less sensitive than `PyDESeq2`, but seldom introduces new DEGs. More precisely, differences in DEGs are mainly due to p-values that are close to the 0.05 threshold, as can be seen in Fig. 2-C. We further validate the biological soundness of our results by performing gene set enrichment analysis (GSEA) using the `fgsea` package [21] on both sets of DEGs. As shown in Fig. 2-B, this results in near-identical sets of significantly enriched pathways.

We refer to the Methods section for more details on the setting of our experiments, and to the supplementary material for additional results.

### FedPyDESeq2 significantly outperforms local methods on siloed RNA-seq data

We compare `FedPyDESeq2` to a set of meta-analysis baselines [8] which can be run on siloed data without sharing local transcriptomic or covariate data. The meta-analysis methods benchmarked in our work include: (i) a fixed-effects model, (ii) random-effects models, with either the Paule-Mandel [22] iterative estimator or the DerSimonian and Laird one-step estimator [23], and (iii) p-value combination with Stouffer and Fisher methods (see, e.g., [24]). We also include a naive method which consists in restricting the analysis to the largest local cohort. For all methods, we use the results of `PyDESeq2` applied to the pooled dataset as reference.

In addition to the geographical split, we introduce a set of synthetic splits to further evaluate the robustness of our methods w.r.t. data heterogeneity. To do so, we group two colorectal cancer (CRC) datasets together, TCGA-COAD and TCGA-READ. We then simulate various scenarios of two centers containing CRC patients data with increasing levels of heterogeneity. At the lowest level of heterogeneity, each of the two datasets contains the same proportion of patients from TCGA-COAD and TCGA-READ, while at the highest level of heterogeneity, one of the datasets only contains TCGA-COAD patients and the other only TCGA-READ. We apply the same procedure to two non-small cell lung cancer (NSCLC) datasets: TCGA-LUAD and TCGA-LUSC.

We perform our benchmark on both the heterogeneous splits and on the geographical split introduced previously. The results of our experiments are summarized in Fig. 3. While in both CRC and NSCLC, `FedPyDESeq2` retrieves about 93% of `PyDESeq2` DEGs, meta-analysis methods perform much worse all-around, which makes them a poor substitute for federated methods. Overall, the low performance of meta-analysis is mostly due to poor sensitivity (i.e., most DEGs are missed), with the exception of the DerSimonian and Laird random-effects model [23], which also introduces many false positives. Second, whereas `FedPyDESeq2` is insensitive to data heterogeneity, the recall of meta-analysis deteriorates sharply as heterogeneity increases, i.e., fewer and fewer DEGs are correctly identified. As a notable exception, the performance of `PyDESeq2` restricted to the largest CRC cohort increases with heterogeneity. However, we believe that this phenomenon is an artifact due to the unbalanced numbers of replicates between TCGA-COAD and TCGA-READ, resulting in a maximum number of replicates in the largest cohort at the maximum heterogeneity level.

**Fig. 3.**
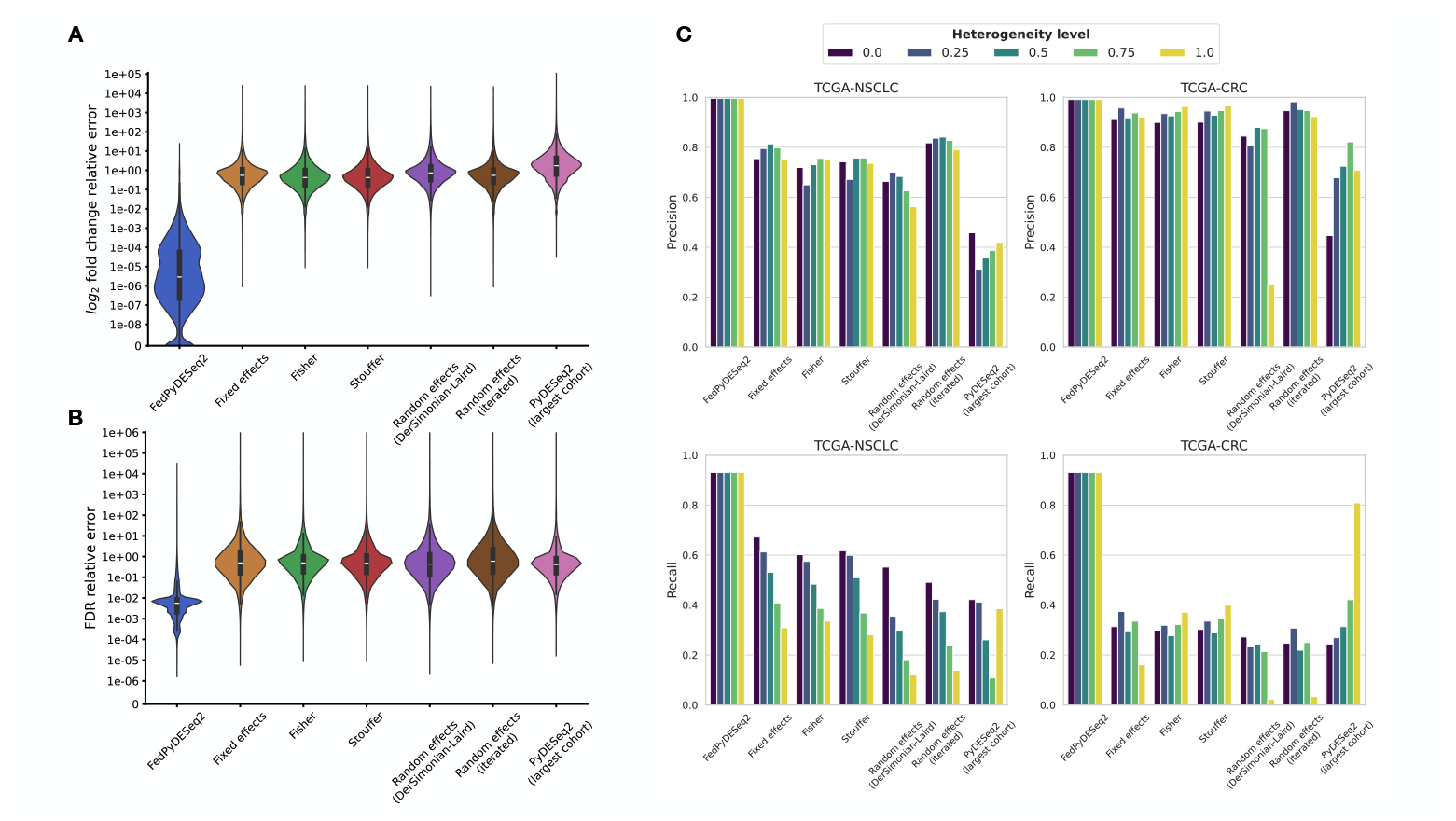
Comparison of `FedPyDESeq2` with meta-analysis on siloed data. **(A)** LFC and **(B)** adjusted p-value relative errors of `FedPyDESeq2` and baseline methods applied on geographically split data, compared with `PyDESeq2` on pooled data 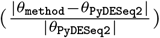, evaluated on TCGA-LUAD. **(C)** Precision and recall scores of `FedPyDESeq2` and baselines on siloed TCGA-NSLCL and TCGA-CRC data with increasing heterogeneity, compared with `PyDESeq2` on pooled data.

We refer to the Methods section for more information on experiments, including a detailed description of splitting procedures and meta-analysis baselines, and to the supplementary material for a finer-grain report of each method’s output.

## Discussion

`FedPyDESeq2` is dedigned to address the issue of data silos in bulk RNA-seq DEA, which limits the size of cohorts. `FedPyDESeq2` leverages FL to perform DEA without sharing individual patient data, such as RNA transcripts or design covariates, relying instead on data aggregation. Our experiments show that the results of `FedPyDESeq2` applied on siloed data are very close to those `PyDESeq2` would yield if it were applied on pooled data. Further, `FedPyDESeq2` strongly outperforms methods based on the aggregation of local (shareable) `PyDESeq2` results, such as fixed or random-effects models, or p-value combination methods.

`FedPyDESeq2` opens the way for DEA studies which would be completely impossible to perform without pooling the data together. Indeed, `FedPyDESeq2` allows comparing cohorts that are perfectly partitioned across data silos. As an example, it could be used to determine whether patients from a hospital A have DEGs compared with patients from a hospital B. This enables comparing the effects of treatment protocols that are performed exclusively in a given hospital. On the other hand, such an analysis would be intractable using meta-analysis, as it may only aggregate local results, in which the comparison of A patients with B patients is impossible. The same strategy can be employed to mitigate heterogeneity between centers, by adding a center-wise covariate to the design.

`FedPyDESeq2` is based on federated learning, a privacy-enhancing technology that ensures that raw data stays within each center and is not shared across the network. However, on a cautionary note, various studies [11, 25, 26] have shown that privacy leakage can still occur, as intermediate quantities exchanged over the network may contain private information. In the case of machine learning models trained using FedAverage [10, 11], an attacker could in some conditions reconstruct the local data from the gradients exchanged over the network. In `FedPyDESeq2`, the quantities shared with the server consist only in aggregated read counts statistics, such as log-means or variances, or Gram or hat matrices built from the design covariates. We provide an exhaustive list of those quantities in the supplementary material. Although we leave a detailed study evaluating precisely the risks of data leakage to future work, we believe that as long as each center has a sufficient number of samples, an attacker cannot reconstruct initial gene counts from the quantities exchanged over the network.

Further privacy-enhancing technologies can be applied on top of FL in order to strengthen the privacy guarantees. As an example, differential privacy [27, 28] provides theoretical guarantees on privacy leakage by adding random noise to the initial data. However, in doing so, it alters the final output of the FL algorithm. Another privacy-enhancing layer that can be added is secure aggregation [29], which relies on cryptography methods to hide center-level shared states from the server. However, while secure aggregation leaves the outputs of FL algorithms unchanged, it induces strong computational overheads.

A critical aspect of federated learning is the heterogeneity that is inherent to different data sources [11]. In this work, our objective is to produce results that are as close as possible to those that would have been obtained from pooled analysis, and to show the robustness of `FedPyDESeq2` with respect to heterogeneous data silos. In practice, more relevant results could be obtained by taking heterogeneity into account. As advocated in the `DESeq2` vignette, this can either be achieved by including batch variables (e.g., hospital A vs hospital B) in the design of `FedPyDESeq2`, or by using dedicated batch effect correction tools before we perform the analysis, such as `RUVSeq` [30], `SVASeq` [31], `ComBat` [32] and `ComBat-Seq` [33]. In that perspective, a federated version of `ComBat`, `Fed-Combat` [34], was recently proposed.

Finally, let us remark that DEA is only one of the many steps involved in most RNA-seq analysis pipelines. Hence, to build a complete pipeline, federated software handling complementary tasks such as clustering is required (see, e.g., [35] and [36] for federated principal component analysis). More importantly, let us highlight the importance of applying unified sample processing across all centers (library preparation, sequencing technology, read files preprocessing and alignment) when using `FedPyDESeq2`, to ensure that the transcripts which are being compared correspond to the same biological entities.

## Supporting information

Additional Results

Workflow Diagram

## Methods

In this section, we provide a comprehensive understanding of `FedPyDESeq2` and its evaluation.

First, we describe the `DESeq2` methodology. We start with the underlying statistical model before detailing the computational pipeline. This provides the necessary context to understand how we adapt each component to the federated setting.

Next, we present our main contributions: the `FedPyDESeq2` methodology and its implementation. We explain how each step of the `DESeq2` pipeline is adapted to work with distributed data without sharing the data at the server level.

Finally, we detail the experimental setup we used to i) validate the performance of `FedPyDESeq2` against the `(Py)DESeq2` reference and ii) compare it with other methods which do not centralize the data. We begin by describing meta-analysis approaches, which serve as our primary baselines. We then describe the data we used to run our benchmark, and the evaluation methodology. We conclude with a brief overview of our experimental infrastructure and deployment considerations.

### The DESeq2 model

In `DESeq2` [4], the user provides:

- a gene count matrix *Y* = (*y*_*ig*_) ∈ ℝ^*n×G*^, corresponding to *n* input samples and *G* genes (*y*_*ig*_ is the gene count of gene *g* for sample *i*);
- a design matrix *X* ∈ ℝ^*n×p*^, corresponding *n* input samples and *p* design factors (**x**_*i*_ ∈ ℝ^*p*^ will denote the covariate vector of sample *i*, i.e., the *i*-th line of the design matrix). For each gene *g*, `DESeq2` models gene counts 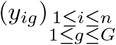 as **independent** samples from negative binomials, parameterized by their means and dispersion parameters:

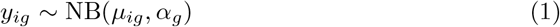

where
- the dispersion parameter *α*_*g*_ depends only on gene *g*;
- the mean parameter *µ*_*ig*_ depends on the covariate vector **x**_*i*_ ∈ ℝ^*p*^, the size factor *γ*_*i*_ of sample *i* (provided or computed in an early step of the pipeline), and the log fold change (LFC) vector *β*_*g*_ ∈ ℝ^*p*^ of gene *g*, as a generalized linear model (GLM): *µ*_*ig*_ = *µ*_*i*_(*β*_*g*_), where

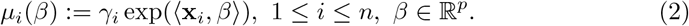

Let 𝓁^*NB*^(*y*|*µ, α*) be the negative log likelihood (NLL) of the negative binomial distribution with parameters *µ* and *α* for a single sample *y*^1^. For each gene *g*, the NLL of the GLM can be written as

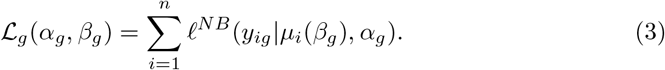

The high-level objective of `DESeq2` is to estimate the dispersion and log fold change parameters *α*_*g*_ and *β*_*g*_ for all genes, successively using method of moments techniques and alternatively minimizing the NLL ℒ_*g*_ (sometimes with prior regularization) w.r.t. *α*_*g*_ and *β*_*g*_.

### The DESeq2 workflow

We now briefly describe the main steps of the `DESeq2` workflow, to serve as a reference when we describe later how they are implemented in `FedPyDESeq2`. We refer to the original `DESeq2` paper [4] and its implementation (https://github.com/thelovelab/DESeq2) for further details.

### Size factors

`DESeq2` estimates size factors using the median of ratios method. Discarding genes which have at least one zero count value, we first compute a pseudo reference *R*_*g*_ for each gene *g* as the geometric mean of its counts across samples:

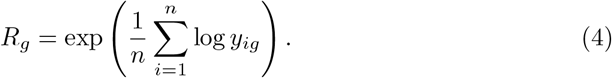

We then define the **size factor** *γ*_*i*_ of sample *i* as

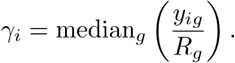

### Initial dispersion estimates

Initial values for dispersions are first estimated using a combination of a method of moments and a rough estimate from a linear regression.

#### Method of moments

Let 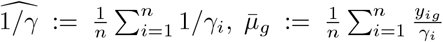 and 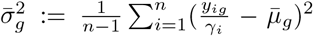. The method-of-moments estimate is defined as

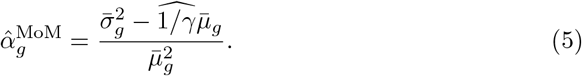

### Rough dispersions estimates

We first estimate 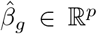 by fitting a linear regression predicting the normalized counts from the design factors:

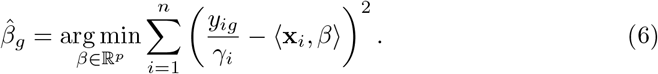

Let 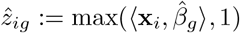. The “rough dispersion estimates” (RDE) are defined as:

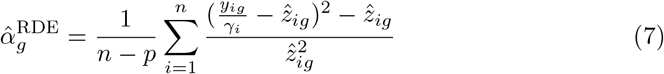

#### Initial estimates

Finally, the initial dispersions estimates are set as

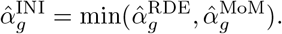

### Initial *µ* estimates

First estimates for means *µ*_*ig*_ are obtained in one of two ways.

#### Case 1

If the number of levels in the design matrix is equal to *p*, the estimates are obtained by fitting a linear model for the normalized counts,

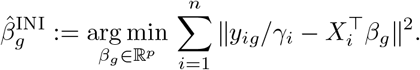

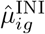 is defined as the smoothed value:

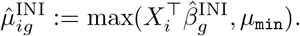

#### Case 2

In all other cases, the estimates are obtained by optimizing the log likelihood of the negative binomial w.r.t. LFCs, using iteratively re-weighted least squares (IRLS, see hereafter). Let

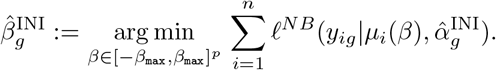

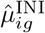 is defined as

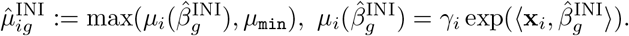

At this stage, only the 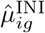 estimate is relevant, and log-fold changes 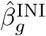 are discarded. *µ*_min_ and *β*_max_ are both standard `DESeq2` parameters with default values.

### Estimating dispersions

#### Gene-wise maximum likelihood estimates

Dispersions are first estimated independently for each gene using maximum likelihood estimation (MLE), i.e., by solving, for each gene, the following optimization problem:

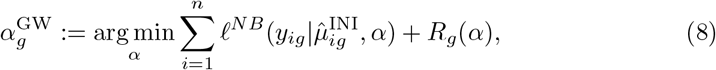

where *R*_*g*_(*α*_*g*_) is a Cox-Reid regularization term [37]:

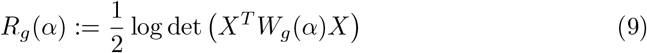

and

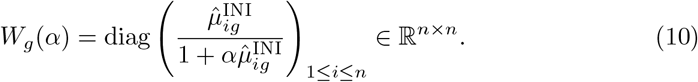

In practice, box constraints are applied to restrict *α*_*g*_ to lie in

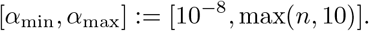

Numerically, the optimization is carried out on a log *α* scale, using line search (`DESeq2`) or L-BFGS-B [19] (`PyDESeq2`).

#### Fitting a trend curve to gene-wise dispersion estimates

Next, information is pooled across genes by fitting a curve relating the gene-wise dispersions *α*^GW^ to the inverse of the empirical normalized means 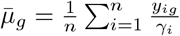:

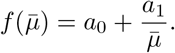

The parameters *a*_0_ and *a*_1_ are fitted with a Gamma-distributed generalized linear model, minimizing the following loss function:

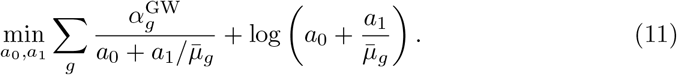

Equation 11 is fit iteratively, excluding genes for which the ratio between the actual value and the fitted curve value is outside of [10^−4^, 15] at each iteration.

In case the optimization does not converge, a constant (flat) curve is used instead, taking the means of *α*^GW^ over all genes, trimmed to remove the top and bottom 10^−3^ quantiles of dispersions, as value.

#### Dispersions variance prior

A dispersion variance prior is then estimated based on the squared mean absolute deviation (MAD) of the terms 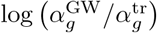(where 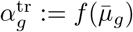), as

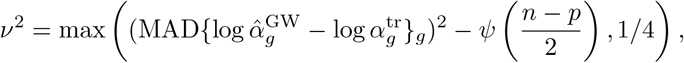

where *ψ* denotes the digamma function.

#### Maximum a posteriori (MAP) dispersions

MAP dispersion estimates are then fitted for each gene. The loss is similar to (8) with an additional regularization term to control the log-deviation from the fitted curve:

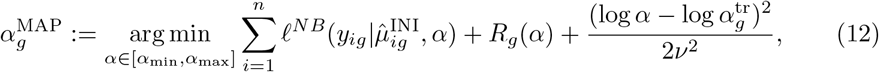

with 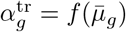. Again, the optimization is performed using line search in log domain, and dispersions are constrained to lie in [*α*_min_, *α*_max_].

#### Filtering outlier dispersions

Finally, when 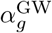 is too far above the trend function, i.e. when

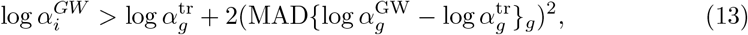

the 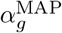 is not computed and 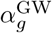 is used instead.

### Log fold change estimation

Based on the final dispersion estimates *α*_*g*_ obtained in the previous step, we can proceed to the log-fold change estimation. Recall from (1) that `PyDESeq2` assumes that the raw counts *y*_*ig*_ are sampled independently (over samples and genes) from negative binomial distributions : *y*_*ig*_ ∼ NB(*µ*_*ig*_, *α*_*g*_), where the mean parameter *µ*_*ig*_ is given as a generalized linear model over the covariate vector **x**_*i*_ (see (2)):

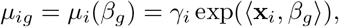

for some gene-dependent vector *β*_*g*_. Recall from (3) the definition of the negative log-likelihood ℒ_*g*_ over all samples for gene *g*:

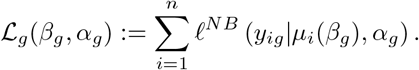

In order to estimate the log-fold change *β*_*g*_, `(Py)DESeq2` minimizes ℒ_*g*_(*β*_*g*_, *α*_*g*_) independently for each gene inside a box:

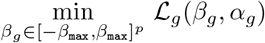

where *β*_max_ is a parameter of the algorithm, with a default value to avoid unrealistic log fold changes.

This optimization is carried out with the iteratively reweighted least-squares (IRLS) algorithm. For simplicity, we will drop the gene subscript *g* in the following description, and assume we are optimizing the function for a given gene.

From a bird’s eye view, IRLS boils down to iterating the following updates, starting from an initial LFC estimate *β*_0_:

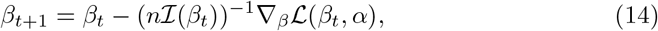

where ℐ(*β*) is the Fisher information, defined as the expectation over samples of the score function (gradient w.r.t. *β* of the sample-wise NLL):

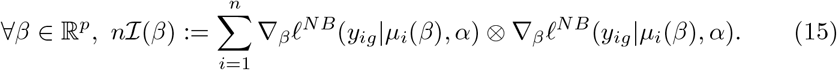

Here, the operator ⊗ denote the tensor product between two vectors, i.e., *a* ⊗ *b* = *ab*^⊤^. In other words, at each iteration, we need to solve

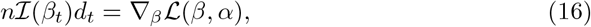

and apply

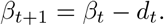

In the present setting, we can decompose the Fisher information as

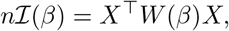

where *W* (*β*) is a diagonal matrix:

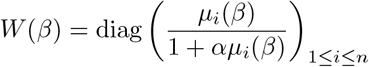

Whenever IRLS fails to converge, or moves out of the box constraints, L-BFGS-B [19] is used as an alternative.

### Detecting and replacing outliers

Cook’s distances [18] are used to assess whether any single read count value has had a disproportionately strong influence on the final dispersion and LFC values. In that case, this read count is replaced with a trimmed mean imputation, and dispersions and LFCs are refit.

#### Computing Cook’s distances

A method of trimmed moments is applied to estimate dispersions based on counts from which extreme values where removed :

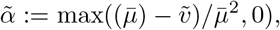

where 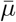 is the empirical mean (i.e., the average of normalized counts), and 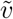 is a trimmed variance, i.e., the empirical variance after a fixed fraction of the largest and smallest counts were removed.

For a gene *g* and sample *i*, Cook’s distance for GLMs is then given by [see 4, 18]:

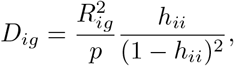

where

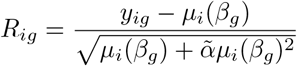

is the Pearson residual of sample *i* and *h*_*ii*_ is the *i*-th diagonal element of the hat matrix *H* (omitting the *β*_*g*_ parameter):

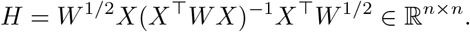

#### Replacing outliers

Then, for all genes *g* and samples *i* for which *D*_*ig*_ is greater than a given threshold (which depends only on the number of samples *n* and the number of parameters *p*), *y*_*ig*_ is replaced with the trimmed mean of *g*’s normalized counts over all samples, multiplied by *γ*_*i*_. Keeping the trend curve unchanged, dispersions and LFCs for all such genes are then refit, and the new values replace the old ones.

This is only applied to samples whose condition (i.e. whose covariates value combination) meet a minimal number of replicates (by default, 7).

### Computing and adjusting p-values

#### Wald tests

A p-value is computed for each gene using a Wald test, which consists in comparing the LFC coefficient of interest *β*_*g,r*_, divided by its standard error 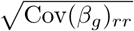, to a normal distribution (multiplied by 2 to achieve a two-tailed test). The following covariance matrix estimator is used (see [4, 38] for details):

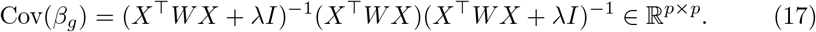

#### P-value Cook filtering

Cook’s distances are then used again to filter out p-values for genes which have remaining outlier count values. More precisely, genes which still have at least one sample with a Cook’s distance above the threshold have their p-value set to `NaN`, unless there are 3 samples or more whose read counts are above those of the maximum Cook’s distance sample.

#### P-value adjustment

Finally, p-values are adjusted by applying the Benjamini-Hochberg procedure [39] and independent filtering (see [4]). We do not detail these steps here: in a federated setting, they can be performed by the server using quantities from previous steps. Hence, they do not require any federation effort.

### Federating DESeq2

Let us now assume that the data is partitioned into *K* different centers {1, …, *K*} which cannot share their data. Let *n*_1_, …, *n*_*K*_ be the number of data samples in each center, *X*^(1)^, …, *X*^(*K*)^ the corresponding design matrices, and *Y* ^(1)^, …, *Y* ^(*K*)^ the corresponding count matrices. In particular, for 1 ≤ *k* ≤ *K*, we have 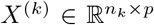 and 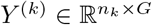. Similarly to the classical setting, we denote by 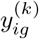 the count for gene *g* of the *i*-th sample of the *k*-th center, and by 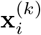 the covariate vector of the *i* -th sample of the *k*-th center.

The objective remains the same: from input designs and counts (*X*^(*k*)^, *Y* ^(*k*)^), 1≤ *k* ≤ *K*, we wish to estimate a dispersion *α*_*g*_ ∈ ℝ_+_, a log fold change vector *β*_*g*_ ∈ ℝ^*p*^ and a p-value in (0, 1) for each gene *g*. However, we now have the additional constraint that centers 1, …, *K* are not allowed to share individual data between themselves or with the server. In particular, this implies that neither the full design matrix nor the full counts matrix may be formed.

As in most FL settings, we will overcome this obstacle by leveraging the fact that the negative likelihood function (NLL), which we minimize to fit dispersions and LFCs, is *separable*: the total NLL is the sum of the negative binomial NLL on each sample. This implies that the total NLL can be expressed as the sum of each center’s local NLL. Indeed, given a gene *g*, it holds:

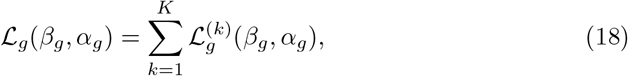

where 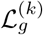 is the NLL on the samples of center *k* and ℒ_*g*_ is the NLL on all samples. Similarly to (2), defining the mean function as

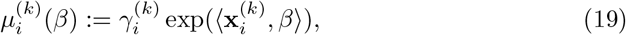

we can write the NLL of center *k* as

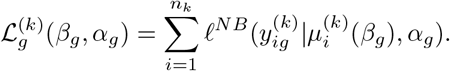

From (18), we can compute the gradient of the NLL (w.r.t either dispersions or LFCs) by aggregating (summing) local gradients.

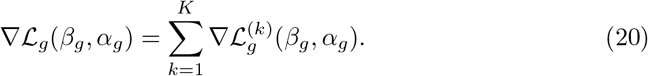

### Federating size factor estimation

We obtain the pseudo reference sample (Equation 4) by computing a federated mean of log-counts: each center *k* communicates 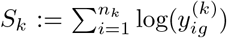 to the server, which then forms

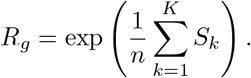

Finally, the server shares back *R*_*g*_ to each center *k*, which can then locally compute their size factor for each sample 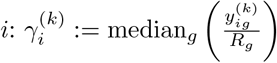.

### Federating initial dispersions estimates

Method-of-moments estimates (Equation 5) are derived by computing global means using the same strategy as for size factors (c.f. above).

To compute “rough dispersion estimates” for a given gene *g*, we start by computing the solution 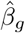 to Equation 6. To do so, we note that 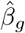 is the solution to a linear system 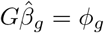 where

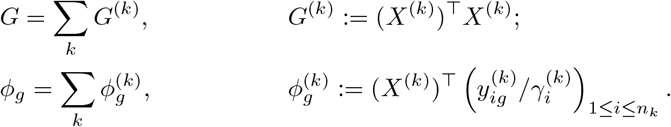

*G*^(*k*)^ and 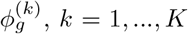, can be computed in each center, and then shared to the server to compute *G* and *ϕ*_*g*_, and hence 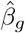. From there, rough dispersion estimates are obtained by computing federated means (c.f. Equation 7).

### Federating initial *µ* estimates

As mentioned in the `DESeq2` workflow section, initial *µ* estimates are obtained using IRLS. We refer to the subsection on federating LFC computation for a federated version of IRLS.

### Fitting dispersions in a federated setting

In `DESeq2`, the dispersions are fitted by minimizing the NLL with fixed log fold changes twice: in the genewise dispersions step, and in the MAP dispersions step. In `DESeq2`, these are fitted by minimizing the NLL using line search. As an implementation of line search in a federated setting is challenging, we instead resort to a simple grid search.

The choice of a grid search is motivated by three main facts:

- the underlying optimization problem is one-dimensional;
- although its objective function is not convex (because of the Cox-Reid term, equation 9), we empirically observe that it has a single optimum;
- in a federated setting, computational costs are dominated by communication costs.

Indeed, since the optimization problem is one-dimensional, performing a grid search is a relatively effective way of finding the minimum.

In the rest of this section, we omit the genewise notation *g*: we describe the process of computing a single dispersion. The computing of the dispersions is then parallelized on all genes, yielding the same communication cost. Recall that in this step, the log-fold change parameter *β* is fixed. To apply a grid search in a federated setting, we must compute the global adjusted NLL (i.e. the NLL plus the Cox-Reid adjustment term, see equation 8) evaluated on the whole federated data at each point of the grid.

#### NLL term

The NLL term can be straightforwardly evaluated leveraging its data separability: if each center computes its local ℒ ^(*k*)^(*β, α*) for all ℒ*α*’s on the grid and communicates it to the server, the server may in turn derive the global ℒ (*β, α*) values by summation (see equation 18).

#### Cox-Reid adjustment term

The Cox-Reid adjustment term (equation 9), however, cannot be obtained directly from a simple summation. Instead, the server computes the global argument of the log-determinant (CRM for Cox-Reid matrix), which can be decomposed over the *K* centers:

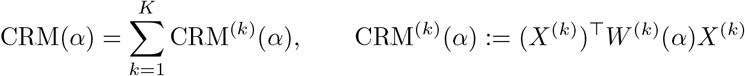

and where *W* ^(*k*)^ is a diagonal matrix defined as in (10). Each center computes CRM(*α*) matrices at each point of the grid and communicates them to the server, which sums them to obtain CRM(*α*) before applying the log-determinant.

The aggregator then sums all the terms and computes the global loss on all grid points in [*α*_min_, *α*_max_], and selects the dispersion on the grid point 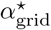 which minimizes the loss. This value is then refined by re-iterating the process: a new grid is defined on 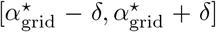, on which a new optimum is sought. By default, `FedPyDESeq2` repeats this process three times on grids of size 100, in logarithmic space.

From a computational point of view, this is very cost-effective: the cost of locally computing 100 adjusted NLL for a batch of genes is negligible compared with the cost of transferring data back and from the centers to the server. By limiting these exchanges to three rounds, iterative grid search achieves much faster convergence than more classic gradient-based FL methods such as FedAvg [10], and does not need any hyper-parameter specification.

#### Gene-wise estimates

Gene-wise estimates *α*^GW^ are derived using the above strategy.

#### Fitting a trend curve to gene-wise dispersion estimates and computing the variance prior

This step does not require any particular federation effort: it can be performed on the server from the gene-wise estimates and means of normalized counts.

#### MAP estimates

As for gene-wise estimates, MAP estimation is performed using grid search, with an additional 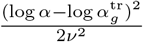 shrinkage term. This term is computed directly on the server from the trend curve dispersion estimates 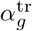 and the prior variance *ν*.

### Fitting log-fold changes in a federated setting

In `DESeq2` [4], two steps require the log fold changes *β*_*g*_ to be fitted with negative log-likelihood minimization:

- to obtain initial 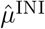 estimates;
- to compute the final log fold changes, estimated from the final dispersions.

`DESeq2` performs the optimization using IRLS, and switches to LBFGS in case of failure. In the federated setting, we adopt the same approach. We first try to apply the federated IRLS algorithm to all genes. This algorithm is pooled-equivalent: it outputs exactly the same results as in the pooled setting (see the next paragraph). However, for genes for which IRLS has not converged, we do not use LBFGS or LBFGS-B, which are both harder to federate and more adapted to a setting where communication bottlenecks do not exist. Instead, we use a Proximal Quasi Newton method [40] with a greedy line search strategy in order to minimize the number of communication rounds, which is the main bottleneck for cross-silo FL.

#### Federating the IRLS algorithm

In a nutshell (c.f. the `DESeq2` workflow section), IRLS consists in iteratively solving

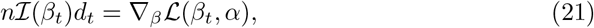

for *d*_*t*_, and applying

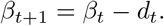

In a federated setting, we cannot obtain the update *d*_*t*_ as the sum of local values: indeed, ℐ (*β*_*t*_)^−1^ is not separable (i.e. linearly decomposable w.r.t centers). Instead, at each iteration, we form *n*ℐ (*β*) and ∇_*β*_ℒ (*β, α*) on the server (which is possible as they are separable), and solve equation 16. The separability of the gradient term comes directly from the separability of the NLL (equation 20):

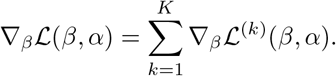

The separability of the Fisher information term comes directly from (15) since it is a sum over the samples: it can be written as 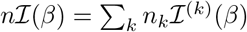, where

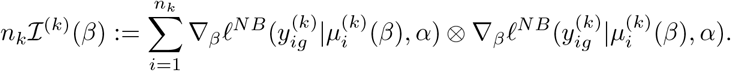

It is therefore possible to federate an IRLS step in a single round of communication (see Algorithm 1).

In the actual implementation of the IRLS algorithm, there are two possible stopping criteria:

- A convergence stopping criterion: if the deviance ratio is smaller than a threshold value, the algorithm is said to have converged;
- A divergence criterion: if the update *β*_*t*_ gets out of the box constraints set by the user or if we reach the maximal number of iterations for IRLS without convergence.

These criteria make it necessary to keep track of the negative log-likelihood at each step (to compute the deviance ratio). Hence, each center additionally computes their local log likelihoods, which the server aggregates.

##### Algorithm 1

Federated IRLS Step

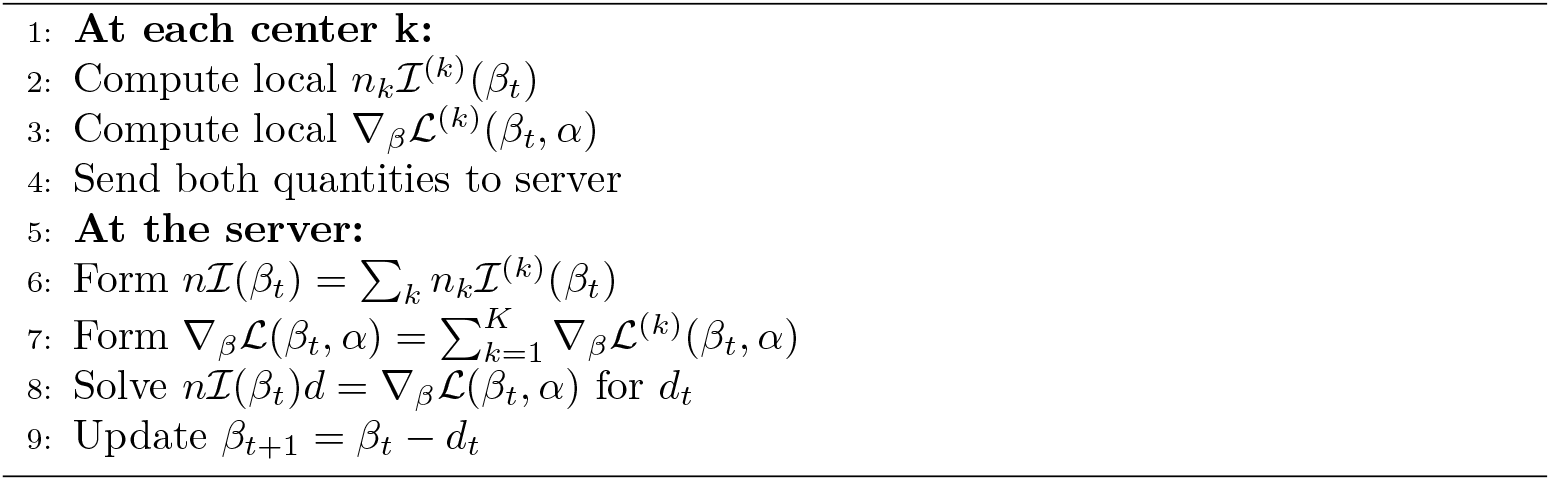

#### The federated Prox Quasi Newton method

When IRLS iterations start diverging, we use instead [40], Algorithm 1, to fit LFCs with the box constraints −*β*_max_ ≤ *β* ≤ *β*_max_. This algorithm consists in iterating the following updates, starting from an initial point *β*_0_:

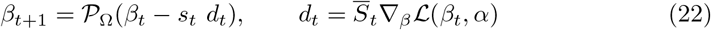

where 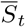 is a modified scaling matrix at time *t, s*_*t*_ is a step size to be fitted with an Armijo line search (see equation 2.5 of [40]), and 𝒫_Ω_ is simply the projection on the box (i.e., clip all coordinates to be inside the box). The modified gradient scaling matrix 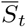 is built from an original gradient scaling matrix *S*_*t*_ (for example, an LBFGS approximation, an inverse Hessian, or an inverse Fisher information matrix) by setting all entries of *S*_*t*_ in a certain index range to 0 (row and column-wise). This index range is completely determined by the original gradient scaling matrix *S*_*t*_, the point *β*_*t*_, and the gradient ∇_*β*_ℒ (*β*_*t*_, *α*) (see equations 2.2 and 2.3 in [40]).

##### Choice of scaling matrix

Since the objective ℒ is nonconvex, using the inverse Hessian as a gradient scaling matrix *S*_*t*_ leads to divergence. By analogy with the original IRLS algorithm, we use the inverse Fisher information matrix as a gradient scaling matrix. This method, known as natural gradient descent, has been shown to have favorable theoretical and experimental properties [41]). Hence, we use the following gradient scaling matrix at step *t*:

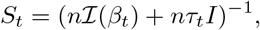

where *τ*_*t*_ is a regularization parameter on a log scale: *τ*_0_ = 1 *τ*_*T*_ = 10^−6^ at the final iteration. The choice of a decreasing regularization term is motivated by the fact that using the inverse Fisher information as a gradient scaling matrix is relevant only close to the optimum. Using large *τ*_*t*_ during the first iterations makes the inverse scaling matrix close to the identity matrix, and the first iterations behave like gradient descent. This scaling matrix can easily be computed in a federated way, exactly as in the IRLS case.

##### Line search

Given a point *β*_*t*_, the gradient at that point ∇_*β*_ *ℒ* (*β*_*t*_, *α*), and an ascent direction *d*_*t*_, the Armijo line search [42] usually works by having a list of exponentially decreasing step sizes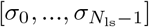, and by iteratively testing the Armijo criterion with *s* = *σ*_*i*_ until it is satisfied:

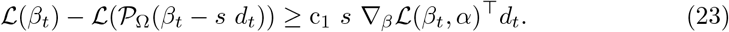

Here, *N*_ls_ is the number of step sizes we are ready to test, the exponential decrease is usually a factor set to 0.5, and c_1_ is a traditionally set constant to 10^−4^. In a pooled (non-federated) setting, such a line search is desirable because we need only evaluate the log-likelihood for each tested step size. However, if we perform this naively in a federated setting, we will pay a communication cost each time we test a step size, since evaluating the log-likelihood requires local communications.

##### Federation

To overcome these limitations, we adopt a greedy approach. Given a direction *d*_*t*_ and a point *β*_*t*_, we ask the centers to compute all Fisher information matrices, gradients, and log-likelihood evaluations for all the potential next points 𝒫_Ω_(*β*_*t*_ − *s d*_*t*_) for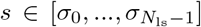. The server can then choose the step size and compute the next point *β*_*t*+1_ and direction *d*_*t*+1_ (see Algorithm 2).The first computation of *d*_0_ is handled separately, and is essentially an IRLS step.

##### A note on privacy

This algorithm shares a large number of quantities from the local centers to the server at each iteration: for each gene, *N*_ls_ gradients, Fisher information matrices and evaluations are shared. Should this constitute an issue, 2 mitigation strategies could be considered. First, all these quantities are actually summed and then used by the server. Hence, a secure aggregation protocol [29] could be applied. Second, we could select the step size first (the centers returning only evaluations of the log-likelihood on the different possible next points) and only compute the necessary quantities to derive the next ascent direction *d*_*t*+1_ in a second step: this would avoid sharing gradients and Fisher information matrices for all possible points, but only when necessary. However, this would double the communication overhead of the algorithm. As communication was our main bottleneck, we chose the version explained in Algorithm 2.

### Detecting and replacing outliers in a federated setting

As seen in the description of the `DESeq2` pipeline, computing Cook’s distances to detect outliers and replacing those outliers with imputed values requires computing trimmed means and variances, i.e. means and variances of counts data from which fixed upper and lower quantiles were removed. Hence, we need to design an algorithm to compute quantiles in a federated setting.

To approximate *r* and 1 − *r* quantiles, we adopt a dichotomy method. Here, we assume the gene and quantile to be fixed: in the code, we perform this dichotomy for all genes and both quantiles in parallel.

#### Algorithm 2

Federated Prox Quasi Newton Step

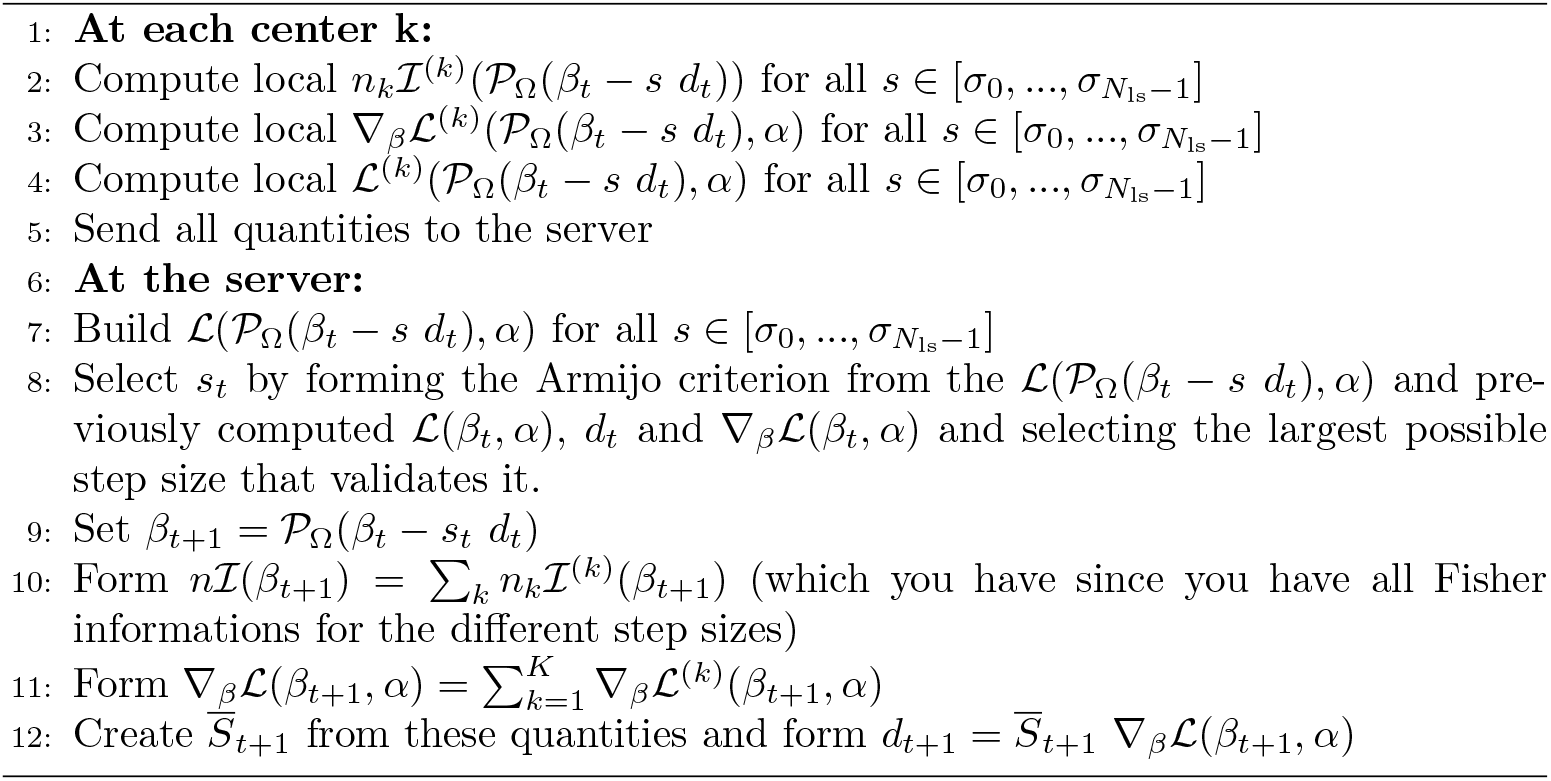

Each center first shares its minimum and maximum normalized counts with the server, which in turn computes the global minimum and maximum. This provides an initial upper and lower bound on the quantile values^2^.

Then, at each iteration, we replace one of the upper and lower bounds by the average of the two, thus dividing the uncertainty in half.

To do so, each center counts the number of samples that is above this mean value. The server then sums these counts and assesses whether the total is above or below the quantile target value (*r* × *n*, where *n* is the total number of samples). If it is above, it replaces the lower bound by the average, otherwise it replaces the upper bound. This procedure is then repeated.

The number of dichotomy iterations thus performed is fixed in advance, with a default number of 40. In particular, this implies that the true quantile value would theoretically be found up to a precision of (max − min)*/*2^40^. Hence, in most cases, machine precision is actually the limiting factor.

### Computing and adjusting p-values in a federated setting

#### Wald tests

For a given gene, Wald tests require two quantities: the LFC coefficient *β*_*g,r*_, and its standard error 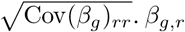 was computed during the IRLS step, and 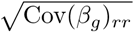 can be derived from *X*^⊤^*WX* (see equation 17), which is computed as the sum of the local (*X* ^(*k*)^)^⊤^ *W*^(*k*)^, 1 ≤ *k* ≤ *K*.

#### P-value Cook filtering

Following the procedure described in the `DESeq2` methodology, we must identify and set the p-value to `NaN` for genes which have:

- (i) at least one sample with a Cook’s distance above the threshold;
- (ii) fewer than 3 samples whose read counts are above the sample with the maximum Cook’s distance;

To achieve (i), centers share whether each gene has remaining Cook outliers, and the server then compiles the total list.

Achieving (ii) requires several steps. First, for each of the genes from (i), we must find the sample with the maximum Cook’s distance, and the corresponding read count value. To do so, centers share (for each gene) the maximum Cook’s distance over their local samples, and the read count of the corresponding sample. The server then derives the global maximum for each gene, and its corresponding counts value threshold, which it shares back to centers.

Second, for each gene, we must count the number of samples above the threshold. This is first performed locally. Centers then share their local numbers, which the server sums.

#### P-value adjustment and independent filtering

P-value adjustment and independent filtering can be performed from p-values and (global) means of normalized counts. As those quantities are computed as part of previous steps, no particular federation effort is required.

### Meta-analysis baselines

Meta-analysis [8] is a set of methods aiming to combine the effects measured in studies with identical or similar designs effect, and derive global effect sizes and variances which can, in turn, be employed to compute a global p-value.

In particular, in the presence of data silos, meta-analysis can be used as a surrogate for federated methods. Indeed, centers need only to share the LFCs and p-values they obtained by running DEA on their local data, and do not have to share any individual RNA transcript or covariate.

We now quickly review the meta-analysis baselines to which we compared the output of `FedPyDESeq2`. Each of the methods described below is first applied independently to each gene, and we then perform the Benjamini-Hochberg procedure [39] to control the false discovery rate.

### P-value combination

P-value combination consists in building a meta-statistic from the set of study (center) p-values, on which a global test is then performed. Note however that these methods only combine p-values, and not the effects (in our case, LFCs). As LFC estimation is a crucial aspect of DEA, we use the average of LFCs weighted by sample size, for lack of a better estimator. P-value combination methods differ in the manner in which the meta-statistic is built. In the following two paragraphs, we describe Stouffer’s and Fisher’s methods (see., e.g., [24] for a description in the context of DEA).

In both cases, we relied on `scipy`’s `combine pvalues` implementation for our experiments.

#### Stouffer’s method

Let 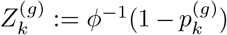 where *ϕ*^−1^ is the inverse cumulative distribution function of the standard normal distribution. Then, under the null hypothesis, it holds

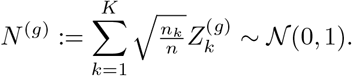

#### Fisher’s method

Under the null hypothesis, it holds

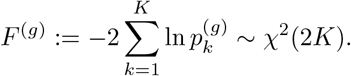

### Fixed-effects model

In a fixed-effects model, we make the assumption that the observed LFCs are distributed around one common true LFC, with a measurement error which is informed entirely by the within-study variance (see, e.g., [43]):

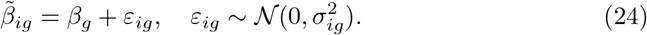

where

- 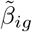is the LFC estimated for gene *g* in *i*-th study, i = 1,…, k;
- 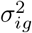is the within-study variance, representing sampling errors conditionally on the *i*-th study;
- *β*_*g*_ is the true global LFC across datasets for gene *g*;

The common LFC is estimated as 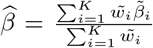, where 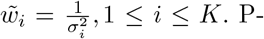 values are then derived thanks to a Wald test, using 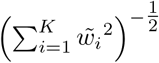 as standard error estimate.

### Random-effects model

Compared with fixed-effects models, random-effects models incorporate a term representing between-study variability (see, e.g., [43]):

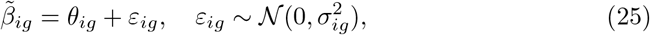

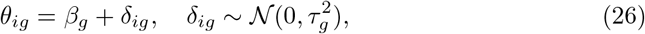

where

- 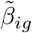 is the LFC estimated for gene g in *i*-th study, i = 1,…, k;
- *θ*_*ig*_ is the true LFC in the *i*-th study;
- 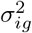 is the within-study variance, representing sampling errors conditionally on the *i*-th study;
- *β*_*g*_ is the true global LFC across datasets for gene *g*;
- *δ*_*ig*_ is the random effect;
- and 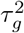 is the between-study variance.

We use 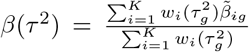, where 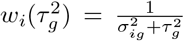, 1 ≤ *i* ≤ *K*, as an estimate of the true LFC *β*_*g*_. P-values are computed thanks to a Wald test, using 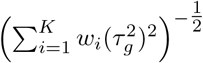 as standard error estimate.

The two random-effects methods that we employ here differ in the manner in which they estimate the between-study variance 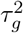. In both cases, we rely on `statsmodels`’s `combine effects` implementation.

#### DerSimonian and Laird (DL) one-step estimator [23]

The DL estimator estimates the between-study variance as

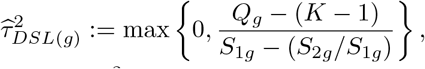

where 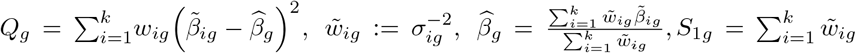 and 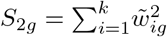.

#### Paule-Mandel iterative estimator [22]

In the Paule-Mandel method, the between-study variance is estimated by iteratively solving for *τ* ^2^ the following equation:

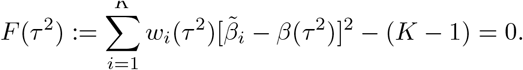

To do so, the following update is applied until convergence:

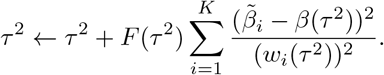

### Largest cohort

We include a final baseline, which consists in using the results output by `PyDESeq2` on the center with the largest cohort. This baseline serves as a sanity check, to confirm that employing federated methods or meta-analysis does bring a benefit compared to individual local results.

### Datasets and preprocessing

Let us now describe the datasets that were used for experiments, and the preprocessing steps they underwent. We used data from The Cancer Genome Atlas (https://www.cancer.gov/tcga) corresponding to 8 different indications:

- TCGA-BRCA: Breast invasive carcinoma
- TCGA-COAD: Colon adenocarcinoma
- TCGA-LUAD: Lung adenocarcinoma
- TCGA-LUSC: Lung squamous cell carcinoma
- TCGA-PAAD: Pancreatic adenocarcinoma
- TCGA-PRAD: Prostate adenocarcinoma
- TCGA-READ: Rectum adenocarcinoma
- TCGA-SKCM: Skin cutaneous melanoma

For each indication, we collected two types of data: (i) bulk RNA-seq raw counts (not restricted to protein-coding genes, for a total of 60k+ genes) and (ii) sample metadata, to be used in the design of our DEA experiments. Both were downloaded from the `recount3` database [44].

#### Read count preprocessing

Counts are indexed by TCGA sample barcode, with columns corresponding to the gene id in ENSEMBL convention. PAR Y genes are filtered out, as they are not common to all samples.

#### Metadata preprocessing

From the metadata, we extract the following factors:

- Patient `gender`;
- Tumor `stage`. Originally from I to IV, we transform `stage` into a binary covariate by grouping stages I-II-III (`Non-advanced`) against IV (`Advanced`);
- `CPE` (for “Consensus measurement of purity estimation”), an aggregate of different purity estimations obtained from [45], taking values between 0 and 1;

We also extract tissue source sites (TSS) from the TCGA barcodes: see “Geographical split” below.

Samples corresponding to normal (non-tumoral) tissues are removed from our experiments. Likewise, any sample with missing design covariate information is dropped from the corresponding experiment (c.f. “Experimental designs” below).

Scripts to obtain the data, including download and preprocessing, are available in the following repository: https://github.com/owkin/fedpydeseq2-datasets.

#### Experimental designs

In the geographical split setting, we perform experiments with three different designs:

- A single-factor design, with tumoral `stage` as sole design factor;
- A multi-factor design with categorical covariates, including tumoral `stage` and `gender` as factors. Because of the gender covariate, TCGA-BRCA and TCGA-PRAD are excluded from this experiment;
- A multi-factor design with a numerical covariate, including tumoral `stage` and `CPE` as factors. TCGA-PAAD is excluded from this experiment, as CPE scores are unavailable.

In the case of the synthetic heterogeneous split, only the single-factor design was used. For all three designs, we tested `Advanced` compared to `Non-Advanced stage` samples, with a LFC=0 null hypothesis.

#### Geographical split

Following the methodology introduced in [17], we simulate a realistic federated setting by separating samples according to the locations they were gathered from. More precisely, we extract TSS from TCGA barcodes (c.f. https://docs.gdc.cancer.gov/Encyclopedia/pages/TCGABarcode/). We then map each TSS code to its source (https://gdc.cancer.gov/resources-tcga-users/tcga-code-tables/tissue-source-site-codes), which we group in 7 regions: Canada, Europe, North-east (USA), Midwest (USA), South (USA), West (USA) and Other. The TSS to region correspondence file is available in the following repository: https://github.com/owkin/fedpydeseq2-datasets. Each region then acts as a federated data center in our experiments.

The corresponding splits are summarized in Tables 2 and 3. The differences between these two tables are due to the fact that we drop samples with no available `CPE` score. Finally, let us note that in some centers, some conditions may have 0 replicates (e.g., they only contain samples with `Non-advanced` tumoral `stages`). In that case, as mentioned in the Discussion, it is not possible to run `PyDESeq2` locally on those centers.

**Table 1.**
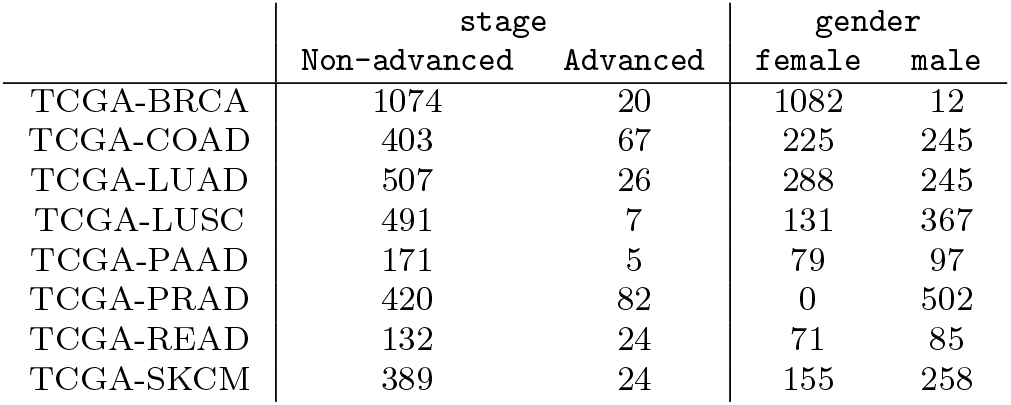
The (stage, gender) multi-factor metadata.

|  | stage |  | gender |  |
| --- | --- | --- | --- | --- |
|  | Non-advanced | Advanced | female | male |
| TCGA-BRCA | 1074 | 20 | 1082 | 12 |
| TCGA-COAD | 403 | 67 | 225 | 245 |
| TCGA-LUAD | 507 | 26 | 288 | 245 |
| TCGA-LUSC | 491 | 7 | 131 | 367 |
| TCGA-PAAD | 171 | 5 | 79 | 97 |
| TCGA-PRAD | 420 | 82 | 0 | 502 |
| TCGA-READ | 132 | 24 | 71 | 85 |
| TCGA-SKCM | 389 | 24 | 155 | 258 |

**Table 2.**
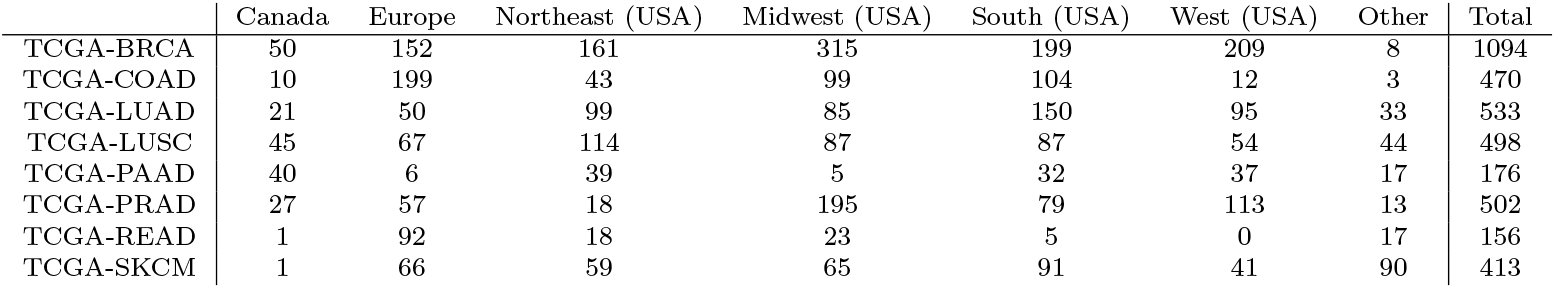
The single-factor and (stage, gender) multi-factor splits.

|  | Canada | Europe | Northeast (USA) | Midwest (USA) | South (USA) | West (USA) | Other | Total |
| --- | --- | --- | --- | --- | --- | --- | --- | --- |
| TCGA-BRCA | 50 | 152 | 161 | 315 | 199 | 209 | 8 | 1094 |
| TCGA-COAD | 10 | 199 | 43 | 99 | 104 | 12 | 3 | 470 |
| TCGA-LUAD | 21 | 50 | 99 | 85 | 150 | 95 | 33 | 533 |
| TCGA-LUSC | 45 | 67 | 114 | 87 | 87 | 54 | 44 | 498 |
| TCGA-PAAD | 40 | 6 | 39 | 5 | 32 | 37 | 17 | 176 |
| TCGA-PRAD | 27 | 57 | 18 | 195 | 79 | 113 | 13 | 502 |
| TCGA-READ | 1 | 92 | 18 | 23 | 5 | 0 | 17 | 156 |
| TCGA-SKCM | 1 | 66 | 59 | 65 | 91 | 41 | 90 | 413 |

**Table 3.** The (stage, CPE) multi-factor split.

|  | Canada | Europe | Northeast (USA) | Midwest (USA) | South (USA) | West (USA) | Other | Total |
| --- | --- | --- | --- | --- | --- | --- | --- | --- |
| TCGA-BRCA | 50 | 152 | 160 | 309 | 191 | 205 | 8 | 1075 |
| TCGA-COAD | 10 | 199 | 43 | 98 | 99 | 12 | 3 | 464 |
| TCGA-LUAD | 21 | 50 | 99 | 83 | 149 | 95 | 33 | 530 |
| TCGA-LUSC | 45 | 67 | 114 | 87 | 87 | 54 | 44 | 498 |
| TCGA-PRAD | 27 | 57 | 18 | 195 | 79 | 108 | 13 | 497 |
| TCGA-READ | 1 | 92 | 18 | 23 | 5 | 0 | 17 | 156 |
| TCGA-SKCM | 1 | 66 | 59 | 63 | 89 | 41 | 90 | 409 |

#### Heterogeneous splits

To better measure the effects of data heterogeneity between centers on our method, we generate two sets of 2-centric synthetic splits: TCGA-CRC, which is based on the union of TCGA-COAD and TCGA-READ, and TCGA-NSCLC, based on TCGA-LUAD and TCGA-LUSC.

Hence, they are not included in meta-analysis baselines. However, `FedPyDESeq2` is still capable to take them into account.

For TCGA-CRC, we create two centers, and assign each TCGA-COAD and TCGA-READ sample to either according a binomial distribution with parameter *h/*2, *h* ∈ {0, 0.25, 0.5, 0.75, 1}. In particular, when *h* = 1, each center receives half of TCGA-COAD and half of TCGA-READ in expectation, whereas when *h* = 0, one of the centers contains all TCGA-COAD samples, and the other all TCGA-READ samples. The same procedure is applied for TCGA-NSCLC. As can be seen in Fig. 4, TCGA-COAD and TCGA-READ (resp. TCGA-LUAD and TCGA-LUSC) have fairly dissimilar sets of DEGs. Hence, we expect that uneven indication splits between two centers results in heterogeneous center data distributions.

**Fig. 4.**
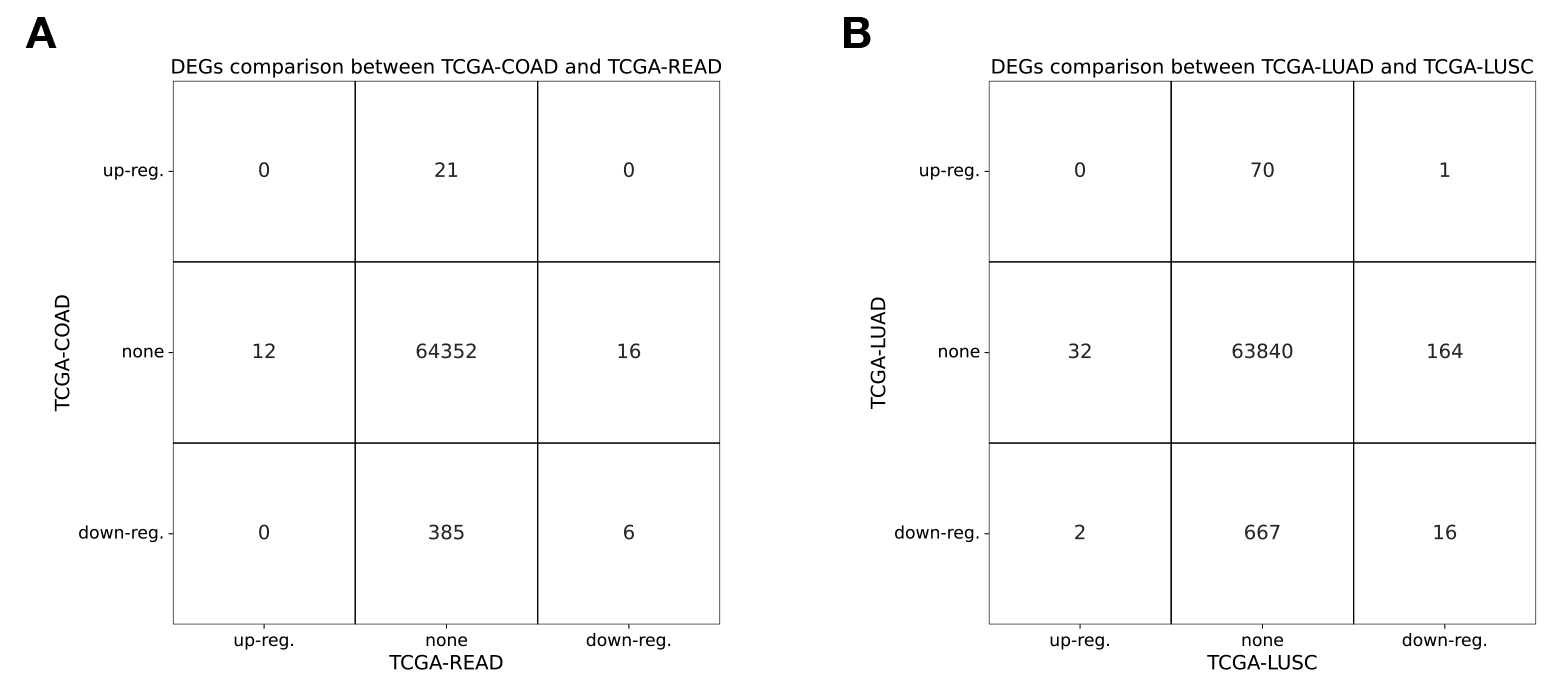
`PyDESeq2` DEGs in the single-factor design. (A) TCGA-COAD vs TCGA-READ, (B) TCGA-LUAD vs TCGA-LUSC

To ensure that our results are reproducible, we fix a random seed (`np.random.seed(42)`). The corresponding splits are summarized in Table 4.

**Table 4.** The heterogeneous single-factor split.

| Heterogeneity level | TCGA-CRC |  |  |  | TCGA-NSCLC |  |  |  |
| --- | --- | --- | --- | --- | --- | --- | --- | --- |
|  | Center 0 |  | Center 1 |  | Center 0 |  | Center 1 |  |
|  | TCGA-COAD | TCGA-READ | TCGA-COAD | TCGA-READ | TCGA-LUAD | TCGA-LUSC | TCGA-LUAD | TCGA-LUSC |
| 0.0 | 470 | 0 | 0 | 156 | 533 | 0 | 0 | 498 |
| 0.25 | 407 | 25 | 63 | 131 | 454 | 58 | 79 | 440 |
| 0.5 | 358 | 43 | 122 | 113 | 387 | 110 | 146 | 388 |
| 0.75 | 287 | 58 | 183 | 98 | 320 | 165 | 213 | 327 |
| 1.0 | 229 | 76 | 241 | 80 | 253 | 261 | 280 | 237 |

### Evaluation

As the goal of `FedPyDESeq2` is to reproduce, from siloed data, the results of `PyDESeq2` on pooled data, we use the output of `PyDESeq2` on pooled data as a ground truth for our benchmarks, and perform evaluations both on `FedPyDESeq2` and our baselines.

#### Comparison of DEGs

For each combination of dataset, split and experimental design, we compare the set of `FedPyDESeq2` and baslines DEGs to those of `PyDESeq2`, setting |LFC| ≥ 2 and padj *<* 0.05 thresholds for differential expression.

The results for `FedPyDESeq2` on the geographical split with the single-factor design are summarized in Fig. 2 A in the form of a confusion matrix, where percentages are expressed w.r.t. `PyDESeq2` DEGs. Results for other experiments are included in the supplementary material.

#### Gene set enrichment analysis

To further validate the output of `FedPyDESeq2`, we perform gene set enrichment analysis (GSEA) on DEGs identified by `FedPyDESeq2` and baselines on siloed RNA-seq, and compare the results to those obtained from pooled `PyDESeq2` DEGs.

We perform GSEA on MSigDB human reactome gene sets [46, 47] using `fgsea` [21] and DEA test statistics as gene ranking criterion.

#### LFC and adjusted p-value relative errors

Finally, we report `FedPyDESeq2` and baseline LFC and adjusted p-value errors relative to pooled `PyDESeq2` LFC and adjusted p-values, that is:

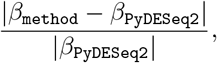

and

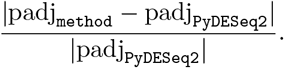

### Deployment and runtime

Our experiments were performed by deploying `Substra` on a set on independent Google Cloud Platform (GCP) instances. For the geographical split, 8 machines were deployed (one for each of the 7 centers, plus one to act as central server). Likewise, 3 machines were used in the synthetic split experiments. Both the centers and the server were allocated `n1-standard-8` GCP instances with 8 virtual processors and 30 Gb of RAM each. With this configuration, each DEA experiment ran in times ranging from 8 to 12 hours.

Note that for demonstration or debugging purposes it is also possible to run `FedPyDESeq2` in simulated mode on a single machine. On a modern laptop, the simulated mode of each experiment runs in about an hour.

## Declarations

## Software availability

The code for `FedPyDESeq2` is available at https://github.com/owkin/fedpydeseq2 under an MIT license. The code to download and preprocess TCGA datasets and to reproduce our experiments is available at https://github.com/owkin/fedpydeseq2-datasets and https://github.com/owkin/fedpydeseq2-paper-experiments.

## Data availability

The data used in this study can be downloaded from The Cancer Genome Atlas (https://www.cancer.gov/tcga) and processed using the following code: https://github.com/owkin/fedpydeseq2-datasets.

## Acknowledgment

The authors would like to thank Sarah Diot-Girard, Guilhem Barthés, Thibault Fouqueray and Thibault Camalon for their invaluable insights and support on how to use `Substra` for federated transcriptomics, Vincent Cabeli for his careful proofreading, and Jade Lum and Davide Mantiero for their help with the FL figure.

The results published here are in whole or part based upon data generated by the TCGA Research Network: https://www.cancer.gov/tcga.

## Footnotes

1 Formally, 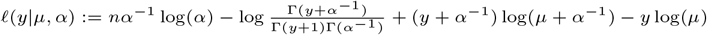.

2 It is also possible to return an upper bound (resp. lower) bound per center which is not the exact maximum (resp. minimum) value, in order to enhance the privacy of the operation, by adding (resp. subtracting) a random non-negative number.

