## Additional Results for "FedPyDESeq2: a federated framework for bulk RNA-seq differential expression analysis"

### A Supplementary results

#### A.1 Supplementary figure: zoom on TCGA-BRCA

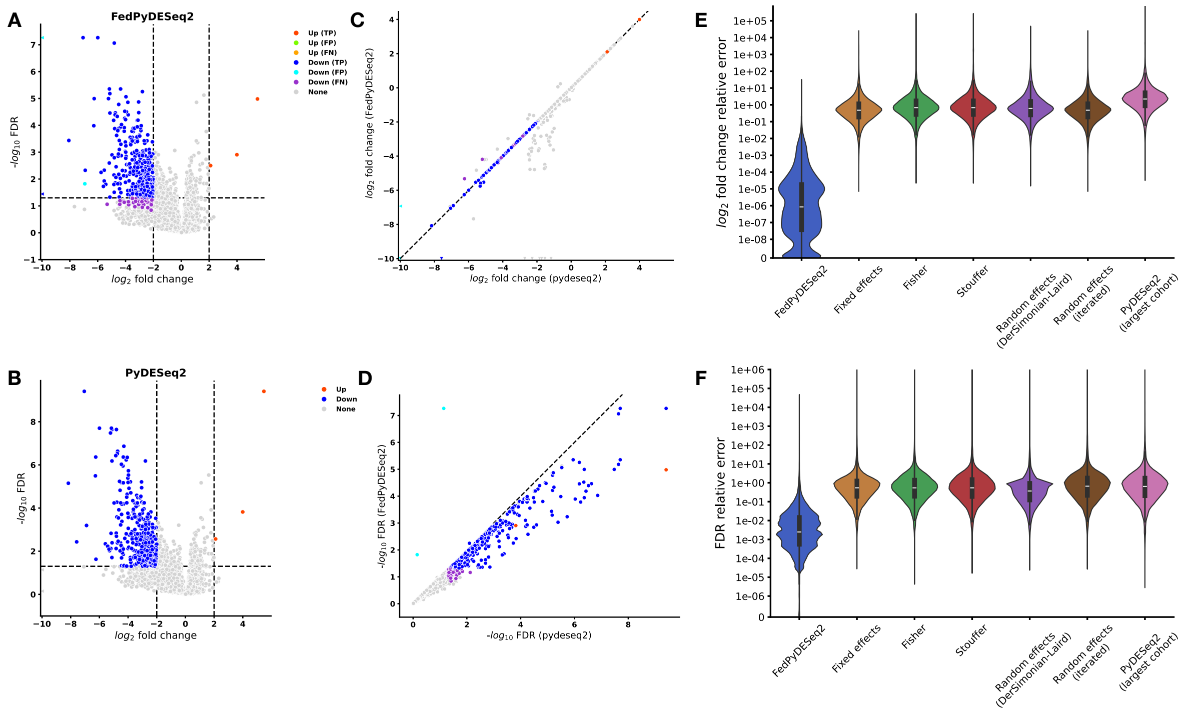


Figure 1: Results of the single-factor design experiment (`Advanced` vs `Non-advanced` tumoral `stage`) with a geographical split on TCGA-BRCA. **(A - B)** Volcano plots of DEGs (with padj $\leq$ 0.05 and $|\text{LFC}| \geq 2$) (top: `FedPyDESeq2`, siloed data; bottom: `PyDESeq2`, pooled data). Differences in DEGs are marked as false positives/negatives (FP/FN) on the `FedPyDESeq2` plot.**(C - D)** LFCs (resp $\log_{10}$ FDR) according to `FedPyDESeq2` against LFCs (resp $\log_{10}$ FDR) according to `PyDESeq2` (from pooled data). **(E - F)** LFC and adjusted p-value relative errors of `FedPyDESeq2` and baseline methods applied on geographically split data, compared with `PyDESeq2` on pooled data ($\frac{|\theta_{\texttt{method}} - \theta_{\texttt{PyDESeq2}}|}{|\theta_{\texttt{PyDESeq2}}|}$). Differences in DEGs are mainly due to `FedPyDESeq2` p-values being more conservative than `PyDESeq2`'s.

#### A.2 Supplementary figure: performance of meta-analysis baselines

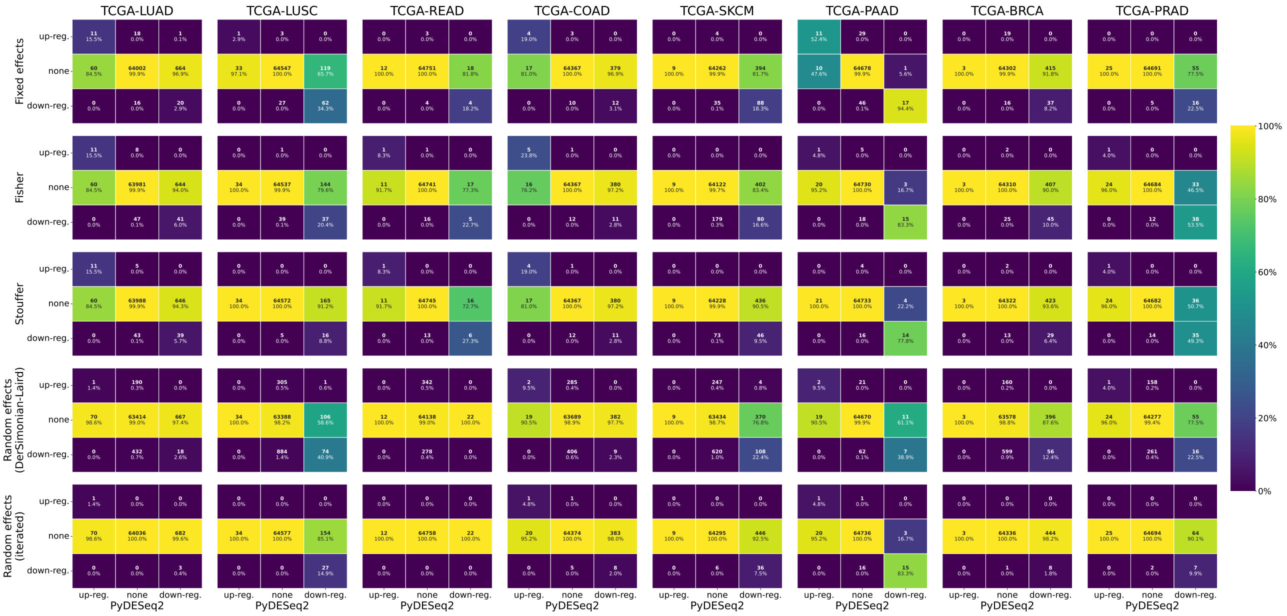


Figure 2: Meta-analysis results of the single-factor design experiment (`Advanced` vs `Non-advanced` tumoral `stage`) with a geographical split. DEGs (with padj $\leq 0.05$ and $|\text{LFC}| \geq 2$) according to baselines (from siloed data) are compared to `PyDESeq2` (from pooled data). Percentages are expressed w.r.t. column totals.

#### A.3 Supplementary figure: multi-factor design

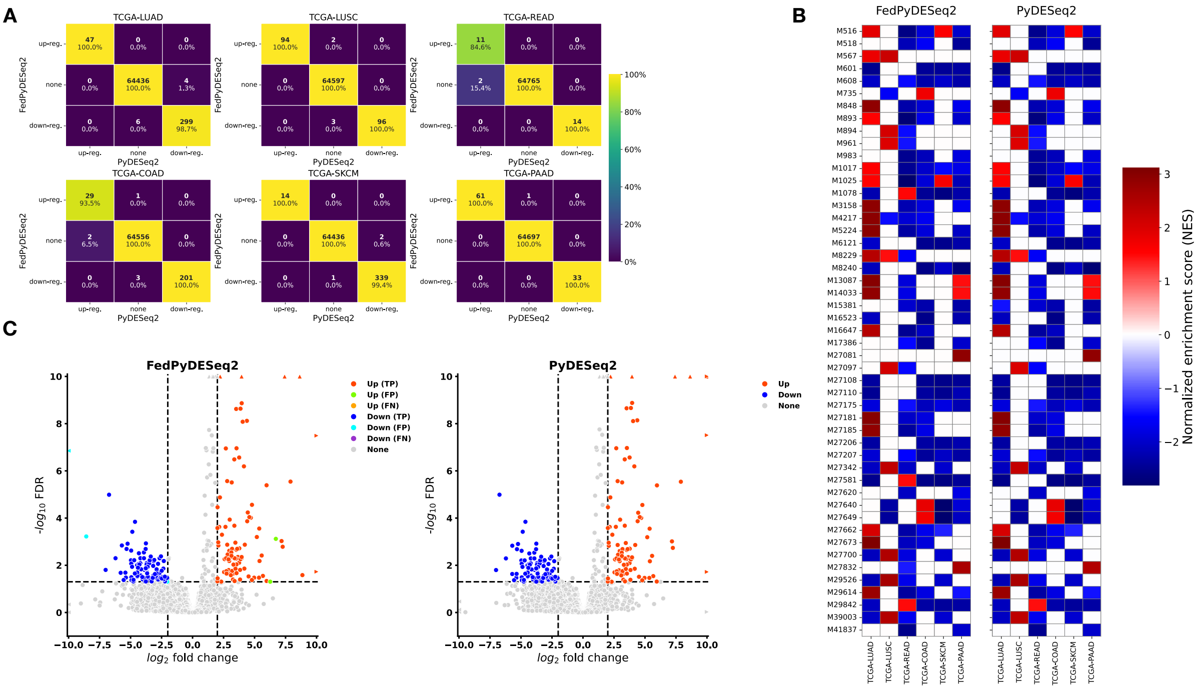


Figure 3: Results of the multi-factor design experiment (design: (`stage`, `gender`), contrast: `Advanced` vs `Non-advanced` tumoral `stage`) with a geographical split. **(A)** DEGs (with padj $\leq$ 0.05 and $|\mathrm{LFC}| \geq 2$) according to `FedPyDESeq2` (from siloed data) and `PyDESeq2` (from pooled data). Percentages are expressed w.r.t. column totals. **(B)** Significantly enriched pathways (padj $\leq$ 0.05) obtained with the `fgsea` package, using Wald statistics as gene-ranking metric. Only pathways that are in the top 10 (according to the adjusted p-value) for at least one indication are represented. If for a given indication, a pathway is not significantly enriched, the corresponding square is left blank. **(C)** Volcano plots of DEGs in TCGA-LUSC (left: `FedPyDESeq2`, siloed data; right: `PyDESeq2`, pooled data). Differences in DEGs are marked as false positives/negatives (FP/FN) on the `FedPyDESeq2` plot.

#### A.4 Supplementary figure: multi-factor design with a numerical covariate

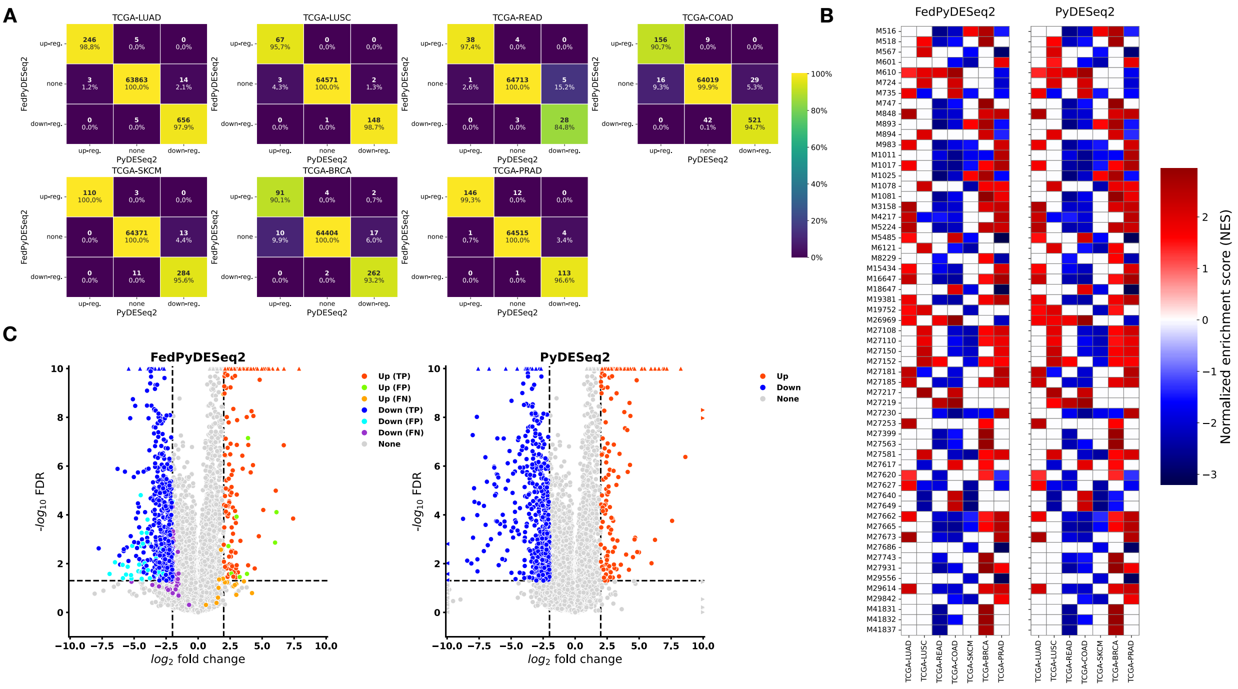


Figure 4: Results of the multi-factor design with a numerical covariate experiment (design: (`stage`, `CPE`), contrast: `Advanced` vs `Non-advanced` tumoral `stage`) with a geographical split. **(A)** DEGs (with padj $\leq 0.05$ and $|\text{LFC}| \geq 2$) according to `FedPyDESeq2` (from siloed data) and `PyDESeq2` (from pooled data). Percentages are expressed w.r.t. column totals. **(B)** Significantly enriched pathways (padj $\leq 0.05$) obtained with the `fgsea` package, using Wald statistics as gene-ranking metric. Only pathways that are in the top 10 (according to the adjusted p-value) for at least one indication are represented. If for a given indication, a pathway is not significantly enriched, the corresponding square is left blank. **(C)** Volcano plots of DEGs in TCGA-COAD (left: `FedPyDESeq2`, siloed data; right: `PyDESeq2`, pooled data). Differences in DEGs are marked as false positives/negatives (FP/FN) on the `FedPyDESeq2` plot.

### B Supplementary results: workflow diagrams and exchanged quantities

In the context of federated learning, which aims to enhance privacy by keeping sensitive data local, transparency regarding data exchange is crucial. We provide comprehensive workflow graphs and detailed tables to explicitly document every quantity shared between the centers and the central server.

The complete workflow graph and its associated table of exchanged quantities can be found in the dedicated supplementary material.

Here, we present two workflow diagrams: one showing the complete workflow with a depth limit of 2 for clarity (meaning that we do not unfold the complete pipeline in terms of server-level and center-level functions, as is done in the dedicated supplementary material), and another focusing on the function computing the log-fold changes, which implements the log-fold change computation through IRLS (Iteratively Reweighted Least Squares) and proximal quasi-Newton optimization.

#### Content of graphs and tables

##### Graphs

In the graphs, white squares represent quantities exchanged between centers and the server, with reference numbers in the table providing detailed descriptions of each exchange. Pink blocks indicate functions executed at the center level, while blue blocks represent operations performed at the server level. Finally, green blocks are functional blocks, which themselves contain a sequence of server-level and center-level functions. The workflow is organized into functional groups, denoted by dashed blocks, which correspond to specific routines implemented in the FedPyDESeq2 package.

##### Tables of exchanged quantities

The tables below present a detailed description of the quantities exchanged during the federated routine. Each row is identified by an ID that corresponds to the numbered white squares in the workflow graph. The table provides the name and type of each exchanged quantity, along with its shape when applicable. The "Description" column offers mathematical definitions and contextual explanations for each quantity.

##### Notations

- $K$: the total number of centers. Centers will be indexed by the letter $k$.
- $p$: the number of parameters in the linear model (number of columns in the design matrix). They will be indexed by the letter $j$.
- $n_k$: the number of observations in center $k$, and $n := \sum_k n_k$ the total number of observations (samples). Samples in any center $k$ will be indexed by the letter $i$.
- $G$ the total number of input genes. Genes will be indexed by the letter $g$. We will also use $G_{\texttt{nz}}$ the number of non-zero genes (computed during the algorithm), which are the genes with at least one sample with non-zero counts, and $G_{\texttt{act}}$, the number of active genes, which will be the number of genes which are still being optimized in a given algorithm.
- $X^{(k)} \in \mathbb{R}^{n_k \times p}$: the design matrix of center $k$. We will denote with $X_{ij}^{(k)}$ the value of the $i$-th row and $j$-th column of the design matrix of center $k$. The local design matrices are built from the input parameters during the first steps of the algorithm.
- $\gamma^{(k)} \in \mathbb{R}^{n_k}$ : the size factors of the samples of center $k$, indexed by samples $i$. $\gamma_i^{(k)}$ denotes the size factor of sample $i$ of center $k$.
- $Y^{(k)} \in \mathbb{R}^{n_k \times G}$ the count matrix of center $k$, indexed by sample number $i$ and gene $g$, s.t. $Y_{ig}^{(k)}$ denotes the count of gene $g$ in sample $i$ of center $k$.
- $Z_{ig}^{(k)} \in \mathbb{R}^{n_k \times G}$ the normalized count matrix indexed by samples $i$ and gene $g$, s.t. $Z_{ig}^{(k)} := Y_{ig}^{(k)} / \gamma_i^{(k)}$ denotes the normalized count of gene $g$ in sample $i$ of center $k$.

#### B.1 Entire workflow (with limited depth)

##### B.1.1 Workflow graph

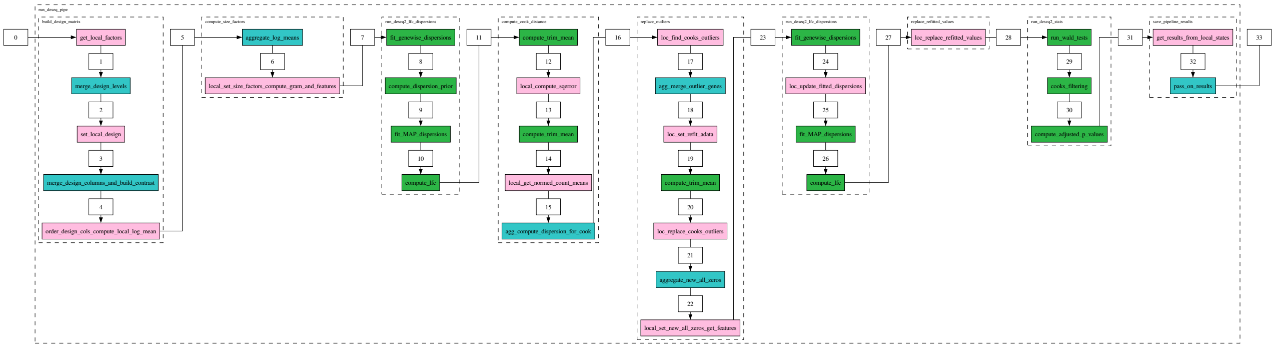


Figure 5: Complete workflow of the FedPyDESeq2 implementation. White squares represent exchanged quantities, pink blocks indicate center-level functions, blue blocks represent server-level operations, green blocks represent other groups of server and center level functions which are not unfolded, and dashed blocks denote functional groups.

##### B.1.2 Table of exchanged quantities

| ID | Name | Type | Shape | Description | Computed by | Sent to |
|---|---|---|---|---|---|---|
| 1 | local_levels | dict | | A dictionary whose keys are the names of the categorical factors, and whose values are the list of values taken by this factor in a given center. For example {stage: [Advanced, Non-advanced], gender: [female]}. | Each center | Server |
| 2 | merged_levels | dict | | A dictionary whose keys are the names of the categorica factors and whose values are arrays containing the list o values taken by this factor across all centers. For example {stage: [Advanced , Non-advanced], gender: [female, male]}. | Server | Center |
| 3 | design_columns | Index | $(p,)$ | The name of the columns of the local design matrix, before aggregation. They are of the form intercept, factor for continuous factors, and factor_level_vs_factor_ref_level otherwise. For example [intercept, stage_Advanced_vs_Non-advanced, gender_male_vs_female] | Each center | Server |
| 4 | merged_columns | Index | $(p,)$ | The union of the design columns across all centers. Local design matrices will then be updated to include al columns. | Server | Center |
| | contrast | list | | A list of three strings representing the contrast o interest, in case it is not specified by the user Of the form [factor, level1, level2]. For example [stage, Advanced, Non-advanced]. | Server | Center |
| 5 | log_mean | nparray | $(G,)$ | For each gene, the mean of the log of the counts across all samples in a center $\overline{\log(Y)}_g^{(k)} = \frac{1}{n_k}\sum_{i=1}^{n_k}\log(Y_{ig}^{(k)})$ | Each center | Server |
| | n_samples | int | | The number of samples in a center $n_k$ for each center $k$. | Each center | Server |
| 6 | global_log_mean | nparray | $(G,)$ | The mean of the log of the counts across all samples in all centers $\overline{\log(Y)}_g = \sum_{k=1}^{K}\frac{n_k}{n}\overline{\log(Y)}_g^{(k)}$. | Server | Center |
| 7 | local_features | nparray | $(p, G)$ | $\Phi^{(k)} := X^{(k)\top}Z^{(k)}$. | Each center | Server |
| | local_gram_matrix | nparray | $(p, p)$ | The gram matrix of the local design matrix $G^{(k)} := (X^{(k)})^\top X^{(k)}$. | Each center | Server |
| 8 | genewise_dispersions | nparray | $(G,)$ | For each gene $g$, the current estimate of the dispersion parameter $\alpha_g$. This estimate is computed by first computing the global nll (summing all local nlls) as well as the global Cox-Reid regularization term, which is half the log determinant of the sum of the local Cox-Reid matrices. The Cox-Reid regularized nll per gene and per dispersion in the grid is obtained by summing the regularization term and the nll. Finally, for every gene, the dispersion parameter is estimated by taking the minizer of this regularized nll on the grid of size $N_{\mathrm{gs}}$. | Server | Center |
| | lower_log_bounds | nparray | $(G,)$ | For each gene $g$, the maximum of the log of the min dispersion and $\alpha_g - \delta$ where $\delta$ is the current mesh size of the grid and $\alpha_g$ is the current dispersion estimate. This value will be used as a lower bound for the next grid search fo the dispersion parameter. | Server | Center |
| | upper_log_bounds | nparray | $(G,)$ | For each gene $g$, the minimum of the log of the max dispersion and $\alpha_g + \delta$ where $\delta$ is the current mesh size o the grid and $\alpha_g$ is the current dispersion estimate. This value will be used as an upper bound for the next grid search for the dispersion parameter. | Server | Center |
| 9 | disp_function_type | str | | The type of the dispersion function used to model the dispersions. It can be either "parametric", if the iterative scheme to fit the trend curve has converged, the LBFGS-B method used to fit the parameters has converged, and the coefficienst of the trend curve are non-negative; o "mean" otherwise. | Server | Center |
| | prior_disp_var | float | () | A prior on the variance of the log dispersions around the log trend curve $\sigma^2_{\mathtt{trend}}$, estimated as the maximum between 0.25 and _squared_log_res $- \psi_1((n-p)/2)$, where $\psi_1(f/2)$ is the variance of the log of a $\chi^2_f$ distribution For more details, see [2]. | Server | Center |

| | | | | | | |
|---|---|---|---|---|---|---|
| | _squared_logres | float | () | The squared mean absolute deviation of the differenc between the log of the genewise dispersions and the log of the fitted dispersions, restricted to the non-zero genes whose gene-wise dispersions are above $100 \times$ min_disp. The mean absolute deviation estimate is defined as the median of the absolute difference between the log residua and its median, scaled by the percent point function o the normal distribution at 0.75. | Server | Center |
| | trend_coeffs | nparray | (2, ) | The coefficients of the trend curve fitted to the dispersions. We model the dispersions $\alpha_g$ of the gene $g$ as a sample from an exponential distribution whose mean $\alpha_{\mathtt{trend}}(\overline{Z}_g)$ is a function of $\overline{Z}_g$ parametrized by two coefficients $\alpha_0$ and $\mathrm{a}_1$; $\alpha_{\mathtt{trend}}(\overline{Z}_g) = \alpha_0 + \frac{\mathrm{a}_1}{\overline{Z}_g}$. $\alpha_{\mathtt{trend}}$ is called the trend curve. The coefficients are $(\alpha_0, \mathrm{a}_1)$ and are obtained by starting with the set of non zero genes and by iteratively i) minimizing the negative log likelihood of the exponential distribution on the set o genes with LBFGS-B and ii) removing the genes where the ratio between the dispersion $\alpha_g$ and the trend curve $\alpha_{\mathtt{trend}}(\overline{Z}_g)$ is above 15 or below $10^{-4}$ (we consider these genes as outliers to this model), until the set of genes is stable (see [2] for more details). | Server | Center |
| | fitted_dispersions | nparray | $(G, )$ | For each gene $g$ with zero counts across all centers nan. For each non-zero gene $g$, either $\alpha_{\mathtt{trend}}(\overline{Z}_g)$ if the disp_function_type is "parametric" (i.e., the fitting of the parameters has converged), or mean_disp otherwise (i.e., i the disp_function_type is "mean"). Denoted with $\alpha_g^{\mathtt{trend}}$. | Server | Center |
| | mean_disp | NoneType | | None if the dispersion function type is "parametric" and the trimmed mean of the genewise dispersions whose value is above $10 \times$ min_disp, with trimming proportion 0.001 otherwise. | Server | Center |
| 12 | trimmed_mean_normed_counts | DataFrame | $(G, \lvert\mathcal{L}_{\geq 3}\rvert)$ | For each gene $g$ and each level $l \in \mathcal{L}_{\geq 3}$, the corresponding entry $\overline{Z}^{\mathtt{trim}}_{g,l}$ is the approximation of the trimmed mean of the normed counts for gene $g$ and samples whose line in the design corresponds to level $l$ with trim ratio $r_{\mathtt{trim}}$ computed by summing the trimmed_local_sums and dividing by the sum of the local n_samples for the corresponding gene and level. | Server | Center |
| 14 | varEst | nparray | $(G, )$ | For each gene $g$, the trimmed variance estimate of the normed counts, denoted with $V_g^{\mathtt{trim}}$. If use_lvl is True, for each gene $g$, it is the maximum across all admissible levels of the trimmed mean of the squared error for the gene and level in question, with trim ratio $r_{\mathtt{trim}}$, scaled (multiplied) by the scale factor 1.51. If use_level is False, the trimmed mean of the squared error scaled by 1.51 with trim ratio $r_{\mathtt{trim}}$. | Server | Center |
| 15 | local_hat_matrix | nparray | $(G_{\mathtt{nz}}, p, p)$ | For each gene $g$, the hat matrix $H_g^{(k)} = (X^{(k)})^\top W_g^{(k)} X^{(k)}$ where $W_g^{(k)} \in \mathbb{R}^{n_k \times n_k}$ is the diagonal matrix with diagonal entries $\frac{\mu_{ig}^{(k)}}{1+\mu_{ig}^{(k)}\alpha_g}$ for $1 \leq i \leq n_k$. $\alpha_g$ is the dispersion estimate of the gene and $\mu_{ig}^{(k)}$ is the expected value of the gene for sample $i$ for parameter $\beta$, that is $\gamma_g^{(k)} \exp(X_i^{(k)} \cdot \beta_g)$. | Each center | Server |
| | mean_normed_counts | nparray | $(G, )$ | For each gene, the mean of the local normed counts, i.e. $\overline{Z}_g^{(k)} = \frac{1}{n_k} \sum_{i=1}^{n_k} Z_{ig}^{(k)}$. | Each center | Server |
| | n_samples | int | | The number of samples in a center $n_k$ for each center $k$. | Each center | Server |
| | varEst | nparray | $(G, )$ | For each gene $g$, the trimmed variance estimate of the normed counts, denoted with $V_g^{\mathtt{trim}}$. This quantitiy is passed on from the previous shared state. | Each center | Server |
| | _skip_cooks | bool | | A boolean indicating whether to skip the computation o the intermediate quantities to compute the of the Cook's distance. This is set to True if the Cook's distance is stored in the local state (which is not the case by default due to memory issues). Otherwise, it is set to False (default behaviour). | Each center | Server |
| 16 | cooks_dispersions | nparray | $(G, )$ | For each gene $g$, a robust estimate of the dispersion parameter $\alpha_g^{\mathtt{cooks}}$ computed from the trimmed variance estimate and the global mean of the normed counts. We compute this estimate as $\max((V_g^{\mathtt{trim}} - \overline{Y}_g)/\overline{Y}_g^2, 0.04)$. | Server | Center |
| | global_hat_matrix_inv | nparray | $(G_{\mathtt{nz}}, p, p)$ | For each gene $g$, we compute the global hat matrix as the sum of the local hat matrices, and its inverse. | Server | Center |
| 17 | replaceable_samples | bool | () | A boolean indicating if there are any replaceable samples in the center. A sample $i$ of center $k$ is said to be replaceable if there are at least min_replicates samples across all centers which share the same design factor levels as $i$. min_replicates is a user defined parameter, set to 7 by default. | Each center | Server |
| | local_genes_to_replace | set | | The set of genes $g$ for which the Cook's distance is above the cutoff value for any sample in the center. For a given gene $g$ and given sample $i$, the Cook's distance is computed as $\frac{h_{ig}^{(k)}}{p(1-h_{ig}^{(k)})^2} (R^2)_{ig}^{(k)}$ where $(R^2)_{ig}^{(k)}$ is the squared Pearson residual of the negative binomial GLM with log fold changes $\beta_g$ and dispersions $\alpha_g^{\mathtt{cooks}}$ computed as $(Y_{ig}^{(k)} - \mu_{ig}^{(k)})^2/(V^{\mathtt{NB}})_{ig}^{(k)}$, and $h_{ig}^{(k)}$ is the $i$-th diagonal element of $X^{(k)} H_g^{-1} (X^{(k)})^\top$, where $H_g^{-1}$ is the inverse of the global hat matrix. The cutoff value is set to the 0.99-th quantile of the F-distribution with $p$ and $n-p$ degrees of freedom. Here $\mu_{ig}^{(k)} = \gamma_i^{(k)} \exp(X_i^{(k)} \cdot \beta_g)$ and $(V^{\mathtt{NB}})_{ig}^{(k)} = \mu_{ig}^{(k)}(1 + \mu_{ig}^{(k)} \alpha_g^{\mathtt{cooks}})$. | Each center | Server |
| 18 | genes_to_replace | set | | The set of genes $g$ for which the Cook's distance is above the cutoff value for any sample in any center (the union of the local genes to replace across all centers). | Server | Center |
| 20 | trimmed_mean_normed_counts | nparray | $(G_{\mathtt{r}}, )$ | For each gene to replace $g$, the trimmed mean of the normed counts across all samples, denoted with $\overline{Z}_g^{\mathtt{trim}}$ with trim ratio set to 0.2. | Server | Center |

| | | | | | | |
|---|---|---|---|---|---|---|
| 21 | loc_new_all_zeroes | nparray | $(G_{\mathtt{r}},)$ | A boolean array which for each gene to replace $g$ indicates if the new count matrix of the center is all zeroes (across samples) for this gene (the new count matrix is computed by imputing the Cook's outliers with the trimmed mean of the normed counts times the size factor). | Each center | Server |
| 22 | new_all_zeroes | nparray | $(G_{\mathtt{r}},)$ | A boolean array which for each gene to replace $g$ indicates if all counts across all centers are zero for this gene. | Server | Center |
| 23 | local_features | nparray | $(p, G_{\mathtt{act}})$ | $\Phi^{(k)} := (X^{(k)})^\top Z^{(k)}$, where $Z^{(k)}$ is the normalized counts in the center $k$ on the set of genes to replace, where the value of Cook's outliers have been replaced using a trimmed mean. | Each center | Server |
| 24 | genewise_dispersions | nparray | $(G_{\mathtt{r}},)$ | For each gene $g$, the current estimate of the dispersion parameter $\alpha_g$. This estimate is computed by first computing the global nll (summing all local nlls) as well as the global Cox-Reid regularization term, which is half the log determinant of the sum of the local Cox-Reid matrices. The Cox-Reid regularized nll per gene and per dispersion in the grid is obtained by summing the regularization term and the nll. Finally, for every gene, the dispersion parameter is estimated by taking the minizer of this regularized nll on the grid of size $N_{\mathtt{gs}}$. | Server | Center |
| | lower_log_bounds | nparray | $(G_{\mathtt{r}},)$ | For each gene $g$, the maximum of the log of the min dispersion and $\alpha_g - \delta$ where $\delta$ is the current mesh size of the grid and $\alpha_g$ is the current dispersion estimate. This value will be used as a lower bound for the next grid search for the dispersion parameter. | Server | Center |
| | upper_log_bounds | nparray | $(G_{\mathtt{r}},)$ | For each gene $g$, the minimum of the log of the max dispersion and $\alpha_g + \delta$ where $\delta$ is the current mesh size of the grid and $\alpha_g$ is the current dispersion estimate. This value will be used as an upper bound for the next grid search for the dispersion parameter. | Server | Center |
| 29 | global_hat_matrix_inv | nparray | $(G_{\mathtt{nz}}, p, p)$ | For each gene $g$, we compute the global hat matrix as the sum of the local hat matrices, and its inverse. | Server | Center |
| | cooks_dispersions | nparray | $(G,)$ | For each gene $g$, a robust estimate of the dispersion parameter $\alpha_g^{\mathtt{cooks}}$ computed from the trimmed variance estimate and the global mean of the normed counts. We compute this estimate as $\max((V_g^{\mathtt{trim}} - \overline{Y}_g)/\overline{Y}_g^2, 0.04)$. | Server | Center |
| | p_values | nparray | $(G,)$ | For each gene, the p-value of the Wald statistic, computed from the survival function of the normal distribution applied to the wald statistic. | Server | Center |
| | wald_statistics | nparray | $(G,)$ | For each gene, the Wald statistic of the gene expression. This statistics depends on the `lfc_null` parameter which sets the null hypothesis on the log fold change (set to 0 by default), and the `alt_hypothesis` parameter, which defines the alternative hypothesis on the log fold change (set to `None` by default, can be `greater`, `greaterAbs`, `less`, `lessAbs`). If the alternative hypothesis is `None`, then the Wald statistic is computed as the centered normalized log fold change. | Server | Center |
| | wald_se | nparray | $(G,)$ | For each gene, the standard error on the log fold change value for the given contrast given by the GLM. | Server | Center |
| 30 | p_values | nparray | $(G,)$ | If Cook's filtering is enabled (which is the case by default with the `cooks_filter` parameter), then the p-values of genes which have $\leq 2$ samples above the gene count of the sample maximizing the Cook's distance across all centers are set to `nan`. Otherwise, passed on without modification. | Server | Center |
| | wald_statistics | nparray | $(G,)$ | For each gene, the Wald statistic of the gene expression. Passed on without modification. | Server | Center |
| | wald_se | nparray | $(G,)$ | For each gene, the standard error on the log fold change value for the given contrast given by the GLM. Passed on without modification. | Server | Center |
| 32 | contrast | list | | A list of three strings representing the contrast of interest, in case it is not specified by the user. Of the form `[factor, level1, level2]`. For example `[stage, Advanced, Non-advanced]`. | Each center | Server |
| | _squared_logres | float | () | The squared mean absolute deviation of the difference between the log of the genewise dispersions and the log of the fitted dispersions, restricted to the non-zero genes whose gene-wise dispersions are above $100 \times$ `min_disp`. The mean absolute deviation estimate is defined as the median of the absolute difference between the log residual and its median, scaled by the percent point function of the normal distribution at 0.75. | Each center | Server |
| | refitted | nparray | $(G,)$ | A boolean array marking genes which can be replaced and which, after replacing the count value by the imputation value, are non-zero. | Each center | Server |
| | replaced | nparray | $(G,)$ | A boolean array marking genes $g$ for which the Cook's distance is above the cutoff value for any sample in any center (the union of the local genes to replace across all centers). | Each center | Server |
| | wald_se | nparray | $(G,)$ | For each gene, the standard error on the log fold change value for the given contrast given by the GLM. | Each center | Server |
| | wald_statistics | nparray | $(G,)$ | For each gene, the Wald statistic of the gene expression. | Each center | Server |
| | p_values | nparray | $(G,)$ | For each gene, the p-value of the Wald statistic, computed from the survival function of the normal distribution applied to the wald statistic, and set to `nan` if the gene is a Cook's outlier if `cooks_filter` is enabled. | Each center | Server |
| | prior_disp_var | float | () | A prior on the variance of the log dispersions around the log trend curve $\sigma^2_{\mathtt{trend}}$, estimated as the maximum between 0.25 and `_squared_log_res` $- \psi_1((n-p)/2)$, where $\psi_1(f/2)$ is the variance of the log of a $\chi^2_f$ distribution. For more details, see [2]. | Each center | Server |
| | LFC | DataFrame | $(G, p)$ | The log fold changes of the gene expression. This dataframe is indexed by genes on one hand, and by design column names on the other. | Each center | Server |
| | fitted_dispersions | nparray | $(G,)$ | For each gene $g$ with zero counts across all centers, `nan`. For each non-zero gene $g$, either $\alpha_{\mathtt{trend}}(\overline{Z}_g)$ if the disp_function_type is "parametric" (i.e., the fitting of the parameters has converged), or `mean_disp` otherwise (i.e., if the disp_function_type is "mean"). Denoted with $\alpha_g^{\mathtt{trend}}$. | Each center | Server |
| | non_zero | nparray | $(G,)$ | A boolean array indicating which genes have non-zero counts in at least one center. | Each center | Server |
| | genewise_dispersions | nparray | $(G,)$ | For each gene $g$, the current estimate of the dispersion parameter $\alpha_g$. This estimate is computed by first computing the global nll (summing all local nlls) as well as the global Cox-Reid regularization term, which is half the log determinant of the sum of the local Cox-Reid matrices. The Cox-Reid regularized nll per gene and per dispersion in the grid is obtained by summing the regularization term and the nll. Finally, for every gene, the dispersion parameter is estimated by taking the minizer of this regularized nll on the grid of size $N_{\mathtt{gs}}$. | Each center | Server |

| | | | | | | |
|---|---|---|---|---|---|---|
| | dispersions | nparray | $(G, )$ | The estimated dispersions for each gene, which are the MAP dispersions if the gene is not an outlier w.r.t. the trend curve, and the gene-wise dispersions otherwise. | Each center | Server |
| | MAP_dispersions | nparray | $(G, )$ | For each gene $g$, the current estimate of the MAP dispersion parameter $\alpha_g$. This estimate is computed by first computing the global nll (summing all local nlls) as well as the global Cox-Reid regularization term, which is half the log determinant of the sum of the local Cox-Reid matrices, and the regularization coming from the prior on the log dispersions around the trend curve. The Cox-Reid, prior regularized nll per gene and per dispersion in the grid is obtained by summing the regularization terms and the nll. Finally, for every gene, the MAP dispersion parameter is estimated by taking the minizer of this regularized nll on the grid of size $N_{\texttt{gs}}$. | Each center | Server |
| | gene_names | Index | $(G, )$ | The gene names. | Each center | Server |
| | padj | Series | $(G, )$ | The adjusted p-values for each gene, computed from the p-values using independent filtering if the `independent_filter` parameter is `True` (default), and the Benjamini-Hochberg procedure otherwise. | Each center | Server |
| 33 | prior_disp_var | float | $()$ | A prior on the variance of the log dispersions around the log trend curve $\sigma^2_{\texttt{trend}}$, estimated as the maximum between 0.25 and `_squared_log_res` $- \psi_1((n-p)/2)$, where $\psi_1(f/2)$ is the variance of the log of a $\chi^2_f$ distribution. For more details, see [2]. | Server | All |
| | refitted | nparray | $(G, )$ | A boolean array marking genes which can be replaced and which, after replacing the count value by the imputation value, are non-zero. | Server | All |
| | replaced | nparray | $(G, )$ | A boolean array marking genes $g$ for which the Cook's distance is above the cutoff value for any sample in any center (the union of the local genes to replace across all centers). | Server | All |
| | wald_se | nparray | $(G, )$ | For each gene, the standard error on the log fold change value for the given contrast given by the GLM. | Server | All |
| | wald_statistics | nparray | $(G, )$ | For each gene, the Wald statistic of the gene expression. | Server | All |
| | p_values | nparray | $(G, )$ | For each gene, the p-value of the Wald statistic, computed from the survival function of the normal distribution applied to the wald statistic, and set to `nan` if the gene is a Cook's outlier if `cooks_filter` is enabled. | Server | All |
| | padj | Series | $(G, )$ | The adjusted p-values for each gene, computed from the p-values using independent filtering if the `independent_filter` parameter is `True` (default), and the Benjamini-Hochberg procedure otherwise. | Server | All |
| | genewise_dispersions | nparray | $(G, )$ | For each gene $g$, the current estimate of the dispersion parameter $\alpha_g$. This estimate is computed by first computing the global nll (summing all local nlls) as well as the global Cox-Reid regularization term, which is half the log determinant of the sum of the local Cox-Reid matrices. The Cox-Reid regularized nll per gene and per dispersion in the grid is obtained by summing the regularization term and the nll. Finally, for every gene, the dispersion parameter is estimated by taking the minizer of this regularized nll on the grid of size $N_{\texttt{gs}}$. | Server | All |
| | fitted_dispersions | nparray | $(G, )$ | For each gene $g$ with zero counts across all centers, `nan`. For each non-zero gene $g$, either $\alpha_{\texttt{trend}}(\overline{Z}_g)$ if the disp_function_type is "parametric" (i.e., the fitting of the parameters has converged), or `mean_disp` otherwise (i.e., if the disp_function_type is "mean"). Denoted with $\alpha_g^{\texttt{trend}}$. | Server | All |
| | non_zero | nparray | $(G, )$ | A boolean array indicating which genes have non-zero counts in at least one center. | Server | All |
| | dispersions | nparray | $(G, )$ | The estimated dispersions for each gene, which are the MAP dispersions if the gene is not an outlier w.r.t. the trend curve, and the gene-wise dispersions otherwise. | Server | All |
| | MAP_dispersions | nparray | $(G, )$ | For each gene $g$, the current estimate of the MAP dispersion parameter $\alpha_g$. This estimate is computed by first computing the global nll (summing all local nlls) as well as the global Cox-Reid regularization term, which is half the log determinant of the sum of the local Cox-Reid matrices, and the regularization coming from the prior on the log dispersions around the trend curve. The Cox-Reid, prior regularized nll per gene and per dispersion in the grid is obtained by summing the regularization terms and the nll. Finally, for every gene, the MAP dispersion parameter is estimated by taking the minizer of this regularized nll on the grid of size $N_{\texttt{gs}}$. | Server | All |
| | gene_names | Index | $(G, )$ | The gene names. | Server | All |
| | _squared_logres | float | $()$ | The squared mean absolute deviation of the difference between the log of the genewise dispersions and the log of the fitted dispersions, restricted to the non-zero genes whose gene-wise dispersions are above $100 \times$ `min_disp`. The mean absolute deviation estimate is defined as the median of the absolute difference between the log residual and its median, scaled by the percent point function of the normal distribution at 0.75. | Server | All |
| | LFC | DataFrame | $(G, p)$ | The log fold changes of the gene expression. This dataframe is indexed by genes on one hand, and by design column names on the other. | Server | All |
| | contrast | list | | A list of three strings representing the contrast of interest, in case it is not specified by the user. Of the form `[factor, level1, level2]`. For example `[stage, Advanced, Non-advanced]`. | Server | All |

#### B.2 Focus on the computing of log-fold changes (LFCs)

##### B.2.1 Workflow graph

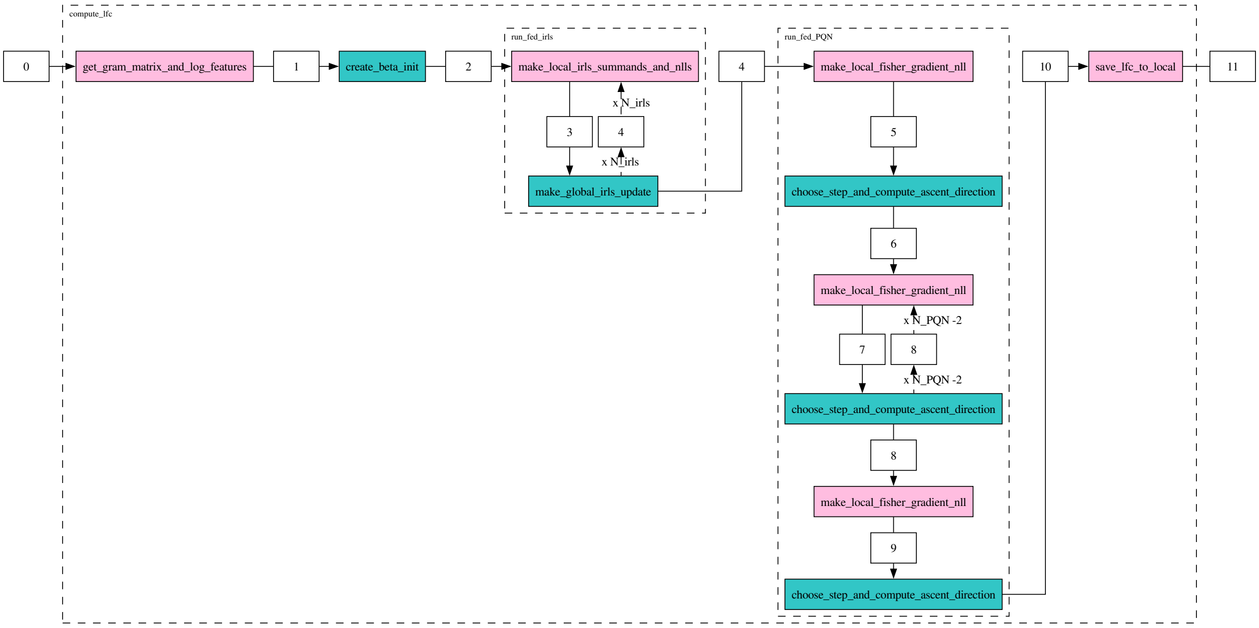


Figure 6: Workflow which is used to compute the log-fold changes using IRLS and PQN. White squares represent exchanged quantities, pink blocks indicate center-level functions, blue blocks represent server-level operations, and dashed blocks denote functional groups.

##### B.2.2 Table of exchanged quantities

| ID | Name | Type | Shape | Description | Computed by | Sent to |
|---|---|---|---|---|---|---|
| 1 | gram_full_rank | bool | () | A boolean indicating whether the global Gramm matrix $G$ is full rank. | Each center | Server |
| | n_non_zero_genes | int | | The number of genes with non-zero counts in at least one center. | Each center | Server |
| | local_log_features | nparray | $(G_{\texttt{nz}}, p)$ | $\Phi_k = (X^{(k)})^\top \log(Z^{(k)} + 0.1)$. | Each center | Server |
| | global_gram_matrix | nparray | $(p, p)$ | The sum of the local Gramm matrix across all centers $G = \sum_k G^{(k)}$. | Each center | Server |
| 2 | beta | nparray | $(G_{\texttt{nz}}, p)$ | The initial value of the $\beta$ parameter. If the Gramm matrix is full rank, is set to $\beta_g = G^{-1}(\sum_k \Phi_g^{(k)})$ for all genes $g$. Otherwise, it is set as a weighed average of the normed log means, i.e., $\beta_g = \sum_k \frac{n_k}{n} \overline{\log(Z)}_g^{(k)}$, where $\overline{\log(Z)}_g^{(k)} = \frac{1}{n_k} \sum_{i=1}^{n_k} \log(Z_{ig}^{(k)})$ is computed locally. | Server | Center |
| | irls_diverged_mask | nparray | $(G_{\texttt{nz}},)$ | A boolean array indicating which non zero genes have caused the IRLS algorithm to diverge. Initialized to `False` | Server | Center |
| | irls_mask | nparray | $(G_{\texttt{nz}},)$ | A boolean array indicating which non zero genes are currently being optimized by the IRLS algorithm. Initialized to `True`. | Server | Center |
| | global_nll | nparray | $(G_{\texttt{nz}},)$ | For each gene $g$, the negative log likelihood of the negative binomial GLM, initialized at 1000 for all genes. | Server | Center |
| | round_number_irls | int | | The current round number of the IRLS algorithm. Initialized at 0. | Server | Center |
| 3 | round_number_irls | int | | The current round number of the IRLS algorithm. Simply passed on. | Each center | Server |
| | global_nll | nparray | $(G_{\texttt{nz}},)$ | For each gene $g$, the negative log likelihood at the $\beta_g$ parameter. Simply passed on. | Each center | Server |
| | irls_mask | nparray | $(G_{\texttt{nz}},)$ | A boolean array indicating which non zero genes are currently being optimized by the IRLS algorithm. Simply passed on. | Each center | Server |
| | irls_diverged_mask | nparray | $(G_{\texttt{nz}},)$ | A boolean array indicating which non zero genes have caused the IRLS algorithm to diverge. Simply passed on | Each center | Server |
| | beta | nparray | $(G_{\texttt{nz}}, p)$ | The log fold change $\beta$ parameter. Not modified. | Each center | Server |
| | local_features | nparray | $(G_{\texttt{act}}, p)$ | For each active gene $g$, the projected features $(X^{(k)})^\top W_g^{(k)} \phi_g^{(k)}$, where $\phi_{ig}^{(k)} = \log\left(\frac{\mu_{ig}^{(k)}}{\gamma_i}\right) + \frac{Y_{ig}^{(k)} - \mu_{ig}^{(k)}}{\mu_{ig}^{(k)}}$ and $\phi_g^{(k)}$ is the vector of $(\phi_{ig}^{(k)})_{1 \le i \le n_k}$. | Each center | Server |
| | local_hat_matrix | nparray | $(G_{\texttt{act}}, p, p)$ | For each gene $g$, the hat matrix $H_g^{(k)} = (X^{(k)})^\top W_g^{(k)} X^{(k)}$ where $W_g^{(k)} \in \mathbb{R}^{n_k \times n_k}$ is the diagonal matrix with diagonal entries $\frac{\mu_{ig}^{(k)}}{1+\mu_{ig}^{(k)} \alpha_g}$ for $1 \le i \le n_k$. $\alpha_g$ is the dispersion estimate of the gene and $\mu_{ig}^{(k)}$ is the expected value of the gene for sample [illegible] for parameter $\beta$, that is $\gamma_g^{(k)} \exp(X_i^{(k)} \cdot \beta_g)$. | Each center | Server |

| | | | | | | |
|---|---|---|---|---|---|---|
| | local_nll | nparray | $(G_{\text{act}},)$ | For each gene $g$, the negative log likelihood of the negative binomial GLM with fixed dispersion at the $\beta_g$ parameter, computed on the local data. | Each center | Server |
| | irls_gene_names | Index | $(G_{\text{act}},)$ | The gene names of the active genes. | Each center | Server |
| 4 | round_number_irls | int | | The current round number of the IRLS algorithm, incremented by 1 during the local step. | Server | Center |
| | global_nll | nparray | $(G_{\text{nz}},)$ | For each gene $g$, the negative log-likelihood at the $\beta_g$ parameter, built by summing the local nlls from the centers | Server | Center |
| | beta | nparray | $(G_{\text{nz}}, p)$ | The updated value of the $\beta$ parameter after applying the IRLS update for the active genes. At the last step, is not updated. | Server | Center |
| | irls_diverged_mask | nparray | $(G_{\text{nz}},)$ | A boolean array indicating which non zero genes have caused the IRLS algorithm to diverge. A gene $g$ is considered to have diverged if any component of $\beta_g \in \mathbb{R}^p$ has absolute value above the threshold `max_beta`. | Server | Center |
| | irls_mask | nparray | $(G_{\text{nz}},)$ | A boolean array indicating which non zero genes are currently being optimized by the IRLS algorithm. Genes whose $\beta$ parameter has diverged are removed from the optimization. Moreover, the deviance ratio is computed for each active gene $g$, as $\frac{\lvert 2\,\mathcal{L}(\beta_g) - 2\cdot\mathcal{L}(\beta_{g,\text{prev}})\rvert}{\lvert 2\,\mathcal{L}(\beta)\rvert + 0.1}$ where $\beta_{g,\text{prev}}$ is the value of $\beta_g$ at the previous iteration. If this deviance is smaller than the user inputed `beta_tol`, the gene is considered to have converged and is set to `False` in the `irls_mask`, meaning it will not be optimized further. | Server | Center |
| 5 | ascent_direction_on_mask | NoneType | | The ascent direction on the active genes, which is set to `None` at the beginning. | Each center | Server |
| | irls_diverged_mask | nparray | $(G_{\text{nz}},)$ | A boolean array indicating which non zero genes have caused the IRLS algorithm to diverge. Simply passed on during the Fed Proximal Quasi-Newton algorithm, since the server is stateless. | Each center | Server |
| | round_number_PQN | int | | The current round number of the Proximal Quasi-Newton algorithm, initialized at 0. | Each center | Server |
| | newton_decrement_on_mask | NoneType | | The newton decrement on the active genes, which is set to `None` at the beginning. | Each center | Server |
| | PQN_mask | nparray | $(G_{\text{nz}},)$ | A boolean array indicating which non zero genes are currently being optimized by the Proximal Quasi-Newton algorithm (active genes). The genes which have diverged during the IRLS algorithm and those which have not converged at the end of the prescribed iterations are set to `True`. | Each center | Server |
| | global_reg_nll | nparray | $(G_{\text{nz}},)$ | The regularized negative log likelihood of the negative binomial GLM with fixed dispersion at the $\beta_g$ parameter intialized at `np.nan` since no nll has been computed at this stage. | Each center | Server |
| | local_gradient | nparray | $(1, G_{\text{act}}, p)$ | For each gene $g$, the gradient of the negative log likelihood of the negative binomial GLM with fixed dispersion at the $\beta_g$ parameter, computed on the local data, i.e., $\nabla_\beta \mathcal{L}^{(k)}(\beta_g) = -(X^{(k)})^\top Y_g^{(k)} + (X^{(k)})^\top \left(\frac{1}{\alpha_g} + Y_g^{(k)}\right) \frac{\mu_g^{(k)}}{\frac{1}{\alpha_g} + \mu_g^{(k)}}$, where $\mu_g^{(k)}$ and $Y_g^{(k)}$ are both vectors in $\mathbb{R}^{n_k}$ and $\alpha_g$ is a scalar. TODO ref equation. | Each center | Server |
| | local_fisher | nparray | $(1, G_{\text{act}}, p, p)$ | For each gene $g$, the Fisher information matrix of the negative binomial GLM with fixed dispersion, $n_k \mathcal{I}_g^{(k)} = (X^{(k)})^\top W_g^{(k)} X^{(k)}$ where $W_g^{(k)} \in \mathbb{R}^{n_k \times n_k}$ is the diagonal matrix with diagonal entries $W_{gii}^{(k)} = \frac{\mu_{ig}^{(k)}}{1+\mu_{ig}^{(k)}\alpha_g}$ for $1 \leq i \leq n_k$. $\alpha_g$ is the dispersion estimate of the gene and $\mu_{ig}^{(k)}$ is the expected value of the gene $g$ for sample [illegible] for parameter $\beta_g$, that is $\gamma_g^{(k)} \exp(X_i^{(k)} \cdot \beta_g)$. | Each center | Server |
| | local_nll | nparray | $(1, G_{\text{act}})$ | The nll of the negative binomial GLM with fixed dispersion at the $\beta_g$ parameter, computed on the local data, for each gene $g$. | Each center | Server |
| | beta | nparray | $(G_{\text{nz}}, p)$ | For evert gene $g$, the inital value of the $\beta_g$ parameter for the ProxQuasiNewton algorithm (PQN), which is either i) the value computed by IRLS if IRLS has converged on that gene (in that case, the gene will not be optimized) or ii) the initial value of the $\beta_g$ parameter for the IRLS algorithm, if IRLS has not converged on that gene. | Each center | Server |
| | PQN_diverged_mask | nparray | $(G_{\text{nz}},)$ | A boolean array indicating which non zero genes have caused the Proximal Quasi-Newton algorithm to diverge. Initialized to `False`. | Each center | Server |
| 6 | round_number_PQN | int | | The current round number of the Proximal Quasi-Newton algorithm, incremented by 1. | Server | Center |
| | irls_diverged_mask | nparray | $(G_{\text{nz}},)$ | A boolean array indicating which non zero genes have caused the IRLS algorithm to diverge. Passed on without modification. | Server | Center |
| | global_reg_nll | nparray | $(G_{\text{nz}},)$ | The regularized negative log likelihood of the negative binomial GLM with fixed dispersion at the $\beta_g$ parameter computed as the sum of the local negative log likelihoods with $L^2$ regularization with $\lambda = 10^{-6}$. | Server | Center |
| | newton_decrement_on_mask | nparray | $(G_{\text{act}},)$ | The newton decrement on the active genes. It is computed as $\nu_g = \Delta_g \cdot \nabla_\beta \mathcal{L}^\lambda(\beta_g)$ where $\Delta_g$ is the ascent direction on the active genes and $\mathcal{L}^\lambda$ is the regularized log likelihood. | Server | Center |
| | ascent_direction_on_mask | nparray | $(G_{\text{act}}, p)$ | The ascent direction on the active genes. To compute the ascent direction, we first compute the global Fisher information matrix for each gene $g$, $n\mathcal{I}_g = \sum_k n_k \mathcal{I}_g^{(k)}$ and the global gradient $\nabla_\beta \mathcal{L}(\beta_g) = \sum_k \nabla_\beta \mathcal{L}^{(k)}(\beta_g)$. From these two quantities, we build the Fisher information and gradients of the regularized negative log likelihood, with $L^2$ regularization $\lambda = 10^{-6}$. We add another regularization term to the Fisher matrix which depends on the iteration number. The goal is for the ascent direction to be close to the gradient for small iterations and to be close to the real natural gradient descent update near the end of the optimization. The ascent direction is then computed as from the regularized Fisher and gradient (see main paper). Roughly, it is the solution of the linear system $n\,\mathcal{I}_g^\lambda \Delta_g = \nabla_\beta \mathcal{L}^\lambda(\beta_g)$, where certain components are dropped in to handle boundary conditions as the optimization is constrained to values of $\beta_g$ in $[-\texttt{max\_beta}, \texttt{max\_beta}]$. | Server | Center |
| | PQN_diverged_mask | nparray | $(G_{\text{nz}},)$ | A boolean array indicating which non zero genes have caused the Proximal Quasi-Newton algorithm to diverge. Passed on without modification. | Server | Center |

| | | | | | | |
|---|---|---|---|---|---|---|
| | PQN_mask | nparray | $(G_{\mathtt{nz}},)$ | A boolean array indicating which non zero genes are currently being optimized by the Proximal Quasi-Newton algorithm (active genes), passed on without modification. | Server | Center |
| | beta | nparray | $(G_{\mathtt{nz}}, p)$ | For each gene $g$, the log fold change $\beta_g$, which remains unchanged during the first iteration of the Proximal Quasi-Newton algorithm. | Server | Center |
| 7 | ascent_direction_on_mask | nparray | $(G_{\mathtt{act}}, p)$ | The ascent direction on the active genes, passed on without modification. | Each center | Server |
| | irls_diverged_mask | nparray | $(G_{\mathtt{nz}},)$ | A boolean array indicating which non zero genes have caused the IRLS algorithm to diverge. Passed on without modification. | Each center | Server |
| | round_number_PQN | int | | The current round number of the Proximal Quasi-Newton algorithm, passed on without modification. | Each center | Server |
| | newton_decrement_on_mask | nparray | $(G_{\mathtt{act}},)$ | The newton decrement on the active genes, passed on without modification. | Each center | Server |
| | global_reg_nll | nparray | $(G_{\mathtt{nz}},)$ | The regularized negative log likelihood of the negative binomial GLM with fixed dispersion at the $\beta_g$ parameter simply passed on. | Each center | Server |
| | PQN_mask | nparray | $(G_{\mathtt{nz}},)$ | A boolean array indicating which non zero genes are currently being optimized by the Proximal Quasi-Newton algorithm (active genes), passed on without modification. | Each center | Server |
| | PQN_diverged_mask | nparray | $(G_{\mathtt{nz}},)$ | A boolean array indicating which non zero genes have caused the Proximal Quasi-Newton algorithm to diverge Passed on without modification. | Each center | Server |
| | local_gradient | nparray | $(N_{\mathtt{ls}}, G_{\mathtt{act}}, p)$ | For each potential step size $\delta$ and each gene $g$, the gradient of the negative log likelihood of the negative binomial GLM with fixed dispersion at the $\beta_g - \delta\Delta_g$ parameter computed on the local data. | Each center | Server |
| | local_fisher | nparray | $(N_{\mathtt{ls}}, G_{\mathtt{act}}, p, p)$ | For each potential step size $\delta \in \{1, 1/2, 1/4, ..., 1/2^{N_{\mathtt{ls}}-1}\}$, and each gene $g$, the Fisher information matrix of the negative binomial GLM with fixed dispersion computed at $\beta_g - \delta\Delta_g$ Formally, $n_k \mathcal{I}^{(k)}_{\delta,g} = (X^{(k)})^\top W^{(k)}_{\delta,g} X^{(k)}$ where $W^{(k)}_{\delta,g} \in \mathbb{R}^{n_k \times n_k}$ is the diagonal matrix with diagonal entries $W^{(k)}_{\delta,gii} = \frac{\mu^{(k)}_{\delta,ig}}{1+\mu^{(k)}_{\delta,ig}\alpha_g}$ for $1 \leq i \leq n_k$. $\alpha_g$ is the dispersion estimate of the gene and $\mu^{(k)}_{\delta,ig}$ is the expected value of the gene $g$ for sample $i$ for parameter $\beta_g - \delta\Delta_g$, that is $\gamma^{(k)}_g \exp(X^{(k)}_i \cdot (\beta_g - \delta\Delta_g))$. | Each center | Server |
| | local_nll | nparray | $(N_{\mathtt{ls}}, G_{\mathtt{act}})$ | The nll of the negative binomial GLM with fixed dispersion at the $\beta_g$ parameter, computed on the local data, for each gene $g$ and for each potential next step $\beta_g - \delta\Delta_g$ for $\delta \in \{1, 1/2, 1/4, ..., 1/2^{N_{\mathtt{ls}}-1}\}$. | Each center | Server |
| | beta | nparray | $(G_{\mathtt{nz}}, p)$ | For each gene $g$, the log fold change $\beta_g$, which is simply passed on. | Each center | Server |
| 8 | irls_diverged_mask | nparray | $(G_{\mathtt{nz}},)$ | A boolean array indicating which non zero genes have caused the IRLS algorithm to diverge. Passed on without modification. | Server | Center |
| | global_reg_nll | nparray | $(G_{\mathtt{nz}},)$ | The regularized negative log likelihood of the negative binomial GLM with fixed dispersion at the $\beta_g$ parameter computed as the sum of the local negative log likelihoods with $L^2$ regularization with $\lambda = 10^{-6}$. | Server | Center |
| | round_number_PQN | int | | The current round number of the Proximal Quasi-Newton algorithm, incremented by 1. | Server | Center |
| | newton_decrement_on_mask | nparray | $(G_{\mathtt{act}},)$ | The newton decrement on the new active genes $g$, computed at the new $\beta_g$ iterate. For more details, see description of 19. | Server | Center |
| | ascent_direction_on_mask | nparray | $(G_{\mathtt{act}}, p)$ | The ascent direction on the new active genes $g$, computed at the new $\beta_g$ iterate. | Server | Center |
| | PQN_diverged_mask | nparray | $(G_{\mathtt{nz}},)$ | A boolean array indicating which non zero genes have caused the Proximal Quasi-Newton algorithm to diverge A gene $g$ is considered to have diverged if for all step sizes $\delta \in \{1, 1/2, 1/4, ..., 1/2^{N_{\mathtt{ls}}-1}\}$, and for the computed ascent direction $\Delta_g$, the Armijo condition is not satisfied i.e., if $\mathcal{L}^\lambda(\beta_g) - \mathcal{L}^\lambda(\beta_g - \delta\Delta_g) \leq \delta\ \mathtt{PQN\_c1}\ \nabla_\beta \mathcal{L}^\lambda(\beta_g) \cdot \Delta_g$ where $\mathcal{L}^\lambda$ is the regularized negative log likelihood, `PQN_c1` is the Armijo parameter which can be set by the user (default is $10^{-4}$), and $\nabla_\beta \mathcal{L}^\lambda(\beta_g)$ is the gradient of the regularized negative log likelihood. | Server | Center |
| | PQN_mask | nparray | $(G_{\mathtt{nz}},)$ | A boolean array indicating which non zero genes are currently being optimized by the Proximal Quasi-Newton algorithm (active genes). Genes that had converged with the relative error criterion or diverged (no stepsize satisfying the Armijo condition) have set to `False`. | Server | Center |
| | beta | nparray | $(G_{\mathtt{nz}}, p)$ | For each gene $g$, the log fold change $\beta_g$. This value has been updated by the Proximal Quasi-Newton algorithm for all active genes where an admissible step size was found using the Armijo condition (see equation 2.5 of [1]) | Server | Center |
| 9 | ascent_direction_on_mask | nparray | $(G_{\mathtt{act}}, p)$ | The ascent direction on the active genes, passed on without modification. | Each center | Server |
| | irls_diverged_mask | nparray | $(G_{\mathtt{nz}},)$ | A boolean array indicating which non zero genes have caused the IRLS algorithm to diverge. Passed on without modification. | Each center | Server |
| | round_number_PQN | int | | The current round number of the Proximal Quasi-Newton algorithm, passed on without modification. | Each center | Server |
| | newton_decrement_on_mask | nparray | $(G_{\mathtt{act}},)$ | The newton decrement on the active genes, passed on without modification. | Each center | Server |
| | global_reg_nll | nparray | $(G_{\mathtt{nz}},)$ | The regularized negative log likelihood of the negative binomial GLM with fixed dispersion at the $\beta_g$ parameter simply passed on. | Each center | Server |
| | PQN_mask | nparray | $(G_{\mathtt{nz}},)$ | A boolean array indicating which non zero genes are currently being optimized by the Proximal Quasi-Newton algorithm (active genes), passed on without modification. | Each center | Server |
| | PQN_diverged_mask | nparray | $(G_{\mathtt{nz}},)$ | A boolean array indicating which non zero genes have caused the Proximal Quasi-Newton algorithm to diverge Passed on without modification. | Each center | Server |
| | local_gradient | nparray | $(N_{\mathtt{ls}}, G_{\mathtt{act}}, p)$ | For each potential step size $\delta$ and each gene $g$, the gradient of the negative log likelihood of the negative binomial GLM with fixed dispersion at the $\beta_g - \delta\Delta_g$ parameter computed on the local data. | Each center | Server |

| | | | | | | |
|---|---|---|---|---|---|---|
| | local_fisher | nparray | $(N_{\texttt{ls}}, G_{\texttt{act}}, p, p)$ | For each potential step size $\delta \in \{1, 1/2, 1/4, \dots, 1/2^{N_{\texttt{ls}}-1}\}$, and each gene $g$, the Fisher information matrix of the negative binomial GLM with fixed dispersion computed at $\beta_g - \delta\Delta_g$. Formally, $n_k \mathcal{I}^{(k)}_{\delta,g} = (X^{(k)})^\top W^{(k)}_{\delta,g} X^{(k)}$ where $W^{(k)}_{\delta,g} \in \mathbb{R}^{n_k \times n_k}$ is the diagonal matrix with diagonal entries $W^{(k)}_{\delta,gii} = \frac{\mu^{(k)}_{\delta,ig}}{1+\mu^{(k)}_{\delta,ig}\alpha_g}$ for $1 \leq i \leq n_k$. $\alpha_g$ is the dispersion estimate of the gene and $\mu^{(k)}_{\delta,ig}$ is the expected value of the gene $g$ for sample $i$ for parameter $\beta_g - \delta\Delta_g$, that is $\gamma^{(k)}_g \exp(X^{(k)}_i \cdot (\beta_g - \delta\Delta_g))$. | Each center | Server |
| | local_nll | nparray | $(N_{\texttt{ls}}, G_{\texttt{act}})$ | The nll of the negative binomial GLM with fixed dispersion at the $\beta_g$ parameter, computed on the local data, for each gene $g$ and for each potential next step $\beta_g - \delta\Delta_g$ for $\delta \in \{1, 1/2, 1/4, \dots, 1/2^{N_{\texttt{ls}}-1}\}$. | Each center | Server |
| | beta | nparray | $(G_{\texttt{nz}}, p)$ | For each gene $g$, the log fold change $\beta_g$, which is simply passed on. | Each center | Server |
| 10 | beta | nparray | $(G_{\texttt{nz}}, p)$ | For each gene $g$, the log fold change $\beta_g$. This value has been updated by the Proximal Quasi-Newton algorithm for all active genes where an admissible step size was found using the Armijo condition (see equation 2.5 of [1]) | Server | Center |
| | PQN_diverged_mask | nparray | $(G_{\texttt{nz}}, )$ | A boolean array indicating which non zero genes have caused the Proximal Quasi-Newton algorithm to diverge. As this is the last step of the optimization, a gene $g$ is considered to have diverged not only if the Armijo condition is never satisfied, but also if the convergence criterion is not met. A gene is said to converge if the relative difference between two successive iterates is smaller than a threshold value PQN_ftol, i.e., $\left\lvert \mathcal{L}^\lambda(\beta_g) - \mathcal{L}^\lambda(\beta_{g,\text{prev}}) \right\rvert \leq$ PQN_ftol $\max(\mathcal{L}^\lambda(\beta_g), \mathcal{L}^\lambda(\beta_{g,\text{prev}}), 1)$. | Server | Center |
| | irls_diverged_mask | nparray | $(G_{\texttt{nz}}, )$ | A boolean array indicating which non zero genes have caused the IRLS algorithm to diverge. Passed on without modification. | Server | Center |
