## Supplementary material for "FedPyDESeq2: a federated framework for bulk RNA-seq differential expression analysis": Workflow Diagram

### Supplementary material: precise description of the shared quantities between centers and server

In the context of federated learning, which aims to enhance privacy by keeping sensitive data local, transparency regarding data exchange is crucial. We provide a comprehensive workflow graph and detailed table to explicitly document every quantity shared between the centers and the central server. This transparency is essential for privacy assessment, allowing stakeholders to clearly understand what information leaves their local environment and what aggregated data they receive.

The complete workflow graph and its associated table present the federated implementation of the DESeq2 pipeline. In the graph, white squares represent quantities exchanged between centers and the server, with reference numbers in the table providing detailed descriptions of each exchange. Pink blocks indicate functions executed at the center level, while blue blocks represent operations performed at the server level. The workflow is organized into functional groups, denoted by dashed blocks, which correspond to specific routines implemented in the FedPyDESeq2 package.

### Workflow graph

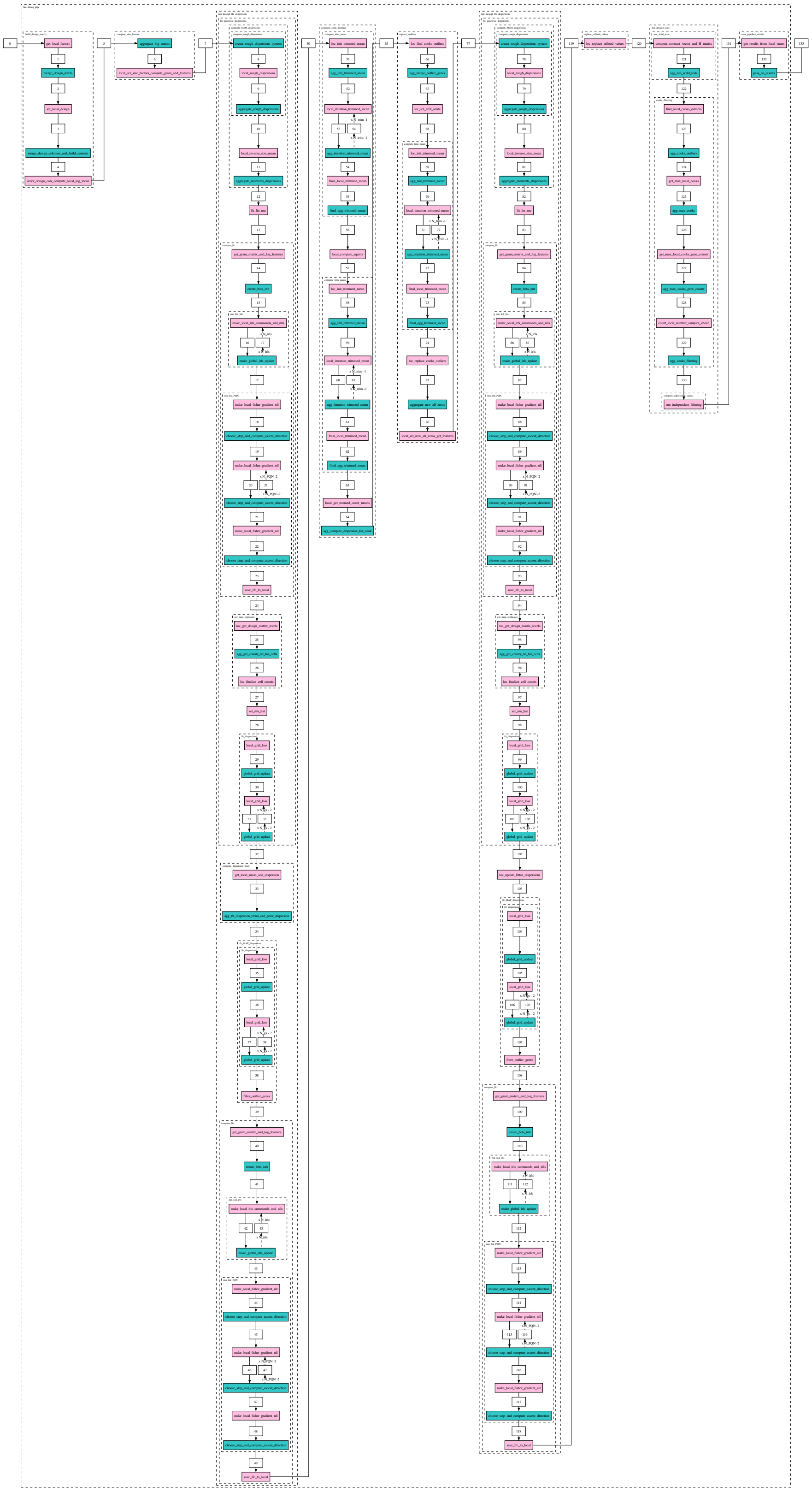

Figure 1: Complete workflow of the FedPyDESeq2 implementation. White squares represent exchanged quantities, pink blocks indicate center-level functions, blue blocks represent server-level operations, dashed blocks denote functional groups.

### Table of exchanged quantities

The table below presents a detailed description of the quantities exchanged during the federated DESeq2 analysis. Each row is identified by an ID that corresponds to the numbered white squares in the workflow graph (Figure 1), allowing direct mapping between the visual representation and the detailed specifications. The table provides the name and type of each exchanged quantity, along with its shape when applicable. The "Description" column offers mathematical definitions and contextual explanations for each quantity. The "Shared with" column indicates whether the quantity is shared with the central server or with the centers.

- $K$: the total number of centers. Centers will be indexed by the letter $k$.
- $p$: the number of parameters in the linear model (number of columns in the design matrix). They will be indexed by the letter $j$.
- $n_k$: the number of observations in center $k$, and $n := \sum_k n_k$ the total number of observations (samples). Samples in any center $k$ will be indexed by the letter $i$.
- $G$ the total number of input genes. Genes will be indexed by the letter $g$. We will also use $G_{\texttt{nz}}$ the number of non-zero genes (computed during the algorithm), which are the genes with at least one sample with non-zero counts, and $G_{\texttt{act}}$, the number of active genes, which will be the number of genes which are still being optimized in a given algorithm.
- $X^{(k)} \in \mathbb{R}^{n_k \times p}$: the design matrix of center $k$. We will denote with $X_{ij}^{(k)}$ the value of the $i$-th row and $j$-th column of the design matrix of center $k$. The local design matrices are built from the input parameters during the first steps of the algorithm.
- $\gamma^{(k)} \in \mathbb{R}^{n_k}$ : the size factors of the samples of center $k$, indexed by samples $i$. $\gamma_i^{(k)}$ denotes the size factor of sample $i$ of center $k$.
- $Y^{(k)} \in \mathbb{R}^{n_k \times G}$ the count matrix of center $k$, indexed by sample number $i$ and gene $g$, s.t. $Y_{ig}^{(k)}$ denotes the count of gene $g$ in sample $i$ of center $k$.
- $Z_{ig}^{(k)} \in \mathbb{R}^{n_k \times G}$ the normalized count matrix indexed by samples $i$ and gene $g$, s.t. $Z_{ig}^{(k)} := Y_{ig}^{(k)}/\gamma_i^{(k)}$ denotes the normalized count of gene $g$ in sample $i$ of center $k$.

| ID | Name | Type | Shape | Description | Computed by | Sent to |
|---|---|---|---|---|---|---|
| 1 | local_levels | dict | | A dictionary whose keys are the names of the categorical factors, and whose values are the list of values taken by this factor in a given center. For example `{stage: [Advanced, Non-advanced], gender: [female]}`. | Each center | Server |
| 2 | merged_levels | dict | | A dictionary whose keys are the names of the categorical factors and whose values are arrays containing the list of values taken by this factor across all centers. For example `{stage: [Advanced , Non-advanced], gender: [female, male]}`. | Server | Center |
| 3 | design_columns | Index | $(p,)$ | The name of the columns of the local design matrix, before aggregation. They are of the form intercept, factor for continuous factors, and factor_level_vs_factor_ref_level otherwise. For example `[intercept, stage_Advanced_vs_Non-advanced, gender_male_vs_female]` | Each center | Server |
| 4 | merged_columns | Index | $(p,)$ | The union of the design columns across all centers. Local design matrices will then be updated to include all columns. | Server | Center |
| | contrast | list | | A list of three strings representing the contrast of interest, in case it is not specified by the user. Of the form `[factor, level1, level2]`. For example `[stage, Advanced, Non-advanced]`. | Server | Center |
| 5 | log_mean | nparray | $(G,)$ | For each gene, the mean of the log of the counts across all samples in a center $\overline{\log(Y)}_g^{(k)} = \frac{1}{n_k}\sum_{i=1}^{n_k}\log(Y_{ig}^{(k)})$ | Each center | Server |
| | n_samples | int | | The number of samples in a center $n_k$ for each center $k$. | Each center | Server |
| 6 | global_log_mean | nparray | $(G,)$ | The mean of the log of the counts across all samples in all centers $\overline{\log(Y)}_g = \sum_{k=1}^{K}\frac{n_k}{n}\overline{\log(Y)}_g^{(k)}$. | Server | Center |
| 7 | local_gram_matrix | nparray | $(p,p)$ | The gram matrix of the local design matrix $G^{(k)} := (X^{(k)})^\top X^{(k)}$. | Each center | Server |
| | local_features | nparray | $(p,G)$ | $\Phi^{(k)} := X^{(k)\top} Z^{(k)}$. | Each center | Server |
| 8 | global_feature_vector | nparray | $(p,G)$ | The sum of the local features across all centers $\Phi = \sum_k \Phi^{(k)}$. | Server | Center |
| | global_gram_matrix | nparray | $(p,p)$ | The sum of the local Gramm matrix across all centers $G = \sum_k G^{(k)}$. | Server | Center |
| 9 | local_rough_dispersions | nparray | $(G,)$ | A rough estimate of the dispersions of the genes on the local data. It is computed as a method of moments estimator of the dispersion of the normalized counts $(\alpha_g^{\texttt{MoM},2})^{(k)} = \sum_{i=1}^{n_k}((Z_{ig}^{(k)})^2 - (\widehat{Z}_{ig}^{(k)})^2)/(\widehat{Z}_{ig}^{(k)})^2$ where $\widehat{Z}_g^{(k)} = X^{(k)}G^{-1}\Phi_g$ corresponds to a linear estimate of the normed counts. | Each center | Server |
| | local_n_obs | int | | The number of observations in a center $n_k$. | Each center | Server |
| | local_n_params | int | | The number of parameters in a center $p$ (i.e., the number of columns of the design matrix). This number is the same across all centers, but the server does not directly have access to it since it is stateless. | Each center | Server |
| 10 | rough_dispersions | nparray | $(G,)$ | A rough estimate of the dispersions obtained by essentially averaging local rough dispersions ($\alpha_g^{\texttt{MoM},2} = \max(\frac{1}{n-p}\sum_k(\alpha_g^{\texttt{MoM},2})^{(k)}, 0)$) | Server | Center |
| 11 | local_inverse_size_mean | float | $()$ | The average of the inverse of the size factors in a center $\overline{1/\gamma}^{(k)} = \frac{1}{n_k}\sum_{i=1}^{n_k} 1/\gamma_i^{(k)}$. | Each center | Server |
| | local_counts_mean | nparray | $(G,)$ | The averag e of the counts in a center, per gene $\overline{Y}_g^{(k)} = \frac{1}{n_k}\sum_{i=1}^{n_k} Y_{ig}^{(k)}$. | Each center | Server |
| | local_squared_squared_mean | nparray | $(G,)$ | The average of the square normed counts in a center, per gene $\overline{Y^2} := \frac{1}{n_k}\sum_{i=1}^{n_k}(Y_{ig}^{(k)})^2$. | Each center | Server |

| | | | | | | |
|---|---|---|---|---|---|---|
| | local_n_obs | int | | The number of observations in a center $n_k$. | Each center | Server |
| | rough_dispersions | nparray | $(G, )$ | The rough estimates computed at the previous step anc passed on. | Each center | Server |
| 12 | MoM_dispersions | nparray | $(G, )$ | The dispersions obtained by the method of moments. First, the aggregated mean counts, squarec mean counts and inverse size factors are computec ($\overline{1/\gamma} = \sum_k \frac{n_k}{n} \overline{1/\gamma}^{(k)}$), $\overline{Y}_g = \sum_k \frac{n_k}{n} \overline{Y}_g^{(k)}$, $\overline{Y^2} = \sum_k \frac{n_k}{n} \overline{Y^2}_g^{(k)}$)). The variance is then estimated with $\hat{\sigma}_g^2 = \frac{n}{n-1}(\overline{Y^2}_g - \overline{Y}_g^2)$ and a method of moments estimate is the constructed as $\alpha_g^{\mathtt{MoM},1} = \frac{\hat{\sigma}_g^2 - \overline{1/\gamma}_g \times \overline{Y}_g}{\overline{Y}_g^2}$. The final MoM estimate $\alpha_g^{\mathtt{MoM}}$ for gene $g$ is the clipped value o: $\min(\alpha_g^{\mathtt{MoM},1}, \alpha_g^{\mathtt{MoM},2})$ between `min_disp` and $\max($`max_disp`$, n)$ | Server | Center |
| | non_zero | nparray | $(G, )$ | A boolean array indicating which genes have non-zerc counts in at least one center. | Server | Center |
| | tot_num_samples | int | | The total number of samples across all centers $n$. | Server | Center |
| | tot_counts_mean | nparray | $(G, )$ | The average of the counts across all centers, per gene obtained by summing the local counts means. | Server | Center |
| 14 | gram_full_rank | bool | $()$ | A boolean indicating whether the global Gramm matrix $G$ is full rank. | Each center | Server |
| | n_non_zero_genes | int | | The number of genes with non-zero counts in at least one center. | Each center | Server |
| | local_log_features | nparray | $(G_{\mathtt{nz}}, p)$ | $\Phi_k = (X^{(k)})^\top \log(Z^{(k)} + 0.1)$. | Each center | Server |
| | global_gram_matrix | nparray | $(p, p)$ | The sum of the local Gramm matrix across all centers $G = \sum_k G^{(k)}$. | Each center | Server |
| 15 | round_number_irls | int | | The current round number of the IRLS algorithm. Initialized at 0. | Server | Center |
| | global_nll | nparray | $(G_{\mathtt{nz}}, )$ | For each gene $g$, the negative log likelihood of the negative binomial GLM, initialized at 1000 for all genes. | Server | Center |
| | irls_mask | nparray | $(G_{\mathtt{nz}}, )$ | A boolean array indicating which non zero genes are currently being optimized by the IRLS algorithm. Initializec to `True`. | Server | Center |
| | irls_diverged_mask | nparray | $(G_{\mathtt{nz}}, )$ | A boolean array indicating which non zero genes have caused the IRLS algorithm to diverge. Initialized to `False` | Server | Center |
| | beta | nparray | $(G_{\mathtt{nz}}, p)$ | The initial value of the $\beta$ parameter. If the Gramm matrix is full rank, is set to $\beta_g = G^{-1}(\sum_k \Phi_g^{(k)})$ for al genes $g$. Otherwise, it is set as a weighed average of the normed log means, i.e., $\beta_g = \sum_k \frac{n_k}{n} \overline{\log(Z)}_g^{(k)}$, where $\overline{\log(Z)}_g^{(k)} = \frac{1}{n_k} \sum_{i=1}^{n_k} \log(Z_{ig}^{(k)})$ is computed locally. | Server | Center |
| 16 | beta | nparray | $(G_{\mathtt{nz}}, p)$ | The log fold change $\beta$ parameter. Not modified. | Each center | Server |
| | local_nll | nparray | $(G_{\mathtt{act}}, )$ | For each gene $g$, the negative log likelihood of the negative binomial GLM with fixed dispersion at the $\beta_g$ parameter, computed on the local data. | Each center | Server |
| | local_hat_matrix | nparray | $(G_{\mathtt{act}}, p, p)$ | For each gene $g$, the hat matrix $H_g^{(k)} = (X^{(k)})^\top W_g^{(k)} X^{(k)}$ where $W_g^{(k)} \in \mathbb{R}^{n_k \times n_k}$ is the diagonal matrix with diagonal entries $\frac{\mu_{ig}^{(k)}}{1+\mu_{ig}^{(k)} \alpha_g}$ fo: $1 \leq i \leq n_k$. $\alpha_g$ is the dispersion estimate of the gene and $\mu_{ig}^{(k)}$ is the expected value of the gene for sample : for parameter $\beta$, that is $\gamma_g^{(k)} \exp(X_i^{(k)} \cdot \beta_g)$. | Each center | Server |
| | local_features | nparray | $(G_{\mathtt{act}}, p)$ | For each active gene $g$, the projected features $(X^{(k)})^\top W_g^{(k)} \phi_g^{(k)}$, where $\phi_{ig}^{(k)} = \log\left(\frac{\mu_{ig}^{(k)}}{\gamma_i}\right) + \frac{Y_{ig}^{(k)} - \mu_{ig}^{(k)}}{\mu_{ig}^{(k)}}$ and $\phi_g^{(k)}$ is the vector of $(\phi_{ig}^{(k)})_{1 \leq i \leq n_k}$. | Each center | Server |
| | irls_gene_names | Index | $(G_{\mathtt{act}}, )$ | The gene names of the active genes. | Each center | Server |
| | irls_diverged_mask | nparray | $(G_{\mathtt{nz}}, )$ | A boolean array indicating which non zero genes have caused the IRLS algorithm to diverge. Simply passed on | Each center | Server |
| | irls_mask | nparray | $(G_{\mathtt{nz}}, )$ | A boolean array indicating which non zero genes are currently being optimized by the IRLS algorithm. Simply passed on. | Each center | Server |
| | global_nll | nparray | $(G_{\mathtt{nz}}, )$ | For each gene $g$, the negative log likelihood at the $\beta_g$ parameter. Simply passed on. | Each center | Server |
| | round_number_irls | int | | The current round number of the IRLS algorithm. Simply passed on. | Each center | Server |
| 17 | global_nll | nparray | $(G_{\mathtt{nz}}, )$ | For each gene $g$, the negative log-likelihood at the $\beta_g$ parameter, built by summing the local nlls from the centers | Server | Center |
| | irls_mask | nparray | $(G_{\mathtt{nz}}, )$ | A boolean array indicating which non zero genes are currently being optimized by the IRLS algorithm. Genes whose $\beta$ parameter has diverged are removed from the optimization. Moreover, the deviance ratio is computec for each active gene $g$, as $\frac{\lvert 2\,\mathcal{L}(\beta_g) - 2 \cdot \mathcal{L}(\beta_{g,\text{prev}})\rvert}{\lvert 2\,\mathcal{L}(\beta)\rvert + 0.1}$ where $\beta_{g,\text{prev}}$ is the value of $\beta_g$ at the previous iteration. I: this deviance is smaller than the user inputed `beta_tol`, the gene is considered to have converged and is set to `False` ir the `irls_mask`, meaning it will not be optimized further. | Server | Center |
| | round_number_irls | int | | The current round number of the IRLS algorithm, incremented by 1 during the local step. | Server | Center |
| | beta | nparray | $(G_{\mathtt{nz}}, p)$ | The updated value of the $\beta$ parameter after applying the IRLS update for the active genes. At the last step, is not updated. | Server | Center |
| | irls_diverged_mask | nparray | $(G_{\mathtt{nz}}, )$ | A boolean array indicating which non zero genes have caused the IRLS algorithm to diverge. A gene $g$ is considered to have diverged if any component of $\beta_g \in \mathbb{R}^p$ has absolute value above the threshold `max_beta`. | Server | Center |
| 18 | ascent_direction_on_mask | NoneType | | The ascent direction on the active genes, which is set tc `None` at the beginning. | Each center | Server |
| | irls_diverged_mask | nparray | $(G_{\mathtt{nz}}, )$ | A boolean array indicating which non zero genes have caused the IRLS algorithm to diverge. Simply passed or during the Fed Proximal Quasi-Newton algorithm, since the server is stateless. | Each center | Server |
| | round_number_PQN | int | | The current round number of the Proximal Quasi-Newtor algorithm, initialized at 0. | Each center | Server |
| | newton_decrement_on_mask | NoneType | | The newton decrement on the active genes, which is set to `None` at the beginning. | Each center | Server |
| | PQN_mask | nparray | $(G_{\mathtt{nz}}, )$ | A boolean array indicating which non zero genes are currently being optimized by the Proximal Quasi-Newton algorithm (active genes). The genes which have divergec during the IRLS algorithm and those which have not converged at the end of the prescribed iterations are set tc `True`. | Each center | Server |
| | global_reg_nll | nparray | $(G_{\mathtt{nz}}, )$ | The regularized negative log likelihood of the negative binomial GLM with fixed dispersion at the $\beta_g$ parameter intialized at `np.nan` since no nll has been computed at this stage. | Each center | Server |

| | | | | | | |
|---|---|---|---|---|---|---|
| | local_gradient | nparray | $(1, G_{\texttt{act}}, p)$ | For each gene $g$, the gradient of the negative log likelihood of the negative binomial GLM with fixed dispersion at the $\beta_g$ parameter, computed on the local data, i.e., $\nabla_\beta \mathcal{L}^{(k)}(\beta_g) = -(X^{(k)})^\top Y_g^{(k)} + (X^{(k)})^\top \left(\frac{1}{\alpha_g} + Y_g^{(k)}\right) \frac{\mu_g^{(k)}}{\frac{1}{\alpha_g}+\mu_g^{(k)}}$, where $\mu_g^{(k)}$ anc $Y_g^{(k)}$ are both vectors in $\mathbb{R}^{n_k}$ and $\alpha_g$ is a scalar. TODC ref equation. | Each center | Server |
| | local_fisher | nparray | $(1, G_{\texttt{act}}, p, p)$ | For each gene $g$, the Fisher information matrix of the negative binomial GLM with fixed dispersion, $n_k \mathcal{I}_g^{(k)} = (X^{(k)})^\top W_g^{(k)} X^{(k)}$ where $W_g^{(k)} \in \mathbb{R}^{n_k \times n_k}$ is the diagonal matrix with diagonal entries $W_{gii}^{(k)} = \frac{\mu_{ig}^{(k)}}{1+\mu_{ig}^{(k)}\alpha_g}$ fo $1 \leq i \leq n_k$. $\alpha_g$ is the dispersion estimate of the gene and $\mu_{ig}^{(k)}$ is the expected value of the gene $g$ for sample [illegible] for parameter $\beta_g$, that is $\gamma_g^{(k)} \exp(X_i^{(k)} \cdot \beta_g)$. | Each center | Server |
| | local_nll | nparray | $(1, G_{\texttt{act}})$ | The nll of the negative binomial GLM with fixed dispersion at the $\beta_g$ parameter, computed on the local data, fo each gene $g$. | Each center | Server |
| | beta | nparray | $(G_{\texttt{nz}}, p)$ | For evert gene $g$, the inital value of the $\beta_g$ parameter fo the ProxQuasiNewton algorithm (PQN), which is eithe i) the value computed by IRLS if IRLS has converged o that gene (in that case, the gene will not be optimized) or ii) the initial value of the $\beta_g$ parameter for the IRLS algorithm, if IRLS has not converged on that gene. | Each center | Server |
| | PQN_diverged_mask | nparray | $(G_{\texttt{nz}},)$ | A boolean array indicating which non zero genes have caused the Proximal Quasi-Newton algorithm to diverge Initialized to `False`. | Each center | Server |
| 19 | round_number_PQN | int | | The current round number of the Proximal Quasi-Newto algorithm, incremented by 1. | Server | Center |
| | irls_diverged_mask | nparray | $(G_{\texttt{nz}},)$ | A boolean array indicating which non zero genes have caused the IRLS algorithm to diverge. Passed on without modification. | Server | Center |
| | global_reg_nll | nparray | $(G_{\texttt{nz}},)$ | The regularized negative log likelihood of the negative binomial GLM with fixed dispersion at the $\beta_g$ paramete computed as the sum of the local negative log likelihood with $L^2$ regularization with $\lambda = 10^{-6}$. | Server | Center |
| | newton_decrement_on_mask | nparray | $(G_{\texttt{act}},)$ | The newton decrement on the active genes. It is computed as $\nu_g = \Delta_g \cdot \nabla_\beta \mathcal{L}^\lambda(\beta_g)$ where $\Delta_g$ is the ascen direction on the active genes and $\mathcal{L}^\lambda$ is the regularize log likelihood. | Server | Center |
| | beta | nparray | $(G_{\texttt{nz}}, p)$ | For each gene $g$, the log fold change $\beta_g$, which remains unchanged during the first iteration of the Proximal Quasi-Newton algorithm. | Server | Center |
| | PQN_diverged_mask | nparray | $(G_{\texttt{nz}},)$ | A boolean array indicating which non zero genes have caused the Proximal Quasi-Newton algorithm to diverge Passed on without modification. | Server | Center |
| | ascent_direction_on_mask | nparray | $(G_{\texttt{act}}, p)$ | The ascent direction on the active genes. To compute the ascent direction, we first compute the global Fishe information matrix for each gene $g$, $n\mathcal{I}_g = \sum_k n_k \mathcal{I}_g^{(k)}$ and the global gradient $\nabla_\beta \mathcal{L}(\beta_g) = \sum_k \nabla_\beta \mathcal{L}^{(k)}(\beta_g)$ From these two quantities, we build the Fisher information and gradients of the regularized negative log likelihood, with $L^2$ regularization $\lambda = 10^{-6}$. We add anothe regularization term to the Fisher matrix which depend on the iteration number. The goal is for the ascent direction to be close to the gradient for small iterations and to be close to the real natural gradient descent update near the end of the optimization. The ascent direction is then computed as from the regularized Fishe and gradient (see main paper). Roughly, it is the solution of the linear system $n\ \mathcal{I}_g^\lambda \Delta_g = \nabla_\beta \mathcal{L}^\lambda(\beta_g)$, where certain components are dropped in to handle boundary conditions as the optimization is constrained to values o $\beta_g$ in $[-\texttt{max\_beta}, \texttt{max\_beta}]$. | Server | Center |
| | PQN_mask | nparray | $(G_{\texttt{nz}},)$ | A boolean array indicating which non zero genes are currently being optimized by the Proximal Quasi-Newton algorithm (active genes), passed on without modification. | Server | Center |
| 20 | ascent_direction_on_mask | nparray | $(G_{\texttt{act}}, p)$ | The ascent direction on the active genes, passed on without modification. | Each center | Server |
| | beta | nparray | $(G_{\texttt{nz}}, p)$ | For each gene $g$, the log fold change $\beta_g$, which is simply passed on. | Each center | Server |
| | local_nll | nparray | $(N_{\texttt{ls}}, G_{\texttt{act}})$ | The nll of the negative binomial GLM with fixed dispersion at the $\beta_g$ parameter, computed on the local data, fo each gene $g$ and for each potential next step $\beta_g - \delta\Delta_g$ for $\delta \in \{1, 1/2, 1/4, ..., 1/2^{N_{\texttt{ls}}-1}\}$. | Each center | Server |
| | local_fisher | nparray | $(N_{\texttt{ls}}, G_{\texttt{act}}, p, p$ | For each potential step size $\delta \in \{1, 1/2, 1/4, ..., 1/2^{N_{\texttt{ls}}-1}\}$, and each gene $g$, the Fisher information matrix of the negative binomia GLM with fixed dispersion computed at $\beta_g - \delta\Delta_g$ Formally, $n_k \mathcal{I}_{\delta,g}^{(k)} = (X^{(k)})^\top W_{\delta,g}^{(k)} X^{(k)}$ where $W_{\delta,g}^{(k)} \in \mathbb{R}^{n_k \times n_k}$ is the diagonal matrix with diagonal entries $W_{\delta,gii}^{(k)} = \frac{\mu_{\delta,ig}^{(k)}}{1+\mu_{\delta,ig}^{(k)}\alpha_g}$ for $1 \leq i \leq n_k$. $\alpha_g$ is the dispersion estimate of the gene and $\mu_{\delta,ig}^{(k)}$ is the expected value of the gene $g$ for sample $i$ for paramete $\beta_g - \delta\Delta_g$, that is $\gamma_g^{(k)} \exp(X_i^{(k)} \cdot (\beta_g - \delta\Delta_g))$. | Each center | Server |
| | local_gradient | nparray | $(N_{\texttt{ls}}, G_{\texttt{act}}, p)$ | For each potential step size $\delta$ and each gene $g$, the gradient of the negative log likelihood of the negative binomia GLM with fixed dispersion at the $\beta_g - \delta\Delta_g$ parameter computed on the local data. | Each center | Server |
| | PQN_diverged_mask | nparray | $(G_{\texttt{nz}},)$ | A boolean array indicating which non zero genes have caused the Proximal Quasi-Newton algorithm to diverge Passed on without modification. | Each center | Server |
| | PQN_mask | nparray | $(G_{\texttt{nz}},)$ | A boolean array indicating which non zero genes are currently being optimized by the Proximal Quasi-Newton algorithm (active genes), passed on without modification. | Each center | Server |
| | global_reg_nll | nparray | $(G_{\texttt{nz}},)$ | The regularized negative log likelihood of the negative binomial GLM with fixed dispersion at the $\beta_g$ parameter simply passed on. | Each center | Server |
| | newton_decrement_on_mask | nparray | $(G_{\texttt{act}},)$ | The newton decrement on the active genes, passed o without modification. | Each center | Server |
| | round_number_PQN | int | | The current round number of the Proximal Quasi-Newto algorithm, passed on without modification. | Each center | Server |
| | irls_diverged_mask | nparray | $(G_{\texttt{nz}},)$ | A boolean array indicating which non zero genes have caused the IRLS algorithm to diverge. Passed on without modification. | Each center | Server |

| | | | | | | |
|---|---|---|---|---|---|---|
| 21 | irls_diverged_mask | nparray | $(G_{\texttt{nz}},)$ | A boolean array indicating which non zero genes have caused the IRLS algorithm to diverge. Passed on without modification. | Server | Center |
| | global_reg_nll | nparray | $(G_{\texttt{nz}},)$ | The regularized negative log likelihood of the negative binomial GLM with fixed dispersion at the $\beta_g$ parameter computed as the sum of the local negative log likelihoods with $L^2$ regularization with $\lambda = 10^{-6}$. | Server | Center |
| | round_number_PQN | int | | The current round number of the Proximal Quasi-Newton algorithm, incremented by 1. | Server | Center |
| | newton_decrement_on_mask | nparray | $(G_{\texttt{act}},)$ | The newton decrement on the new active genes $g$, computed at the new $\beta_g$ iterate. For more details, see description of 19. | Server | Center |
| | beta | nparray | $(G_{\texttt{nz}}, p)$ | For each gene $g$, the log fold change $\beta_g$. This value has been updated by the Proximal Quasi-Newton algorithm for all active genes where an admissible step size was found using the Armijo condition (see equation 2.5 of [1]) | Server | Center |
| | PQN_diverged_mask | nparray | $(G_{\texttt{nz}},)$ | A boolean array indicating which non zero genes have caused the Proximal Quasi-Newton algorithm to diverge. A gene $g$ is considered to have diverged if for all step sizes $\delta \in \{1, 1/2, 1/4, ..., 1/2^{N_{\texttt{ls}}-1}\}$, and for the computed ascent direction $\Delta_g$, the Armijo condition is not satisfied, i.e., if $\mathcal{L}^\lambda(\beta_g) - \mathcal{L}^\lambda(\beta_g - \delta\Delta_g) \leq \delta \; \texttt{PQN\_c1} \; \nabla_\beta \, \mathcal{L}^\lambda(\beta_g) \cdot \Delta_g$ where $\mathcal{L}^\lambda$ is the regularized negative log likelihood, `PQN_c1` is the Armijo parameter which can be set by the user (default is $10^{-4}$), and $\nabla_\beta \, \mathcal{L}^\lambda(\beta_g)$ is the gradient of the regularized negative log likelihood. | Server | Center |
| | PQN_mask | nparray | $(G_{\texttt{nz}},)$ | A boolean array indicating which non zero genes are currently being optimized by the Proximal Quasi-Newton algorithm (active genes). Genes that had converged with the relative error criterion or diverged (no stepsize satisfying the Armijo condition) have set to `False`. | Server | Center |
| | ascent_direction_on_mask | nparray | $(G_{\texttt{act}}, p)$ | The ascent direction on the new active genes $g$, computed at the new $\beta_g$ iterate. | Server | Center |
| 22 | global_reg_nll | nparray | $(G_{\texttt{nz}},)$ | The regularized negative log likelihood of the negative binomial GLM with fixed dispersion at the $\beta_g$ parameter simply passed on. | Each center | Server |
| | irls_diverged_mask | nparray | $(G_{\texttt{nz}},)$ | A boolean array indicating which non zero genes have caused the IRLS algorithm to diverge. Passed on without modification. | Each center | Server |
| | round_number_PQN | int | | The current round number of the Proximal Quasi-Newton algorithm, passed on without modification. | Each center | Server |
| | newton_decrement_on_mask | nparray | $(G_{\texttt{act}},)$ | The newton decrement on the active genes, passed on without modification. | Each center | Server |
| | ascent_direction_on_mask | nparray | $(G_{\texttt{act}}, p)$ | The ascent direction on the active genes, passed on without modification. | Each center | Server |
| | PQN_mask | nparray | $(G_{\texttt{nz}},)$ | A boolean array indicating which non zero genes are currently being optimized by the Proximal Quasi-Newton algorithm (active genes), passed on without modification. | Each center | Server |
| | local_gradient | nparray | $(N_{\texttt{ls}}, G_{\texttt{act}}, p)$ | For each potential step size $\delta$ and each gene $g$, the gradient of the negative log likelihood of the negative binomial GLM with fixed dispersion at the $\beta_g - \delta\Delta_g$ parameter computed on the local data. | Each center | Server |
| | local_fisher | nparray | $(N_{\texttt{ls}}, G_{\texttt{act}}, p, p)$ | For each potential step size $\delta \in \{1, 1/2, 1/4, ..., 1/2^{N_{\texttt{ls}}-1}\}$, and each gene $g$, the Fisher information matrix of the negative binomial GLM with fixed dispersion computed at $\beta_g - \delta\Delta_g$. Formally, $n_k\mathcal{I}^{(k)}_{\delta,g} = (X^{(k)})^\top W^{(k)}_{\delta,g} X^{(k)}$ where $W^{(k)}_{\delta,g} \in \mathbb{R}^{n_k \times n_k}$ is the diagonal matrix with diagonal entries $W^{(k)}_{\delta,gii} = \frac{\mu^{(k)}_{\delta,ig}}{1+\mu^{(k)}_{\delta,ig}\alpha_g}$ for $1 \leq i \leq n_k$. $\alpha_g$ is the dispersion estimate of the gene and $\mu^{(k)}_{\delta,ig}$ is the expected value of the gene $g$ for sample $i$ for parameter $\beta_g - \delta\Delta_g$, that is $\gamma^{(k)}_g \exp(X^{(k)}_i \cdot (\beta_g - \delta\Delta_g))$. | Each center | Server |
| | local_nll | nparray | $(N_{\texttt{ls}}, G_{\texttt{act}})$ | The nll of the negative binomial GLM with fixed dispersion at the $\beta_g$ parameter, computed on the local data, for each gene $g$ and for each potential next step $\beta_g - \delta\Delta_g$ for $\delta \in \{1, 1/2, 1/4, ..., 1/2^{N_{\texttt{ls}}-1}\}$. | Each center | Server |
| | beta | nparray | $(G_{\texttt{nz}}, p)$ | For each gene $g$, the log fold change $\beta_g$, which is simply passed on. | Each center | Server |
| | PQN_diverged_mask | nparray | $(G_{\texttt{nz}},)$ | A boolean array indicating which non zero genes have caused the Proximal Quasi-Newton algorithm to diverge. Passed on without modification. | Each center | Server |
| 23 | beta | nparray | $(G_{\texttt{nz}}, p)$ | For each gene $g$, the log fold change $\beta_g$. This value has been updated by the Proximal Quasi-Newton algorithm for all active genes where an admissible step size was found using the Armijo condition (see equation 2.5 of [1]) | Server | Center |
| | PQN_diverged_mask | nparray | $(G_{\texttt{nz}},)$ | A boolean array indicating which non zero genes have caused the Proximal Quasi-Newton algorithm to diverge. As this is the last step of the optimization, a gene $g$ is considered to have diverged not only if the Armijo condition is never satisfied, but also if the convergence criterion is not met. A gene is said to converge if the relative difference between two successive iterates is smaller than a threshold value `PQN_ftol`, i.e., $\lvert \mathcal{L}^\lambda(\beta_g) - \mathcal{L}^\lambda(\beta_{g,\text{prev}}) \rvert \leq \texttt{PQN\_ftol} \max(\mathcal{L}^\lambda(\beta_g), \mathcal{L}^\lambda(\beta_{g,\text{prev}}), 1)$. | Server | Center |
| | irls_diverged_mask | nparray | $(G_{\texttt{nz}},)$ | A boolean array indicating which non zero genes have caused the IRLS algorithm to diverge. Passed on without modification. | Server | Center |
| 25 | unique_counts | Series | $(p,)$ | A pandas series indexed by unique level combinations (all design factor levels that coexist) and containing the number of samples in each level combination. In the single factor case the index will simply be the unique levels of the factor. | Each center | Server |
| 26 | counts_by_lvl | Series | $(L,)$ | A pandas series indexed by the unique levels of the design factors across all centers, and whose values are the number of samples in each level combination. | Server | Center |
| 29 | non_zero | nparray | $(G,)$ | A boolean array indicating which genes have non-zero counts in at least one center. | Each center | Server |
| | grid | nparray | $(G, N_{\texttt{gs}})$ | The logarithmic grid between `min_disp` and $\max(\texttt{max\_disp}, n)$ of size $N_{\texttt{gs}}$. | Each center | Server |
| | max_disp | int | | The effective maximum value of the dispersion parameter $\max(\texttt{max\_disp}, n)$. | Each center | Server |

| | | | | | | |
|---|---|---|---|---|---|---|
| | nll | nparray | $(G, N_{\texttt{gs}})$ | For all genes $g$ and all dispersion values $\alpha_g$ in the logarithmic grid between min_disp $\max(\texttt{max\_disp}, n)$ of size $N_{\texttt{gs}}$ the negative log likelihood of the negative binomial GLM with fixed mean parameters $\mu_{ig}^{(k)}$ over the set of loca samples and dispersion parameter $\alpha_g$. If the numbe of unique levels of all factors is equal to the numbe of parameters $p$, then $\mu_{ig}^{(k)}$ is defined as nan for zerc genes, and as the minimum of $\gamma_i^{(k)}$ $(X^{(k)})^\top G^{-1}\Phi_g$ anc min_mu := 0.5 (for the definition of $G$ and $\Phi_g$, see step 8). Otherwise, $\mu_{ig}^{(k)}$ is defined as $\gamma_i^{(k)}$ $\exp(X_i^{(k)} \cdot \beta_g)$ or non-zero genes, where $\beta_g$ is the log fold change result o the IRLS algorithm, and as nan on zero genes. | Each center | Server |
| | CR_summand | nparray | $(G, N_{\texttt{gs}}, p, p)$ | For all genes $g$ and all dispersion values $\alpha_g$ in the logarithmic grid between min_disp $\max(\texttt{max\_disp}, n)$ of size $N_{\texttt{gs}}$ the matrix of the Cox-Reid regularization term pertaining to the samples in the center $k$. This matrix is computec as $\mathrm{CR}_g^{(k)} = (X^{(k)})^\top W_g^{(k)} X^{(k)}$ where $W_g^{(k)} \in \mathbb{R}^{n_k \times n_k}$ is the diagonal matrix with diagonal entries $\frac{\mu_{ig}^{(k)}}{1+\mu_{ig}^{(k)}\alpha_g}$ for $1 \leq i \leq n_k$. $\mu_{ig}^{(k)}$ is the expected value of the gene for sample $i$, whose expression can be found in the description of the nll variable. | Each center | Server |
| 30 | genewise_dispersions | nparray | $(G,)$ | For each gene $g$, the current estimate of the dispersior parameter $\alpha_g$. This estimate is computed by first computing the global nll (summing all local nlls) as well as the global Cox-Reid regularization term, which is half the log determinant of the sum of the local Cox-Reid matrices. The Cox-Reid regularized nll per gene and per dispersion in the grid is obtained by summing the regularization term and the nll. Finally, for every gene, the dispersion parameter is estimated by taking the minizer of this regularized nll on the grid of size $N_{\texttt{gs}}$. | Server | Center |
| | lower_log_bounds | nparray | $(G,)$ | For each gene $g$, the maximum of the log of the min dispersion and $\alpha_g - \delta$ where $\delta$ is the current mesh size of the grid and $\alpha_g$ is the current dispersion estimate. This value will be used as a lower bound for the next grid search for the dispersion parameter. | Server | Center |
| | upper_log_bounds | nparray | $(G,)$ | For each gene $g$, the minimum of the log of the max dispersion and $\alpha_g + \delta$ where $\delta$ is the current mesh size of the grid and $\alpha_g$ is the current dispersion estimate. This value will be used as an upper bound for the next gric search for the dispersion parameter. | Server | Center |
| 31 | nll | nparray | $(G, N_{\texttt{gs}})$ | For all genes $g$ and all dispersion values $\alpha_g$ in the logarithmic grid between the exponential of the lower log bound and of the upper log bound of size $N_{\texttt{gs}}$, the negative log likelihood of the negative binomial GLM with fixed mean parameters $\mu_{ig}^{(k)}$ over the set of local samples and dispersion parameter $\alpha_g$. If the number of unique levels of all factors is equal to the number of parameters $p$, then $\mu_{ig}^{(k)}$ is defined as nan for zero genes, and as the minimum of $\gamma_i^{(k)}$ $(X^{(k)})^\top G^{-1}\Phi_g$ and min_mu := 0.5 (for the definition of $G$ and $\Phi_g$, see step 8). Otherwise $\mu_{ig}^{(k)}$ is defined as $\gamma_i^{(k)}$ $\exp(X_i^{(k)} \cdot \beta_g)$ on non-zero genes where $\beta_g$ is the log fold change result of the IRLS algorithm, and as nan on zero genes. | Each center | Server |
| | CR_summand | nparray | $(G, N_{\texttt{gs}}, p, p)$ | For all genes $g$ and all dispersion values $\alpha_g$ in the logarithmic grid between the exponential of the lower log bound and of the upper log bound of size $N_{\texttt{gs}}$, the matrix of the Cox-Reid regularization term pertaining to the samples in the center $k$. This matrix is computed as $\mathrm{CR}_g^{(k)} = (X^{(k)})^\top W_g^{(k)} X^{(k)}$ where $W_g^{(k)} \in \mathbb{R}^{n_k \times n_k}$ is the diagonal matrix with diagonal entries $\frac{\mu_{ig}^{(k)}}{1+\mu_{ig}^{(k)}\alpha_g}$ for $1 \leq i \leq n_k$. $\mu_{ig}^{(k)}$ is the expected value of the gene for sample $i$, whose expression can be found in the description of the nll variable. | Each center | Server |
| | grid | nparray | $(G, N_{\texttt{gs}})$ | The logarithmic grid between the exponential of the lower log bound and of the upper log bound of size $N_{\texttt{gs}}$. | Each center | Server |
| | max_disp | int | | The effective maximum value of the dispersion parameter $\max(\texttt{max\_disp}, n)$. | Each center | Server |
| | non_zero | nparray | $(G,)$ | A boolean array indicating which genes have non-zero counts in at least one center. | Each center | Server |
| 32 | genewise_dispersions | nparray | $(G,)$ | For each gene $g$, the current estimate of the dispersior parameter $\alpha_g$. This estimate is computed by first computing the global nll (summing all local nlls) as well as the global Cox-Reid regularization term, which is half the log determinant of the sum of the local Cox-Reid matrices. The Cox-Reid regularized nll per gene and per dispersion in the grid is obtained by summing the regularization term and the nll. Finally, for every gene, the dispersion parameter is estimated by taking the minizer of this regularized nll on the grid of size $N_{\texttt{gs}}$. | Server | Center |
| | lower_log_bounds | nparray | $(G,)$ | For each gene $g$, the maximum of the log of the min dispersion and $\alpha_g - \delta$ where $\delta$ is the current mesh size of the grid and $\alpha_g$ is the current dispersion estimate. This value will be used as a lower bound for the next grid search for the dispersion parameter. | Server | Center |
| | upper_log_bounds | nparray | $(G,)$ | For each gene $g$, the minimum of the log of the max dispersion and $\alpha_g + \delta$ where $\delta$ is the current mesh size of the grid and $\alpha_g$ is the current dispersion estimate. This value will be used as an upper bound for the next gric search for the dispersion parameter. | Server | Center |
| 33 | n_params | int | | The number of parameters in the design matrix, i.e., its number of columns. | Each center | Server |
| | genewise_dispersions | nparray | $(G,)$ | For each gene $g$, the current estimate of the dispersior parameter $\alpha_g$, as computed by the grid search. | Each center | Server |
| | non_zero | nparray | $(G,)$ | A boolean array indicating which genes have non-zero counts in at least one center. | Each center | Server |
| | n_obs | int | | The number of samples in the center. | Each center | Server |
| | mean_normed_counts | nparray | $(G,)$ | For each gene, the mean of the local normed counts, i.e. $\overline{Z}_g^{(k)} = \frac{1}{n_k}\sum_{i=1}^{n_k} Z_{ig}^{(k)}$. | Each center | Server |
| 34 | prior_disp_var | float | $()$ | A prior on the variance of the log dispersions around the log trend curve $\sigma^2_{\texttt{trend}}$, estimated as the maximum between 0.25 and $\texttt{\_squared\_log\_res} - \psi_1((n-p)/2)$, where $\psi_1(f/2)$ is the variance of the log of a $\chi^2_f$ distribution. For more details, see [2]. | Server | Center |

| | | | | | | |
|---|---|---|---|---|---|---|
| | _squared_logres | float | () | The squared mean absolute deviation of the difference between the log of the genewise dispersions and the log of the fitted dispersions, restricted to the non-zero genes whose gene-wise dispersions are above $100 \times$ min_disp. The mean absolute deviation estimate is defined as the median of the absolute difference between the log residual and its median, scaled by the percent point function of the normal distribution at 0.75. | Server | Center |
| | trend_coeffs | nparray | $(2,)$ | The coefficients of the trend curve fitted to the dispersions. We model the dispersions $\alpha_g$ of the gene $g$ as a sample from an exponential distribution whose mean $\alpha_{\texttt{trend}}(\overline{Z}_g)$ is a function of $\overline{Z}_g$ parametrized by two coefficients $\alpha_0$ and $\mathrm{a}_1$; $\alpha_{\texttt{trend}}(\overline{Z}_g) = \alpha_0 + \frac{\mathrm{a}_1}{\overline{Z}_g}$. $\alpha_{\texttt{trend}}$ is called the trend curve. The coefficients are $(\alpha_0, \mathrm{a}_1)$ and are obtained by starting with the set of non zero genes and by iteratively i) minimizing the negative log likelihood of the exponential distribution on the set of genes with LBFGS-B and ii) removing the genes where the ratio between the dispersion $\alpha_g$ and the trend curve $\alpha_{\texttt{trend}}(\overline{Z}_g)$ is above 15 or below $10^{-4}$ (we consider these genes as outliers to this model), until the set of genes is stable (see [2] for more details). | Server | Center |
| | fitted_dispersions | nparray | $(G,)$ | For each gene $g$ with zero counts across all centers, nan. For each non-zero gene $g$, either $\alpha_{\texttt{trend}}(\overline{Z}_g)$ if the disp_function_type is "parametric" (i.e., the fitting of the parameters has converged), or mean_disp otherwise (i.e., if the disp_function_type is "mean"). Denoted with $\alpha_g^{\texttt{trend}}$. | Server | Center |
| | disp_function_type | str | | The type of the dispersion function used to model the dispersions. It can be either "parametric", if the iterative scheme to fit the trend curve has converged, the LBFGS-B method used to fit the parameters has converged, and the coefficienst of the trend curve are non-negative; or "mean" otherwise. | Server | Center |
| | mean_disp | NoneType | | None if the dispersion function type is "parametric" and the trimmed mean of the genewise dispersions whose value is above $10 \times$ min_disp, with trimming proportion 0.001 otherwise. | Server | Center |
| 35 | non_zero | nparray | $(G,)$ | A boolean array indicating which genes have non-zero counts in at least one center. | Each center | Server |
| | max_disp | int | | The effective maximum value of the dispersion parameter $\max(\texttt{max\_disp}, n)$. | Each center | Server |
| | reg | nparray | $(G, N_{\texttt{gs}})$ | For all genes $g$ and all dispersion values $\alpha_g$ in the logarithmic grid between min_disp and $\max(\texttt{max\_disp}, n)$ of size $N_{\texttt{gs}}$, a regularization term that is equal to $0.5 \times (\log(\alpha_g) - \log(\alpha_g^{\texttt{trend}}))^2/\sigma^2_{\texttt{trend}}$, and which comes from the prior on the distribution of the log dispersions around the trend curve. | Each center | Server |
| | CR_summand | nparray | $(G, N_{\texttt{gs}}, p, p)$ | For all genes $g$ and all dispersion values $\alpha_g$ in the logarithmic grid between min_disp $\max(\texttt{max\_disp}, n)$ of size $N_{\texttt{gs}}$ the matrix of the Cox-Reid regularization term pertaining to the samples in the center $k$. This matrix is computed as $\mathrm{CR}_g^{(k)} = (X^{(k)})^\top W_g^{(k)} X^{(k)}$ where $W_g^{(k)} \in \mathbb{R}^{n_k \times n_k}$ is the diagonal matrix with diagonal entries $\frac{\mu_{ig}^{(k)}}{1+\mu_{ig}^{(k)}\alpha_g}$ for $1 \leq i \leq n_k$. $\mu_{ig}^{(k)}$ is the expected value of the gene for sample $i$, whose expression can be found in the description of the nll variable. | Each center | Server |
| | nll | nparray | $(G, N_{\texttt{gs}})$ | For all genes $g$ and all dispersion values $\alpha_g$ in the logarithmic grid between min_disp $\max(\texttt{max\_disp}, n)$ of size $N_{\texttt{gs}}$ the negative log likelihood of the negative binomial GLM with fixed mean parameters $\mu_{ig}^{(k)}$ over the set of local samples and dispersion parameter $\alpha_g$. If the number of unique levels of all factors is equal to the number of parameters $p$, then $\mu_{ig}^{(k)}$ is defined as nan for zero genes, and as the minimum of $\gamma_i^{(k)}$ $(X^{(k)})^\top G^{-1} \Phi_g$ and min_mu $:= 0.5$ (for the definition of $G$ and $\Phi_g$, see step 8). Otherwise, $\mu_{ig}^{(k)}$ is defined as $\gamma_i^{(k)}$ $\exp(X_i^{(k)} \cdot \beta_g)$ on non-zero genes, where $\beta_g$ is the log fold change result of the IRLS algorithm, and as nan on zero genes. | Each center | Server |
| | grid | nparray | $(G, N_{\texttt{gs}})$ | The logarithmic grid between min_disp and $\max(\texttt{max\_disp}, n)$ of size $N_{\texttt{gs}}$. | Each center | Server |
| 36 | MAP_dispersions | nparray | $(G,)$ | For each gene $g$, the current estimate of the MAP dispersion parameter $\alpha_g$. This estimate is computed by first computing the global nll (summing all local nlls) as well as the global Cox-Reid regularization term, which is half the log determinant of the sum of the local Cox-Reid matrices, and the regularization coming from the prior on the log dispersions around the trend curve. The Cox-Reid, prior regularized nll per gene and per dispersion in the grid is obtained by summing the regularization terms and the nll. Finally, for every gene, the MAP dispersion parameter is estimated by taking the minizer of this regularized nll on the grid of size $N_{\texttt{gs}}$. | Server | Center |
| | lower_log_bounds | nparray | $(G,)$ | For each gene $g$, the maximum of the log of the min dispersion and $\alpha_g - \delta$ where $\delta$ is the current mesh size of the grid and $\alpha_g$ is the current dispersion estimate. This value will be used as a lower bound for the next grid search for the dispersion parameter. | Server | Center |
| | upper_log_bounds | nparray | $(G,)$ | For each gene $g$, the minimum of the log of the max dispersion and $\alpha_g + \delta$ where $\delta$ is the current mesh size of the grid and $\alpha_g$ is the current dispersion estimate. This value will be used as an upper bound for the next grid search for the dispersion parameter. | Server | Center |
| 37 | reg | nparray | $(G, N_{\texttt{gs}})$ | For all genes $g$ and all dispersion values $\alpha_g$ in the logarithmic grid between the exponential of the lower log bound and of the upper log bound of size $N_{\texttt{gs}}$, a regularization term that is equal to $0.5 \times (\log(\alpha_g) - \log(\alpha_g^{\texttt{trend}}))^2/\sigma^2_{\texttt{trend}}$, and which comes from the prior on the distribution of the log dispersions around the trend curve. | Each center | Server |
| | non_zero | nparray | $(G,)$ | A boolean array indicating which genes have non-zero counts in at least one center. | Each center | Server |
| | max_disp | int | | The effective maximum value of the dispersion parameter $\max(\texttt{max\_disp}, n)$. | Each center | Server |
| | grid | nparray | $(G, N_{\texttt{gs}})$ | The logarithmic grid between the exponential of the lower log bound and of the upper log bound of size $N_{\texttt{gs}}$. | Each center | Server |

| | | | | | | |
|---|---|---|---|---|---|---|
| | nll | nparray | $(G, N_{\texttt{gs}})$ | For all genes $g$ and all dispersion values $\alpha_g$ in the logarithmic grid between the exponential of the lower log bound and of the upper log bound of size $N_{\texttt{gs}}$, the negative log likelihood of the negative binomial GLM with fixed mean parameters $\mu_{ig}^{(k)}$ over the set of local samples and dispersion parameter $\alpha_g$. If the number of unique levels of all factors is equal to the number of parameters $p$, then $\mu_{ig}^{(k)}$ is defined as `nan` for zero genes, and as the minimum of $\gamma_i^{(k)}\ (X^{(k)})^\top G^{-1}\Phi_g$ and `min_mu` $:= 0.5$ (for the definition of $G$ and $\Phi_g$, see step 8). Otherwise $\mu_{ig}^{(k)}$ is defined as $\gamma_i^{(k)}\ \exp(X_i^{(k)} \cdot \beta_g)$ on non-zero genes where $\beta_g$ is the log fold change result of the IRLS algorithm, and as `nan` on zero genes. | Each center | Server |
| | CR_summand | nparray | $(G, N_{\texttt{gs}}, p, p)$ | For all genes $g$ and all dispersion values $\alpha_g$ in the logarithmic grid between the exponential of the lower log bound and of the upper log bound of size $N_{\texttt{gs}}$, the matrix of the Cox-Reid regularization term pertaining to the samples in the center $k$. This matrix is computed as $\text{CR}_g^{(k)} = (X^{(k)})^\top W_g^{(k)} X^{(k)}$ where $W_g^{(k)} \in \mathbb{R}^{n_k \times n_k}$ is the diagonal matrix with diagonal entries $\frac{\mu_{ig}^{(k)}}{1+\mu_{ig}^{(k)}\alpha_g}$ for $1 \leq i \leq n_k$. $\mu_{ig}^{(k)}$ is the expected value of the gene for sample $i$, whose expression can be found in the description of the nll variable. | Each center | Server |
| 38 | MAP_dispersions | nparray | $(G, )$ | For each gene $g$, the current estimate of the MAP dispersion parameter $\alpha_g$. This estimate is computed by first computing the global nll (summing all local nlls) as well as the global Cox-Reid regularization term, which is half the log determinant of the sum of the local Cox-Reid matrices, and the regularization coming from the prior on the log dispersions around the trend curve. The Cox-Reid, prior regularized nll per gene and per dispersion in the grid is obtained by summing the regularization terms and the nll. Finally, for every gene, the MAP dispersion parameter is estimated by taking the minizer of this regularized nll on the grid of size $N_{\texttt{gs}}$. | Server | Center |
| | lower_log_bounds | nparray | $(G, )$ | For each gene $g$, the maximum of the log of the min dispersion and $\alpha_g - \delta$ where $\delta$ is the current mesh size of the grid and $\alpha_g$ is the current dispersion estimate. This value will be used as a lower bound for the next grid search for the dispersion parameter. | Server | Center |
| | upper_log_bounds | nparray | $(G, )$ | For each gene $g$, the minimum of the log of the max dispersion and $\alpha_g + \delta$ where $\delta$ is the current mesh size of the grid and $\alpha_g$ is the current dispersion estimate. This value will be used as an upper bound for the next grid search for the dispersion parameter. | Server | Center |
| 40 | global_gram_matrix | nparray | $(p, p)$ | The sum of the local Gramm matrix across all centers $G = \sum_k G^{(k)}$. | Each center | Server |
| | local_log_features | nparray | $(G_{\texttt{nz}}, p)$ | $\Phi_k = (X^{(k)})^\top \log(Z^{(k)} + 0.1)$. | Each center | Server |
| | gram_full_rank | bool | $()$ | A boolean indicating whether the global Gramm matrix $G$ is full rank. | Each center | Server |
| | n_non_zero_genes | int | | The number of genes with non-zero counts in at least one center. | Each center | Server |
| 41 | beta | nparray | $(G_{\texttt{nz}}, p)$ | The initial value of the $\beta$ parameter. If the Gramm matrix is full rank, is set to $\beta_g = G^{-1}(\sum_k \Phi_g^{(k)})$ for all genes $g$. Otherwise, it is set as a weighed average of the normed log means, i.e., $\beta_g = \sum_k \frac{n_k}{n}\ \overline{\log(Z)}_g^{(k)}$, where $\overline{\log(Z)}_g^{(k)} = \frac{1}{n_k}\sum_{i=1}^{n_k}\log(Z_{ig}^{(k)})$ is computed locally. | Server | Center |
| | irls_diverged_mask | nparray | $(G_{\texttt{nz}}, )$ | A boolean array indicating which non zero genes have caused the IRLS algorithm to diverge. Initialized to `False`. | Server | Center |
| | irls_mask | nparray | $(G_{\texttt{nz}}, )$ | A boolean array indicating which non zero genes are currently being optimized by the IRLS algorithm. Initialized to `True`. | Server | Center |
| | global_nll | nparray | $(G_{\texttt{nz}}, )$ | For each gene $g$, the negative log likelihood of the negative binomial GLM, initialized at 1000 for all genes. | Server | Center |
| | round_number_irls | int | | The current round number of the IRLS algorithm. Initialized at 0. | Server | Center |
| 42 | irls_mask | nparray | $(G_{\texttt{nz}}, )$ | A boolean array indicating which non zero genes are currently being optimized by the IRLS algorithm. Simply passed on. | Each center | Server |
| | round_number_irls | int | | The current round number of the IRLS algorithm. Simply passed on. | Each center | Server |
| | global_nll | nparray | $(G_{\texttt{nz}}, )$ | For each gene $g$, the negative log likelihood at the $\beta_g$ parameter. Simply passed on. | Each center | Server |
| | irls_diverged_mask | nparray | $(G_{\texttt{nz}}, )$ | A boolean array indicating which non zero genes have caused the IRLS algorithm to diverge. Simply passed on. | Each center | Server |
| | irls_gene_names | Index | $(G_{\texttt{act}}, )$ | The gene names of the active genes. | Each center | Server |
| | local_features | nparray | $(G_{\texttt{act}}, p)$ | For each active gene $g$, the projected features $(X^{(k)})^\top W_g^{(k)} \phi_g^{(k)}$, where $\phi_{ig}^{(k)} = \log\left(\frac{\mu_{ig}^{(k)}}{\gamma_i}\right) + \frac{Y_{ig}^{(k)} - \mu_{ig}^{(k)}}{\mu_{ig}^{(k)}}$ and $\phi_g^{(k)}$ is the vector of $(\phi_{ig}^{(k)})_{1 \leq i \leq n_k}$. | Each center | Server |
| | local_hat_matrix | nparray | $(G_{\texttt{act}}, p, p)$ | For each gene $g$, the hat matrix $H_g^{(k)} = (X^{(k)})^\top W_g^{(k)} X^{(k)}$ where $W_g^{(k)} \in \mathbb{R}^{n_k \times n_k}$ is the diagonal matrix with diagonal entries $\frac{\mu_{ig}^{(k)}}{1+\mu_{ig}^{(k)}\alpha_g}$ for $1 \leq i \leq n_k$. $\alpha_g$ is the dispersion estimate of the gene and $\mu_{ig}^{(k)}$ is the expected value of the gene for sample $i$ for parameter $\beta$, that is $\gamma_g^{(k)} \exp(X_i^{(k)} \cdot \beta_g)$. | Each center | Server |
| | local_nll | nparray | $(G_{\texttt{act}}, )$ | For each gene $g$, the negative log likelihood of the negative binomial GLM with fixed dispersion at the $\beta_g$ parameter, computed on the local data. | Each center | Server |
| | beta | nparray | $(G_{\texttt{nz}}, p)$ | The log fold change $\beta$ parameter. Not modified. | Each center | Server |
| 43 | beta | nparray | $(G_{\texttt{nz}}, p)$ | The updated value of the $\beta$ parameter after applying the IRLS update for the active genes. At the last step, is not updated. | Server | Center |
| | irls_diverged_mask | nparray | $(G_{\texttt{nz}}, )$ | A boolean array indicating which non zero genes have caused the IRLS algorithm to diverge. A gene $g$ is considered to have diverged if any component of $\beta_g \in \mathbb{R}^p$ has absolute value above the threshold `max_beta`. | Server | Center |

| | | | | | | |
|---|---|---|---|---|---|---|
| | PQN_diverged_mask | nparray | $(G_{\mathtt{nz}},)$ | A boolean array indicating which non zero genes have caused the Proximal Quasi-Newton algorithm to diverge. As this is the last step of the optimization, a gene $g$ is considered to have diverged not only if the Armijo condition is never satisfied, but also if the convergence criterion is not met. A gene is said to converge if the relative difference between two successive iterates is smaller than a threshold value PQN_ftol, i.e., $\mid \mathcal{L}^{\lambda}(\beta_g) - \mathcal{L}^{\lambda}(\beta_{g,\text{prev}})\mid \leq$ PQN_ftol $\max(\mathcal{L}^{\lambda}(\beta_g), \mathcal{L}^{\lambda}(\beta_{g,\text{prev}}), 1)$. | Server | Center |
| | irls_diverged_mask | nparray | $(G_{\mathtt{nz}},)$ | A boolean array indicating which non zero genes have caused the IRLS algorithm to diverge. Passed on without modification. | Server | Center |
| 51 | use_lvl | bool | () | Whether or not to use the levels of the design matrix to compute the trimmed mean. If use_lvl is True, then one trimmed mean of the normed counts will be computed for each gene and for each level of the design matrix (that is the trimmed mean will be performed on the samples corresponding to the same level of the design matrix across all centers). If use_lvl is False, then one trimmed mean of the normed counts will be computed for each gene and for all samples. This parameter is set to True if there is at least one level of the design matrix with more than 3 replicates across all centers, and to False otherwise. | Each center | Server |
| | $l \in \mathcal{L}_{\geq 3}$ | dict | | Let $\mathcal{L}_{\geq 3}$ bet the set of levels $1 \leq l \leq L$ of the design matrix with at least 3 replicates across all centers. Let $\mathcal{I}_{k,l}$ denote the set of sample indices $1 \leq i \leq n_k$ in center $k$ whose line in the design corresponds to level $l$. Dictionary with two keys.<br>"max_values", which is a numpy array of shape $G$ containing, for each gene $g$, the maximum value of the center normed counts for $g$ on samples in $\mathcal{I}_{k,l}$, denoted with $(Z^{\mathtt{max}}_{g,l})^{(k)}$.<br>"min_values", which is a numpy array of shape $G$ containing, for each gene $g$, the minimum value of the center normed counts for $g$ on samples in $\mathcal{I}_{k,l}$, denoted with $(Z^{\mathtt{min}}_{g,l})^{(k)}$. | Each center | Server |
| 52 | use_lvl | bool | () | Whether or not to use the levels of the design matrix to compute the trimmed mean. Passed on without modification. | Server | Center |
| | $l \in \mathcal{L}_{\geq 3}$ | dict | | Dictionary with two keys.<br>"upper_bounds_thresholds" is a numpy array of shape $(G, 2)$. For each gene $g$ and for each ratio $r \in \{r_{\mathtt{trim}}, 1 - r_{\mathtt{trim}}\}$, will contain an upper bound of the $r$-quantile value of the normed counts of gene $g$ across samples in level $l$. Initialized for both ratios as the maximum of the normed counts of gene $g$ across all samples in level $l$, computed as $\max_k (Z^{\mathtt{max}}_{g,l})^{(k)}$.<br>"lower_bounds_thresholds" is a numpy array of shape $(G, 2)$. For each gene $g$ and for each ratio $r \in \{r_{\mathtt{trim}}, 1 - r_{\mathtt{trim}}\}$, will contain a lower bound of the $r$-quantile value of the normed counts of gene $g$ across samples in level $l$. Initialized for both ratios as the minimum of the normed counts of gene $g$ across all samples in level $l$, computed as $\min_k (Z^{\mathtt{min}}_{g,l})^{(k)}$. | Server | Center |
| 53 | use_lvl | bool | () | Whether or not to use the levels of the design matrix to compute the trimmed mean. Passed on without modification. | Each center | Server |
| | $l \in \mathcal{L}_{\geq 3}$ | dict | | Dictionary with the following keys.<br>"num_strictly_above" is a numpy array of shape $(G, 2)$. For each gene $g$ and each ratio $r$, contains the number of samples in the center and in the level whose normed counts value for gene $g$ is strictly above the average of the upper and lower bounds on the $r$-quantile value of gene $g$.<br>"upper_bounds_thresholds" is a numpy array of shape $(G, 2)$, simply passed on from the previous state.<br>"lower_bounds_thresholds" is a numpy array of shape $(G, 2)$ simply passed on from the previous state.<br>"n_samples" is the number of samples in level $l$ and center $k$.<br>"trim_ratio" is the trim ration, defining the upper and lower ratios whose quantile we want to compute. Denoted with $r_{\mathtt{trim}}$, equal to 0.125. | Each center | Server |
| 54 | use_lvl | bool | () | Whether or not to use the levels of the design matrix to compute the trimmed mean. Passed on without modification. | Server | Center |
| | $l \in \mathcal{L}_{\geq 3}$ | dict | | Dictionary with two keys.<br>"upper_bounds_thresholds" is a numpy array of shape $(G, 2)$. For each gene $g$ and for each ratio $r \in \{r_{\mathtt{trim}}, 1 - r_{\mathtt{trim}}\}$, contains an updated upper bound of the $r$-quantile value of the normed counts of gene $g$ across samples in level $l$. For a given gene $g$ and $r$-quantile, it is updated as either its previous value or the average of the previous upper and lower bounds across samples in level $l$, depending on the number of samples in the level whose normed counts value is strictly above this average. This number of samples can be computed from the previous shared states.<br>"lower_bounds_thresholds" is a numpy array of shape $(G, 2)$. For each gene $g$ and for each ratio $r \in \{r_{\mathtt{trim}}, 1 - r_{\mathtt{trim}}\}$, contains an updated lower bound of the $r$-quantile value of the normed counts of gene $g$ across samples in level $l$. For a given gene $g$ and $r$-quantile, it is updated as either its previous value or the average of the previous upper and lower bounds across samples in level $l$, depending on the number of samples in the level whose normed counts value is strictly above this average. This number of samples can be computed from the previous shared states. | Server | Center |
| 55 | use_lvl | bool | () | Whether or not to use the levels of the design matrix to compute the trimmed mean. Passed on without modification. | Each center | Server |

| | | | | | | |
|---|---|---|---|---|---|---|
| | $l \in \mathcal{L}_{\geq 3}$ | dict | | Dictionary with the following keys. "trimmed_local_sum" is a numpy array of shape $(G,)$. For each gene $g$ and each ration $r \in \{r_{\texttt{trim}}, 1 - r_{\texttt{trim}}\}$, we approximate the $r$-quantile by taking the average of the running upper and lower bounds. The trimmed local sum is computed as the sum of the normed counts across all samples in level $l$ and center $k$, whose value is strictly above the approximation of the $r_{\texttt{trim}}$-quantile and less or equal to the approximation of the $1 - r_{\texttt{trim}}$ quantile. "n_samples" is the number of samples in level $l$ and center $k$. "num_strictly_above" is a numpy array of shape $(G, 2)$. For each gene $g$ and each ratio $r$, contains the number of samples in the center and in the level whose normed counts value for gene $g$ is strictly above the approximation of the $r$-quantile. "upper_bounds_thresholds" is a numpy array of shape $(G, 2)$, simply passed on from the previous state. "lower_bounds_thresholds" is a numpy array of shape $(G, 2)$ simply passed on from the previous state. "trim_ratio", equal to 0.125. | Each center | Server |
| 56 | trimmed_mean_normed_counts | DataFrame | $(G, \lvert\mathcal{L}_{\geq 3}\rvert)$ | For each gene $g$ and each level $l \in \mathcal{L}_{\geq 3}$, the corresponding entry $\overline{Z}^{\texttt{trim}}_{g,l}$ is the approximation of the trimmed mean of the normed counts for gene $g$ and samples whose line in the design corresponds to level $l$ with trim ratio $r_{\texttt{trim}}$, computed by summing the trimmed_local_sums and dividing by the sum of the local n_samples for the corresponding gene and level. | Server | Center |
| 58 | $l \in \mathcal{L}_{\geq 3}$ | dict | | Let $\mathcal{L}_{\geq 3}$ bet the set of levels $1 \leq l \leq L$ of the design matrix with at least 3 replicates across all centers. Let $\mathcal{I}_{k,l}$ denote the set of sample indices $1 \leq i \leq n_k$ in center $k$ whose line in the design corresponds to level $l$. Let $R^{\texttt{trim}}_k$ be the matrix of square errors of size $(n, G)$ whose entries are $(Z^{(k)}_{ig} - \overline{Z}^{\texttt{trim}}_{g,l})^2$ if $i \in \mathcal{I}_{k,l}$ for some $l \in \mathcal{L}_{\geq 3}$ and nan otherwise, for $1 \leq i \leq n_k$ and $1 \leq g \leq G$. Dictionary with two keys. "max_values", which is a numpy array of shape $G$ containing, for each gene $g$, the maximum value of the center squared errors for $g$ on samples in $\mathcal{I}_{k,l}$, denoted with $R^{\texttt{max}}_{k,g,l}$. "min_values", which is a numpy array of shape $G$ containing, for each gene $g$, the minimum value of the center squared errors for $g$ on samples in $\mathcal{I}_{k,l}$, denoted with $R^{\texttt{min}}_{k,g,l}$. | Each center | Server |
| | use_lvl | bool | () | Whether or not to use the levels of the design matrix to compute the trimmed mean. If use_lvl is `True`, then one trimmed mean of the squared errors will be computed for each gene and for each level of the design matrix (that is the trimmed mean will be performed on the samples corresponding to the same level of the design matrix across all centers). If use_lvl is `False`, then one trimmed mean of the squared errors will be computed for each gene and for all samples. This parameter is set to `True` if there is at least one level of the design matrix with more than 3 replicates across all centers, and to `False` otherwise. | Each center | Server |
| 59 | $l \in \mathcal{L}_{\geq 3}$ | dict | | Dictionary with two keys. "upper_bounds_thresholds" is a numpy array of shape $(G, 2)$. For each gene $g$ and for each ratio $r \in \{r_{\texttt{trim}}, 1 - r_{\texttt{trim}}\}$, will contain an upper bound of the $r$-quantile value of the squared errors of gene $g$ across samples in level $l$. Initialized for both ratios as the maximum of the squared errors of gene $g$ across all samples in level $l$, computed as $\max_k R^{\texttt{max}}_{k,g,l}$. "lower_bounds_thresholds" is a numpy array of shape $(G, 2)$. For each gene $g$ and for each ratio $r \in \{r_{\texttt{trim}}, 1 - r_{\texttt{trim}}\}$, will contain a lower bound of the $r$-quantile value of the squared errors of gene $g$ across samples in level $l$. Initialized for both ratios as the minimum of the squared errors of gene $g$ across all samples in level $l$, computed as $\min_k R^{\texttt{min}}_{k,g,l}$. | Server | Center |
| | use_lvl | bool | () | Whether or not to use the levels of the design matrix to compute the trimmed mean. Passed on without modification. | Server | Center |
| 60 | use_lvl | bool | () | Whether or not to use the levels of the design matrix to compute the trimmed mean. Passed on without modification. | Each center | Server |
| | $l \in \mathcal{L}_{\geq 3}$ | dict | | Dictionary with the following keys. "num_strictly_above" is a numpy array of shape $(G, 2)$. For each gene $g$ and each ratio $r$, contains the number of samples in the center and in the level whose squared errors value for gene $g$ is strictly above the average of the upper and lower bounds on the $r$-quantile value of gene $g$. "upper_bounds_thresholds" is a numpy array of shape $(G, 2)$, simply passed on from the previous state. "lower_bounds_thresholds" is a numpy array of shape $(G, 2)$ simply passed on from the previous state. "n_samples" is the number of samples in level $l$ and center $k$. "trim_ratio" is the trim ration, defining the upper and lower ratios whose quantile we want to compute. Denoted with $r_{\texttt{trim}}$, equal to 0.125. | Each center | Server |
| 61 | use_lvl | bool | () | Whether or not to use the levels of the design matrix to compute the trimmed mean. Passed on without modification. | Server | Center |
| | $l \in \mathcal{L}_{\geq 3}$ | dict | | Dictionary with two keys. "upper_bounds_thresholds" is a numpy array of shape $(G, 2)$. For each gene $g$ and for each ratio $r \in \{r_{\texttt{trim}}, 1 - r_{\texttt{trim}}\}$, contains an updated upper bound of the $r$-quantile value of the squared errors of gene $g$ across samples in level $l$. For a given gene $g$ and $r$-quantile, it is updated as either its previous value or the average of the previous upper and lower bounds across samples in level $l$, depending on the number of samples in the level whose squared errors value is strictly above this average. This number of samples can be computed from the previous shared states. "lower_bounds_thresholds" is a numpy array of shape $(G, 2)$. For each gene $g$ and for each ratio $r \in \{r_{\texttt{trim}}, 1 - r_{\texttt{trim}}\}$, contains an updated lower bound of the $r$-quantile value of the squared errors of gene $g$ across samples in level $l$. For a given gene $g$ and $r$-quantile, it is updated as either its previous value or the average of the previous upper and lower bounds across samples in level $l$, depending on the number of samples in the level whose squared errors value is strictly above this average. This number of samples can be computed from the previous shared states. | Server | Center |

| | | | | | | |
|---|---|---|---|---|---|---|
| 62 | $l \in \mathcal{L}_{\geq 3}$ | dict | | Dictionary with the following keys. "trimmed_local_sum" is a numpy array of shape $(G, )$. Fo each gene $g$ and each ration $r \in \{r_{\texttt{trim}}, 1 - r_{\texttt{trim}}\}$, we approximate the $r$-quantile by taking the average of the running upper and lower bounds. The trimmed local sun is computed as the sum of the squared errors across al samples in level $l$ and center $k$, whose value is strictly above the approximation of the $r_{\texttt{trim}}$-quantile and less o equal to the approximation of the $1 - r_{\texttt{trim}}$ quantile. "n_samples" is the number of samples in level $l$ and center $k$. "num_strictly_above" is a numpy array of shape $(G, 2)$ For each gene $g$ and each ratio $r$, contains the number o samples in the center and in the level whose squared errors value for gene $g$ is strictly above the approximation of the $r$-quantile. "upper_bounds_thresholds" is a numpy array of shape $(G, 2)$, simply passed on from the previous state. "lower_bounds_thresholds" is a numpy array of shape $(G, 2)$ simply passed on from the previous state "trim_ratio", equal to 0.125. "scale", equal to 1.51. | Each center | Server |
| | use_lvl | bool | () | Whether or not to use the levels of the design matrix to compute the trimmed mean. Passed on without modification. | Each center | Server |
| 63 | varEst | nparray | $(G, )$ | For each gene $g$, the trimmed variance estimate of the normed counts, denoted with $V_g^{\texttt{trim}}$. If use_lvl is `True`, for each gene $g$, it is the maximum across all admissible levels of the trimmed mean of the squared error for the gene and level in question, with trim ratio $r_{\texttt{trim}}$, scaled (multiplied) by the scale factor 1.51. If use_level is `False`, the trimmed mean of the squared error scaled by 1.51 with trim ratio $r_{\texttt{trim}}$. | Server | Center |
| 64 | _skip_cooks | bool | | A boolean indicating whether to skip the computation of the intermediate quantities to compute the of the Cook's distance. This is set to `True` if the Cook's distance is stored in the local state (which is not the case by default due to memory issues). Otherwise, it is set to `False` (default behaviour). | Each center | Server |
| | varEst | nparray | $(G, )$ | For each gene $g$, the trimmed variance estimate of the normed counts, denoted with $V_g^{\texttt{trim}}$. This quantitiy is passed on from the previous shared state. | Each center | Server |
| | n_samples | int | | The number of samples in a center $n_k$ for each center $k$. | Each center | Server |
| | mean_normed_counts | nparray | $(G, )$ | For each gene, the mean of the local normed counts, i.e. $\overline{Z}_g^{(k)} = \frac{1}{n_k}\sum_{i=1}^{n_k} Z_{ig}^{(k)}$. | Each center | Server |
| | local_hat_matrix | nparray | $(G_{\texttt{nz}}, p, p)$ | For each gene $g$, the hat matrix $H_g^{(k)} = (X^{(k)})^\top W_g^{(k)} X^{(k)}$ where $W_g^{(k)} \in \mathbb{R}^{n_k \times n_k}$ is the diagonal matrix with diagonal entries $\frac{\mu_{ig}^{(k)}}{1+\mu_{ig}^{(k)}\alpha_g}$ for $1 \leq i \leq n_k$. $\alpha_g$ is the dispersion estimate of the gene and $\mu_{ig}^{(k)}$ is the expected value of the gene for sample $i$ for parameter $\beta$, that is $\gamma_g^{(k)} \exp(X_i^{(k)} \cdot \beta_g)$. | Each center | Server |
| 65 | global_hat_matrix_inv | nparray | $(G_{\texttt{nz}}, p, p)$ | For each gene $g$, we compute the global hat matrix as the sum of the local hat matrices, and its inverse. | Server | Center |
| | cooks_dispersions | nparray | $(G, )$ | For each gene $g$, a robust estimate of the dispersion parameter $\alpha_g^{\texttt{cooks}}$ computed from the trimmed variance estimate and the global mean of the normed counts. We compute this estimate as $\max((V_g^{\texttt{trim}} - \overline{Y}_g)/\overline{Y}_g^2, 0.04)$. | Server | Center |
| 66 | replaceable_samples | bool | () | A boolean indicating if there are any replaceable samples in the center. A sample $i$ of center $k$ is said to be replaceable if there are at least `min_replicates` samples across all centers which share the same design factor levels as $i$. `min_replicates` is a user defined parameter, set to 7 by default. | Each center | Server |
| | local_genes_to_replace | set | | The set of genes $g$ for which the Cook's distance is above the cutoff value for any sample in the center. For a given gene $g$ and given sample $i$, the Cook's distance is computed as $\frac{h_{ig}^{(k)}}{p(1-h_{ig}^{(k)})^2}(R^2)_{ig}^{(k)}$ where $(R^2)_{ig}^{(k)}$ is the squared Pearson residual of the negative binomial GLM with log fold changes $\beta_g$ and dispersions $\alpha_g^{\texttt{cooks}}$ computed as $(Y_{ig}^{(k)} - \mu_{ig}^{(k)})^2/(V^{\texttt{NB}})_{ig}^{(k)}$, and $h_{ig}^{(k)}$ is the $i$-th diagonal element of $X^{(k)}H_g^{-1}(X^{(k)})^\top$, where $H_g^{-1}$ is the inverse of the global hat matrix. The cutoff value is set to the 0.99-th quantile of the F-distribution with $p$ and $n-p$ degrees of freedom. Here $\mu_{ig}^{(k)} = \gamma_i^{(k)} \exp(X_i^{(k)} \cdot \beta_g)$ and $(V^{\texttt{NB}})_{ig}^{(k)} = \mu_{ig}^{(k)}(1 + \mu_{ig}^{(k)}\alpha_g^{\texttt{cooks}})$. | Each center | Server |
| 67 | genes_to_replace | set | | The set of genes $g$ for which the Cook's distance is above the cutoff value for any sample in any center (the union of the local genes to replace across all centers). | Server | Center |
| 69 | use_lvl | bool | | Whether or not to use the levels of the design matrix to compute the trimmed mean. Set to `False` in this case, so that the trimmed mean is computed across all samples. | Each center | Server |
| | max_values | nparray | $(G_{\texttt{r}}, )$ | Contains the maximum value of the center normed counts for each gene to replace $g$ across all samples. | Each center | Server |
| | min_values | nparray | $(G_{\texttt{r}}, )$ | Contains the minimum value of the center normed counts for each gene to replace $g$ across all samples. | Each center | Server |
| 70 | use_lvl | bool | | Set to `False` in this case, simply passed on from the previous state. | Server | Center |
| | upper_bounds_thresholds | nparray | $(G_{\texttt{r}}, p)$ | Contains an upper bound of the $r_{\texttt{trim}}$-quantile value of the normed counts for each gene to replace $g$ across all samples. for each gene to replace $g$, the first entry is the upper bound of the $r_{\texttt{trim}}$-quantile value, and the second entry is the upper bound of the $1 - r_{\texttt{trim}}$-quantile value where $r_{\texttt{trim}} = 0.2$. | Server | Center |
| | lower_bounds_thresholds | nparray | $(G_{\texttt{r}}, p)$ | Contains a lower bound of the $r_{\texttt{trim}}$-quantile value of the normed counts for each gene to replace $g$ across all samples. for each gene to replace $g$, the first entry is the lower bound of the $r_{\texttt{trim}}$-quantile value, and the second entry is the lower bound of the $1 - r_{\texttt{trim}}$-quantile value where $r_{\texttt{trim}} = 0.2$. | Server | Center |
| 71 | lower_bounds_thresholds | nparray | $(G_{\texttt{r}}, p)$ | Simply passed on from the previous state. | Each center | Server |
| | n_samples | int | | The number of samples in a center $n_k$ for each center $k$. | Each center | Server |
| | trim_ratio | float | | The trim ratio, defining the upper and lower ratios whose quantile we want to compute. Denoted with $r_{\texttt{trim}}$, equal to 0.2. | Each center | Server |

| | | | | | | |
|---|---|---|---|---|---|---|
| | num_strictly_above | nparray | $(G_{\mathtt{r}}, p)$ | Contains the number of samples whose normed counts value for each gene to replace $g$ is strictly above the average of the upper and lower bounds on the $r_{\mathtt{trim}}$-quantile value of gene $g$ across all samples. for each gene to replace $g$, the first entry is the number of samples whose normed counts value is strictly above the average of the upper and lower bounds on the $r_{\mathtt{trim}}$-quantile value, and the second entry is the number of samples whose normed counts value is strictly above the average of the upper and lower bounds on the $1 - r_{\mathtt{trim}}$-quantile value, where $r_{\mathtt{trim}} = 0.2$. | Each center | Server |
| | use_lvl | bool | | Set to `False` in this case, simply passed on from the previous state. | Each center | Server |
| | upper_bounds_thresholds | nparray | $(G_{\mathtt{r}}, p)$ | Simply passed on from the previous state. | Each center | Server |
| 72 | use_lvl | bool | | Set to `False` in this case, simply passed on from the previous state. | Server | Center |
| | upper_bounds_thresholds | nparray | $(G_{\mathtt{r}}, p)$ | for each gene to replace $g$ and for each ratio $r \in \{r_{\mathtt{trim}}, 1 - r_{\mathtt{trim}}\}$, contains an updated upper bound on the $r$-quantile value of the normed counts of gene $g$ across all samples. For a given gene $g$ and $r$-quantile, it is updated as either its previous value or the average of the previous upper and lower bounds across all samples, depending on the number of samples whose normed counts value is strictly above this average. This number of samples can be computed from the previous shared states. | Server | Center |
| | lower_bounds_thresholds | nparray | $(G_{\mathtt{r}}, p)$ | for each gene to replace $g$ and for each ratio $r \in \{r_{\mathtt{trim}}, 1-r_{\mathtt{trim}}\}$, contains an updated lower bound of the $r$-quantile value of the normed counts of gene $g$ across all samples. For a given gene $g$ and $r$-quantile, it is updated as either its previous value or the average of the previous upper and lower bounds across all samples, depending on the number of samples whose normed counts value is strictly above this average. This number of samples can be computed from the previous shared states. | Server | Center |
| 73 | upper_bounds_thresholds | nparray | $(G_{\mathtt{r}}, p)$ | Simply passed on from the previous state. | Each center | Server |
| | trim_ratio | float | | the trim ratio, set to 0.2 | Each center | Server |
| | lower_bounds_thresholds | nparray | $(G_{\mathtt{r}}, p)$ | Simply passed on from the previous state. | Each center | Server |
| | num_strictly_above | nparray | $(G_{\mathtt{r}}, p)$ | Contains the number of samples whose normed counts value for each gene to replace $g$ is strictly above the average of the upper and lower bounds on the $r_{\mathtt{trim}}$-quantile value of gene $g$ across all samples. for each gene to replace $g$, the first entry is the number of samples whose normed counts value is strictly above the average of the upper and lower bounds on the $r_{\mathtt{trim}}$-quantile value, and the second entry is the number of samples whose normed counts value is strictly above the average of the upper and lower bounds on the $1 - r_{\mathtt{trim}}$-quantile value, where $r_{\mathtt{trim}} = 0.2$. | Each center | Server |
| | n_samples | int | | The number of samples in a center $n_k$ for each center $k$. | Each center | Server |
| | trimmed_local_sum | nparray | $(G_{\mathtt{r}}, )$ | A numpy array of shape $(G, )$ containing the sum of the normed counts across all samples, whose value is strictly above the approximation of the $r_{\mathtt{trim}}$-quantile and less or equal to the approximation of the $1 - r_{\mathtt{trim}}$ quantile. For each gene to replace $g$, the entry is the trimmed local sum. | Each center | Server |
| | use_lvl | bool | | Set to `False` in this case, simply passed on from the previous state. | Each center | Server |
| 74 | trimmed_mean_normed_counts | nparray | $(G_{\mathtt{r}}, )$ | For each gene to replace $g$, the trimmed mean of the normed counts across all samples, denoted with $\overline{Z}_g^{\mathtt{trim}}$ with trim ratio set to 0.2. | Server | Center |
| 75 | loc_new_all_zeroes | nparray | $(G_{\mathtt{r}}, )$ | A boolean array which for each gene to replace $g$ indicates if the new count matrix of the center is all zeroes (across samples) for this gene (the new count matrix is computed by imputing the Cook's outliers with the trimmed mean of the normed counts times the size factor). | Each center | Server |
| 76 | new_all_zeroes | nparray | $(G_{\mathtt{r}}, )$ | A boolean array which for each gene to replace $g$ indicates if all counts across all centers are zero for this gene. | Server | Center |
| 77 | local_features | nparray | $(p, G_{\mathtt{act}})$ | $\Phi^{(k)} := (X^{(k)})^\top Z^{(k)}$, where $Z^{(k)}$ is the normalized counts in the center $k$ on the set of genes to replace, where the value of Cook's outliers have been replaced using a trimmed mean. | Each center | Server |
| 78 | global_feature_vector | nparray | $(p, G_{\mathtt{r}})$ | The sum of the local features across all centers $\Phi = \sum_k \Phi^{(k)}$. | Server | Center |
| 79 | local_rough_dispersions | nparray | $(G_{\mathtt{r}}, )$ | A rough estimate of the dispersions of the genes on the local data. It is computed as a method of moments estimator of the dispersion of the normalized counts $(\alpha_g^{\mathtt{MoM},2})^{(k)} = \sum_{i=1}^{n_k}((Z_{ig}^{(k)})^2 - (\widehat{Z}_{ig}^{(k)})^2)/(\widehat{Z}_{ig}^{(k)})^2$ where $\widehat{Z}_g^{(k)} = X^{(k)} G^{-1} \Phi_g$ corresponds to a linear estimate of the normed counts. | Each center | Server |
| | local_n_obs | int | | The number of observations in a center $n_k$. | Each center | Server |
| | local_n_params | int | | The number of parameters in a center $p$ (i.e., the number of columns of the design matrix). This number is the same across all centers, but the server does not directly have access to it since it is stateless. | Each center | Server |
| 80 | rough_dispersions | nparray | $(G_{\mathtt{r}}, )$ | A rough estimate of the dispersions obtained by essentially averaging local rough dispersions $(\alpha_g^{\mathtt{MoM},2} = \max(\frac{1}{n-p}\sum_k (\alpha_g^{\mathtt{MoM},2})^{(k)}, 0))$ | Server | Center |
| 81 | local_inverse_size_mean | float | $()$ | The average of the inverse of the size factors in a center $\overline{1/\gamma}^{(k)} = \frac{1}{n_k}\sum_{i=1}^{n_k} 1/\gamma_i^{(k)}$. | Each center | Server |
| | rough_dispersions | nparray | $(G_{\mathtt{r}}, )$ | The rough estimates computed at the previous step and passed on. | Each center | Server |
| | local_n_obs | int | | The number of observations in a center $n_k$. | Each center | Server |
| | local_squared_squared_mean | nparray | $(G_{\mathtt{r}}, )$ | The average of the square normed counts in a center, per gene $\overline{Y^2} := \frac{1}{n_k}\sum_{i=1}^{n_k}(Y_{ig}^{(k)})^2$. | Each center | Server |
| | local_counts_mean | nparray | $(G_{\mathtt{r}}, )$ | The averag e of the counts in a center, per gene $\overline{Y}_g^{(k)} = \frac{1}{n_k}\sum_{i=1}^{n_k} Y_{ig}^{(k)}$. | Each center | Server |
| 82 | tot_num_samples | int | | The total number of samples across all centers $n$. | Server | Center |
| | non_zero | nparray | $(G_{\mathtt{r}}, )$ | A boolean array indicating which genes have non-zero counts in at least one center. | Server | Center |

| | | | | | | |
|---|---|---|---|---|---|---|
| | MoM_dispersions | nparray | $(G_{\mathtt{r}},)$ | The dispersions obtained by the method of moments. First, the aggregated mean counts, squared mean counts and inverse size factors are computed ($\overline{1/\gamma} = \sum_k \frac{n_k}{n} \overline{1/\gamma}^{(k)}$), $\overline{Y}_g = \sum_k \frac{n_k}{n} \overline{Y}_g^{(k)}$, $\overline{Y^2} = \sum_k \frac{n_k}{n} \overline{Y^2}_g^{(k)}$)). The variance is then estimated with $\widehat{\sigma}_g^2 = \frac{n}{n-1}(\overline{Y^2}_g - \overline{Y}_g^2)$ and a method of moments estimate is the constructed as $\alpha_g^{\mathtt{MoM},1} = \frac{\widehat{\sigma}_g^2 - \overline{1/\gamma}_g \times \overline{Y}_g}{\overline{Y}_g^2}$. The final MoM estimate $\alpha_g^{\mathtt{MoM}}$ for gene $g$ is the clipped value of $\min(\alpha_g^{\mathtt{MoM},1}, \alpha_g^{\mathtt{MoM},2})$ between `min_disp` and max(`max_disp`, $n$) | Server | Center |
| | tot_counts_mean | nparray | $(G_{\mathtt{r}},)$ | The average of the counts across all centers, per gene obtained by summing the local counts means. | Server | Center |
| 84 | local_log_features | nparray | $(G_{\mathtt{r}}, p)$ | $\Phi_k = (X^{(k)})^\top \log(Z^{(k)} + 0.1)$. | Each center | Server |
| | gram_full_rank | bool | () | A boolean indicating whether the global Gramm matrix $G$ is full rank. | Each center | Server |
| | n_non_zero_genes | int | | The number of genes with non-zero counts in at least one center. | Each center | Server |
| | global_gram_matrix | nparray | $(p, p)$ | The sum of the local Gramm matrix across all centers $G = \sum_k G^{(k)}$. | Each center | Server |
| 85 | round_number_irls | int | | The current round number of the IRLS algorithm. Initialized at 0. | Server | Center |
| | global_nll | nparray | $(G_{\mathtt{r}},)$ | For each gene $g$, the negative log likelihood of the negative binomial GLM, initialized at 1000 for all genes. | Server | Center |
| | irls_mask | nparray | $(G_{\mathtt{r}},)$ | A boolean array indicating which non zero genes are currently being optimized by the IRLS algorithm. Initialized to `True`. | Server | Center |
| | irls_diverged_mask | nparray | $(G_{\mathtt{r}},)$ | A boolean array indicating which non zero genes have caused the IRLS algorithm to diverge. Initialized to `False` | Server | Center |
| | beta | nparray | $(G_{\mathtt{r}}, p)$ | The initial value of the $\beta$ parameter. If the Gramm matrix is full rank, is set to $\beta_g = G^{-1}(\sum_k \Phi_g^{(k)})$ for all genes $g$. Otherwise, it is set as a weighed average of the normed log means, i.e., $\beta_g = \sum_k \frac{n_k}{n} \overline{\log(Z)}_g^{(k)}$, where $\overline{\log(Z)}_g^{(k)} = \frac{1}{n_k} \sum_{i=1}^{n_k} \log(Z_{ig}^{(k)})$ is computed locally. | Server | Center |
| 86 | irls_gene_names | Index | $(G_{\mathtt{act}},)$ | The gene names of the active genes. | Each center | Server |
| | local_features | nparray | $(G_{\mathtt{act}}, p)$ | NaN | Each center | Server |
| | local_hat_matrix | nparray | $(G_{\mathtt{act}}, p, p)$ | For each gene $g$, the hat matrix $H_g^{(k)} = (X^{(k)})^\top W_g^{(k)} X^{(k)}$ where $W_g^{(k)} \in \mathbb{R}^{n_k \times n_k}$ is the diagonal matrix with diagonal entries $\frac{\mu_{ig}^{(k)}}{1+\mu_{ig}^{(k)}\alpha_g}$ for $1 \leq i \leq n_k$. $\alpha_g$ is the dispersion estimate of the gene and $\mu_{ig}^{(k)}$ is the expected value of the gene for sample $i$ for parameter $\beta$, that is $\gamma_g^{(k)} \exp(X_i^{(k)} \cdot \beta_g)$. | Each center | Server |
| | beta | nparray | $(G_{\mathtt{r}}, p)$ | The log fold change $\beta$ parameter. Not modified. | Each center | Server |
| | local_nll | nparray | $(G_{\mathtt{act}},)$ | For each gene $g$, the negative log likelihood of the negative binomial GLM with fixed dispersion at the $\beta_g$ parameter, computed on the local data. | Each center | Server |
| | round_number_irls | int | | The current round number of the IRLS algorithm. Simply passed on. | Each center | Server |
| | global_nll | nparray | $(G_{\mathtt{r}},)$ | For each gene $g$, the negative log likelihood at the $\beta_g$ parameter. Simply passed on. | Each center | Server |
| | irls_mask | nparray | $(G_{\mathtt{r}},)$ | A boolean array indicating which non zero genes are currently being optimized by the IRLS algorithm. Simply passed on. | Each center | Server |
| | irls_diverged_mask | nparray | $(G_{\mathtt{r}},)$ | A boolean array indicating which non zero genes have caused the IRLS algorithm to diverge. Simply passed on | Each center | Server |
| 87 | round_number_irls | int | | The current round number of the IRLS algorithm, incremented by 1 during the local step. | Server | Center |
| | irls_mask | nparray | $(G_{\mathtt{r}},)$ | A boolean array indicating which non zero genes are currently being optimized by the IRLS algorithm. Genes whose $\beta$ parameter has diverged are removed from the optimization. Moreover, the deviance ratio is computed for each active gene $g$, as $\frac{\lvert 2\,\mathcal{L}(\beta_g) - 2 \cdot \mathcal{L}(\beta_{g,\mathrm{prev}})\rvert}{\lvert 2\,\mathcal{L}(\beta)\rvert + 0.1}$ where $\beta_{g,\mathrm{prev}}$ is the value of $\beta_g$ at the previous iteration. If this deviance is smaller than the user inputed `beta_tol`, the gene is considered to have converged and is set to `False` in the `irls_mask`, meaning it will not be optimized further. | Server | Center |
| | irls_diverged_mask | nparray | $(G_{\mathtt{r}},)$ | A boolean array indicating which non zero genes have caused the IRLS algorithm to diverge. A gene $g$ is considered to have diverged if any component of $\beta_g \in \mathbb{R}^p$ has absolute value above the threshold `max_beta`. | Server | Center |
| | beta | nparray | $(G_{\mathtt{r}}, p)$ | The updated value of the $\beta$ parameter after applying the IRLS update for the active genes. At the last step, is not updated. | Server | Center |
| | global_nll | nparray | $(G_{\mathtt{r}},)$ | For each gene $g$, the negative log-likelihood at the $\beta_g$ parameter, built by summing the local nlls from the centers | Server | Center |
| 88 | local_nll | nparray | $(1, G_{\mathtt{act}})$ | The nll of the negative binomial GLM with fixed dispersion at the $\beta_g$ parameter, computed on the local data, for each gene $g$. | Each center | Server |
| | ascent_direction_on_mask | NoneType | | The ascent direction on the active genes, which is set to `None` at the beginning. | Each center | Server |
| | round_number_PQN | int | | The current round number of the Proximal Quasi-Newton algorithm, initialized at 0. | Each center | Server |
| | newton_decrement_on_mask | NoneType | | The newton decrement on the active genes, which is set to `None` at the beginning. | Each center | Server |
| | global_reg_nll | nparray | $(G_{\mathtt{r}},)$ | The regularized negative log likelihood of the negative binomial GLM with fixed dispersion at the $\beta_g$ parameter intialized at `np.nan` since no nll has been computed at this stage. | Each center | Server |
| | PQN_mask | nparray | $(G_{\mathtt{r}},)$ | A boolean array indicating which non zero genes are currently being optimized by the Proximal Quasi-Newton algorithm (active genes). The genes which have diverged during the IRLS algorithm and those which have not converged at the end of the prescribed iterations are set to `True`. | Each center | Server |
| | PQN_diverged_mask | nparray | $(G_{\mathtt{r}},)$ | A boolean array indicating which non zero genes have caused the Proximal Quasi-Newton algorithm to diverge. Initialized to `False`. | Each center | Server |
| | local_gradient | nparray | $(1, G_{\mathtt{act}}, p)$ | For each gene $g$, the gradient of the negative log likelihood of the negative binomial GLM with fixed dispersion at the $\beta_g$ parameter, computed on the local data, i.e., $\nabla_\beta \mathcal{L}^{(k)}(\beta_g) = -(X^{(k)})^\top Y_g^{(k)} + (X^{(k)})^\top \left(\frac{1}{\alpha_g} + Y_g^{(k)}\right) \frac{\mu_g^{(k)}}{\frac{1}{\alpha_g} + \mu_g^{(k)}}$, where $\mu_g^{(k)}$ and $Y_g^{(k)}$ are both vectors in $\mathbb{R}^{n_k}$ and $\alpha_g$ is a scalar. TODO ref equation. | Each center | Server |

| | | | | | | |
|---|---|---|---|---|---|---|
| | irls_diverged_mask | nparray | $(G_{\mathtt{r}},)$ | A boolean array indicating which non zero genes have caused the IRLS algorithm to diverge. Passed on without modification. | Server | Center |
| | beta | nparray | $(G_{\mathtt{r}}, p)$ | For each gene $g$, the log fold change $\beta_g$. This value has been updated by the Proximal Quasi-Newton algorithm for all active genes where an admissible step size was found using the Armijo condition (see equation 2.5 of [1]) | Server | Center |
| | PQN_diverged_mask | nparray | $(G_{\mathtt{r}},)$ | A boolean array indicating which non zero genes have caused the Proximal Quasi-Newton algorithm to diverge. A gene $g$ is considered to have diverged if for all step sizes $\delta \in \{1, 1/2, 1/4, ..., 1/2^{N_{\mathtt{ls}}-1}\}$, and for the computed ascent direction $\Delta_g$, the Armijo condition is not satisfied, i.e., if $\mathcal{L}^{\lambda}(\beta_g) - \mathcal{L}^{\lambda}(\beta_g - \delta\Delta_g) \leq \delta\, \mathtt{PQN\_c1}\, \nabla_\beta\, \mathcal{L}^{\lambda}(\beta_g) \cdot \Delta_g$ where $\mathcal{L}^{\lambda}$ is the regularized negative log likelihood, PQN_c1 is the Armijo parameter which can be set by the user (default is $10^{-4}$), and $\nabla_\beta\, \mathcal{L}^{\lambda}(\beta_g)$ is the gradient of the regularized negative log likelihood. | Server | Center |
| | newton_decrement_on_mask | nparray | $(G_{\mathtt{act}},)$ | The newton decrement on the new active genes $g$, computed at the new $\beta_g$ iterate. For more details, see description of 19. | Server | Center |
| 92 | PQN_mask | nparray | $(G_{\mathtt{r}},)$ | A boolean array indicating which non zero genes are currently being optimized by the Proximal Quasi-Newton algorithm (active genes), passed on without modification. | Each center | Server |
| | beta | nparray | $(G_{\mathtt{r}}, p)$ | For each gene $g$, the log fold change $\beta_g$, which is simply passed on. | Each center | Server |
| | local_nll | nparray | $(N_{\mathtt{ls}}, G_{\mathtt{act}})$ | The nll of the negative binomial GLM with fixed dispersion at the $\beta_g$ parameter, computed on the local data, for each gene $g$ and for each potential next step $\beta_g - \delta\Delta_g$ for $\delta \in \{1, 1/2, 1/4, ..., 1/2^{N_{\mathtt{ls}}-1}\}$. | Each center | Server |
| | local_fisher | nparray | $(N_{\mathtt{ls}}, G_{\mathtt{act}}, p, p)$ | For each potential step size $\delta \in \{1, 1/2, 1/4, ..., 1/2^{N_{\mathtt{ls}}-1}\}$, and each gene $g$, the Fisher information matrix of the negative binomial GLM with fixed dispersion computed at $\beta_g - \delta\Delta_g$. Formally, $n_k \mathcal{I}^{(k)}_{\delta,g} = (X^{(k)})^{\top} W^{(k)}_{\delta,g} X^{(k)}$ where $W^{(k)}_{\delta,g} \in \mathbb{R}^{n_k \times n_k}$ is the diagonal matrix with diagonal entries $W^{(k)}_{\delta,gii} = \frac{\mu^{(k)}_{\delta,ig}}{1+\mu^{(k)}_{\delta,ig}\alpha_g}$ for $1 \leq i \leq n_k$. $\alpha_g$ is the dispersion estimate of the gene and $\mu^{(k)}_{\delta,ig}$ is the expected value of the gene $g$ for sample $i$ for parameter $\beta_g - \delta\Delta_g$, that is $\gamma^{(k)}_g \exp(X^{(k)}_i \cdot (\beta_g - \delta\Delta_g))$. | Each center | Server |
| | local_gradient | nparray | $(N_{\mathtt{ls}}, G_{\mathtt{act}}, p)$ | For each potential step size $\delta$ and each gene $g$, the gradient of the negative log likelihood of the negative binomial GLM with fixed dispersion at the $\beta_g - \delta\Delta_g$ parameter computed on the local data. | Each center | Server |
| | PQN_diverged_mask | nparray | $(G_{\mathtt{r}},)$ | A boolean array indicating which non zero genes have caused the Proximal Quasi-Newton algorithm to diverge. Passed on without modification. | Each center | Server |
| | global_reg_nll | nparray | $(G_{\mathtt{r}},)$ | The regularized negative log likelihood of the negative binomial GLM with fixed dispersion at the $\beta_g$ parameter, simply passed on. | Each center | Server |
| | newton_decrement_on_mask | nparray | $(G_{\mathtt{act}},)$ | The newton decrement on the active genes, passed on without modification. | Each center | Server |
| | round_number_PQN | int | | The current round number of the Proximal Quasi-Newton algorithm, passed on without modification. | Each center | Server |
| | irls_diverged_mask | nparray | $(G_{\mathtt{r}},)$ | A boolean array indicating which non zero genes have caused the IRLS algorithm to diverge. Passed on without modification. | Each center | Server |
| | ascent_direction_on_mask | nparray | $(G_{\mathtt{act}}, p)$ | The ascent direction on the active genes, passed on without modification. | Each center | Server |
| 93 | beta | nparray | $(G_{\mathtt{r}}, p)$ | For each gene $g$, the log fold change $\beta_g$. This value has been updated by the Proximal Quasi-Newton algorithm for all active genes where an admissible step size was found using the Armijo condition (see equation 2.5 of [1]) | Server | Center |
| | irls_diverged_mask | nparray | $(G_{\mathtt{r}},)$ | A boolean array indicating which non zero genes have caused the IRLS algorithm to diverge. Passed on without modification. | Server | Center |
| | PQN_diverged_mask | nparray | $(G_{\mathtt{r}},)$ | A boolean array indicating which non zero genes have caused the Proximal Quasi-Newton algorithm to diverge. As this is the last step of the optimization, a gene $g$ is considered to have diverged not only if the Armijo condition is never satisfied, but also if the convergence criterion is not met. A gene is said to converge if the relative difference between two successive iterates is smaller than a threshold value PQN_ftol, i.e., $\lvert \mathcal{L}^{\lambda}(\beta_g) - \mathcal{L}^{\lambda}(\beta_{g,\text{prev}}) \rvert \leq \mathtt{PQN\_ftol}\, \max(\mathcal{L}^{\lambda}(\beta_g), \mathcal{L}^{\lambda}(\beta_{g,\text{prev}}), 1)$. | Server | Center |
| 95 | unique_counts | Series | $(p,)$ | A pandas series indexed by unique level combinations (all design factor levels that coexist) and containing the number of samples in each level combination. In the single factor case the index will simply be the unique levels of the factor. | Each center | Server |
| 96 | counts_by_lvl | Series | $(L,)$ | A pandas series indexed by the unique levels of the design factors across all centers, and whose values are the number of samples in each level combination. | Server | Center |
| 99 | max_disp | int | | The effective maximum value of the dispersion parameter $\max(\mathtt{max\_disp}, n)$. | Each center | Server |
| | grid | nparray | $(G_{\mathtt{r}}, N_{\mathtt{gs}})$ | The logarithmic grid between min_disp and $\max(\mathtt{max\_disp}, n)$ of size $N_{\mathtt{gs}}$. | Each center | Server |
| | CR_summand | nparray | $(G_{\mathtt{r}}, N_{\mathtt{gs}}, p, p)$ | For all genes $g$ and all dispersion values $\alpha_g$ in the logarithmic grid between min_disp $\max(\mathtt{max\_disp}, n)$ of size $N_{\mathtt{gs}}$, the matrix of the Cox-Reid regularization term pertaining to the samples in the center $k$. This matrix is computed as $\mathrm{CR}^{(k)}_g = (X^{(k)})^{\top} W^{(k)}_g X^{(k)}$ where $W^{(k)}_g \in \mathbb{R}^{n_k \times n_k}$ is the diagonal matrix with diagonal entries $\frac{\mu^{(k)}_{ig}}{1+\mu^{(k)}_{ig}\alpha_g}$ for $1 \leq i \leq n_k$. $\mu^{(k)}_{ig}$ is the expected value of the gene for sample $i$, whose expression can be found in the description of the nll variable. | Each center | Server |

| | | | | | | |
|---|---|---|---|---|---|---|
| | nll | nparray | $(G_{\mathtt{r}}, N_{\mathtt{gs}})$ | For all genes $g$ and all dispersion values $\alpha_g$ in the logarithmic grid between `min_disp` max(`max_disp`, $n$) of size $N_{\mathtt{gs}}$ the negative log likelihood of the negative binomial GLM with fixed mean parameters $\mu_{ig}^{(k)}$ over the set of loca samples and dispersion parameter $\alpha_g$. If the numbe of unique levels of all factors is equal to the numbe of parameters $p$, then $\mu_{ig}^{(k)}$ is defined as `nan` for zerc genes, and as the minimum of $\gamma_i^{(k)}$ $(X^{(k)})^\top G^{-1}\Phi_g$ anc `min_mu` := 0.5 (for the definition of $G$ and $\Phi_g$, see step 8). Otherwise, $\mu_{ig}^{(k)}$ is defined as $\gamma_i^{(k)}$ $\exp(X_i^{(k)} \cdot \beta_g)$ or non-zero genes, where $\beta_g$ is the log fold change result o the IRLS algorithm, and as `nan` on zero genes. | Each center | Server |
| | non_zero | nparray | $(G_{\mathtt{r}},)$ | A boolean array indicating which genes have non-zerc counts in at least one center. | Each center | Server |
| 100 | lower_log_bounds | nparray | $(G_{\mathtt{r}},)$ | For each gene $g$, the maximum of the log of the min dispersion and $\alpha_g - \delta$ where $\delta$ is the current mesh size of the grid and $\alpha_g$ is the current dispersion estimate. This value will be used as a lower bound for the next grid search fo the dispersion parameter. | Server | Center |
| | upper_log_bounds | nparray | $(G_{\mathtt{r}},)$ | For each gene $g$, the minimum of the log of the max dispersion and $\alpha_g + \delta$ where $\delta$ is the current mesh size o the grid and $\alpha_g$ is the current dispersion estimate. This value will be used as an upper bound for the next gric search for the dispersion parameter. | Server | Center |
| | genewise_dispersions | nparray | $(G_{\mathtt{r}},)$ | For each gene $g$, the current estimate of the dispersior parameter $\alpha_g$. This estimate is computed by first computing the global nll (summing all local nlls) as well as the global Cox-Reid regularization term, which is half the log determinant of the sum of the local Cox-Reid matrices. The Cox-Reid regularized nll per gene and pe dispersion in the grid is obtained by summing the regularization term and the nll. Finally, for every gene, the dispersion parameter is estimated by taking the minizer of this regularized nll on the grid of size $N_{\mathtt{gs}}$. | Server | Center |
| 101 | non_zero | nparray | $(G_{\mathtt{r}},)$ | A boolean array indicating which genes have non-zerc counts in at least one center. | Each center | Server |
| | max_disp | int | | The effective maximum value of the dispersion paramete max(`max_disp`, $n$). | Each center | Server |
| | grid | nparray | $(G_{\mathtt{r}}, N_{\mathtt{gs}})$ | The logarithmic grid between the exponential of the lowe log bound and of the upper log bound of size $N_{\mathtt{gs}}$. | Each center | Server |
| | CR_summand | nparray | $(G_{\mathtt{r}}, N_{\mathtt{gs}}, p, p)$ | For all genes $g$ and all dispersion values $\alpha_g$ in the logarithmic grid between the exponential of the lower log bound and of the upper log bound of size $N_{\mathtt{gs}}$, the matrix of the Cox-Reid regularization term pertaining to the samples in the center $k$. This matrix is computed as $\mathrm{CR}_g^{(k)} = (X^{(k)})^\top W_g^{(k)} X^{(k)}$ where $W_g^{(k)} \in \mathbb{R}^{n_k \times n_k}$ is the diagonal matrix with diagonal entries $\frac{\mu_{ig}^{(k)}}{1+\mu_{ig}^{(k)}\alpha_g}$ for $1 \leq i \leq n_k$. $\mu_{ig}^{(k)}$ is the expected value of the gene for sample $i$, whose expression can be found in the description of the nll variable. | Each center | Server |
| | nll | nparray | $(G_{\mathtt{r}}, N_{\mathtt{gs}})$ | For all genes $g$ and all dispersion values $\alpha_g$ in the logarithmic grid between the exponential of the lower log bound and of the upper log bound of size $N_{\mathtt{gs}}$, the negative log likelihood of the negative binomial GLM with fixed mean parameters $\mu_{ig}^{(k)}$ over the set of local samples and dispersion parameter $\alpha_g$. If the number of unique levels of all factors is equal to the number of parameters $p$, then $\mu_{ig}^{(k)}$ is defined as `nan` for zero genes, and as the minimum of $\gamma_i^{(k)}$ $(X^{(k)})^\top G^{-1}\Phi_g$ and `min_mu` := 0.5 (for the definition of $G$ and $\Phi_g$, see step 8). Otherwise $\mu_{ig}^{(k)}$ is defined as $\gamma_i^{(k)}$ $\exp(X_i^{(k)} \cdot \beta_g)$ on non-zero genes where $\beta_g$ is the log fold change result of the IRLS algorithm, and as `nan` on zero genes. | Each center | Server |
| 102 | lower_log_bounds | nparray | $(G_{\mathtt{r}},)$ | For each gene $g$, the maximum of the log of the min dispersion and $\alpha_g - \delta$ where $\delta$ is the current mesh size of the grid and $\alpha_g$ is the current dispersion estimate. This value will be used as a lower bound for the next grid search fo the dispersion parameter. | Server | Center |
| | genewise_dispersions | nparray | $(G_{\mathtt{r}},)$ | For each gene $g$, the current estimate of the dispersior parameter $\alpha_g$. This estimate is computed by first computing the global nll (summing all local nlls) as well as the global Cox-Reid regularization term, which is half the log determinant of the sum of the local Cox-Reid matrices. The Cox-Reid regularized nll per gene and pe dispersion in the grid is obtained by summing the regularization term and the nll. Finally, for every gene, the dispersion parameter is estimated by taking the minizer of this regularized nll on the grid of size $N_{\mathtt{gs}}$. | Server | Center |
| | upper_log_bounds | nparray | $(G_{\mathtt{r}},)$ | For each gene $g$, the minimum of the log of the max dispersion and $\alpha_g + \delta$ where $\delta$ is the current mesh size o the grid and $\alpha_g$ is the current dispersion estimate. This value will be used as an upper bound for the next gric search for the dispersion parameter. | Server | Center |
| 104 | max_disp | int | | The effective maximum value of the dispersion paramete max(`max_disp`, $n$). | Each center | Server |
| | nll | nparray | $(G_{\mathtt{r}}, N_{\mathtt{gs}})$ | For all genes $g$ and all dispersion values $\alpha_g$ in the logarithmic grid between `min_disp` max(`max_disp`, $n$) of size $N_{\mathtt{gs}}$ the negative log likelihood of the negative binomial GLM with fixed mean parameters $\mu_{ig}^{(k)}$ over the set of loca samples and dispersion parameter $\alpha_g$. If the numbe of unique levels of all factors is equal to the numbe of parameters $p$, then $\mu_{ig}^{(k)}$ is defined as `nan` for zerc genes, and as the minimum of $\gamma_i^{(k)}$ $(X^{(k)})^\top G^{-1}\Phi_g$ anc `min_mu` := 0.5 (for the definition of $G$ and $\Phi_g$, see step 8). Otherwise, $\mu_{ig}^{(k)}$ is defined as $\gamma_i^{(k)}$ $\exp(X_i^{(k)} \cdot \beta_g)$ or non-zero genes, where $\beta_g$ is the log fold change result o the IRLS algorithm, and as `nan` on zero genes. | Each center | Server |

| | | | | | | |
|---|---|---|---|---|---|---|
| | CR_summand | nparray | $(G_r, N_{gs}, p, p)$ | For all genes $g$ and all dispersion values $\alpha_g$ in the logarithmic grid between min_disp $\max(\texttt{max\_disp}, n)$ of size $N_{gs}$ the matrix of the Cox-Reid regularization term pertaining to the samples in the center $k$. This matrix is computed as $\mathrm{CR}_g^{(k)} = (X^{(k)})^\top W_g^{(k)} X^{(k)}$ where $W_g^{(k)} \in \mathbb{R}^{n_k \times n_k}$ is the diagonal matrix with diagonal entries $\frac{\mu_{ig}^{(k)}}{1+\mu_{ig}^{(k)}\alpha_g}$ for $1 \leq i \leq n_k$. $\mu_{ig}^{(k)}$ is the expected value of the gene for sample $i$, whose expression can be found in the description of the nll variable. | Each center | Server |
| | grid | nparray | $(G_r, N_{gs})$ | The logarithmic grid between min_disp and $\max(\texttt{max\_disp}, n)$ of size $N_{gs}$. | Each center | Server |
| | non_zero | nparray | $(G_r, )$ | A boolean array indicating which genes have non-zero counts in at least one center. | Each center | Server |
| | reg | nparray | $(G_r, N_{gs})$ | For all genes $g$ and all dispersion values $\alpha_g$ in the logarithmic grid between min_disp and $\max(\texttt{max\_disp}, n)$ of size $N_{gs}$, a regularization term that is equal to $0.5 \times (\log(\alpha_g) - \log(\alpha_g^{\texttt{trend}}))^2/\sigma^2_{\texttt{trend}}$, and which comes from the prior on the distribution of the log dispersions around the trend curve. | Each center | Server |
| 105 | lower_log_bounds | nparray | $(G_r, )$ | For each gene $g$, the maximum of the log of the min dispersion and $\alpha_g - \delta$ where $\delta$ is the current mesh size of the grid and $\alpha_g$ is the current dispersion estimate. This value will be used as a lower bound for the next grid search for the dispersion parameter. | Server | Center |
| | MAP_dispersions | nparray | $(G_r, )$ | For each gene $g$, the current estimate of the MAP dispersion parameter $\alpha_g$. This estimate is computed by first computing the global nll (summing all local nlls) as well as the global Cox-Reid regularization term, which is half the log determinant of the sum of the local Cox-Reid matrices, and the regularization coming from the prior on the log dispersions around the trend curve. The Cox-Reid, prior regularized nll per gene and per dispersion in the grid is obtained by summing the regularization terms and the nll. Finally, for every gene, the MAP dispersion parameter is estimated by taking the minizer of this regularized nll on the grid of size $N_{gs}$. | Server | Center |
| | upper_log_bounds | nparray | $(G_r, )$ | For each gene $g$, the minimum of the log of the max dispersion and $\alpha_g + \delta$ where $\delta$ is the current mesh size of the grid and $\alpha_g$ is the current dispersion estimate. This value will be used as an upper bound for the next grid search for the dispersion parameter. | Server | Center |
| 106 | CR_summand | nparray | $(G_r, N_{gs}, p, p)$ | For all genes $g$ and all dispersion values $\alpha_g$ in the logarithmic grid between the exponential of the lower log bound and of the upper log bound of size $N_{gs}$, the matrix of the Cox-Reid regularization term pertaining to the samples in the center $k$. This matrix is computed as $\mathrm{CR}_g^{(k)} = (X^{(k)})^\top W_g^{(k)} X^{(k)}$ where $W_g^{(k)} \in \mathbb{R}^{n_k \times n_k}$ is the diagonal matrix with diagonal entries $\frac{\mu_{ig}^{(k)}}{1+\mu_{ig}^{(k)}\alpha_g}$ for $1 \leq i \leq n_k$. $\mu_{ig}^{(k)}$ is the expected value of the gene for sample $i$, whose expression can be found in the description of the nll variable. | Each center | Server |
| | reg | nparray | $(G_r, N_{gs})$ | For all genes $g$ and all dispersion values $\alpha_g$ in the logarithmic grid between the exponential of the lower log bound and of the upper log bound of size $N_{gs}$, a regularization term that is equal to $0.5 \times (\log(\alpha_g) - \log(\alpha_g^{\texttt{trend}}))^2/\sigma^2_{\texttt{trend}}$, and which comes from the prior on the distribution of the log dispersions around the trend curve. | Each center | Server |
| | non_zero | nparray | $(G_r, )$ | A boolean array indicating which genes have non-zero counts in at least one center. | Each center | Server |
| | max_disp | int | | The effective maximum value of the dispersion parameter $\max(\texttt{max\_disp}, n)$. | Each center | Server |
| | nll | nparray | $(G_r, N_{gs})$ | For all genes $g$ and all dispersion values $\alpha_g$ in the logarithmic grid between the exponential of the lower log bound and of the upper log bound of size $N_{gs}$, the negative log likelihood of the negative binomial GLM with fixed mean parameters $\mu_{ig}^{(k)}$ over the set of local samples and dispersion parameter $\alpha_g$. If the number of unique levels of all factors is equal to the number of parameters $p$, then $\mu_{ig}^{(k)}$ is defined as nan for zero genes, and as the minimum of $\gamma_i^{(k)}$ $(X^{(k)})^\top G^{-1}\Phi_g$ and $\texttt{min\_mu} := 0.5$ (for the definition of $G$ and $\Phi_g$, see step 8). Otherwise $\mu_{ig}^{(k)}$ is defined as $\gamma_i^{(k)}$ $\exp(X_i^{(k)} \cdot \beta_g)$ on non-zero genes where $\beta_g$ is the log fold change result of the IRLS algorithm, and as nan on zero genes. | Each center | Server |
| | grid | nparray | $(G_r, N_{gs})$ | The logarithmic grid between the exponential of the lower log bound and of the upper log bound of size $N_{gs}$. | Each center | Server |
| 107 | upper_log_bounds | nparray | $(G_r, )$ | For each gene $g$, the minimum of the log of the max dispersion and $\alpha_g + \delta$ where $\delta$ is the current mesh size of the grid and $\alpha_g$ is the current dispersion estimate. This value will be used as an upper bound for the next grid search for the dispersion parameter. | Server | Center |
| | lower_log_bounds | nparray | $(G_r, )$ | For each gene $g$, the maximum of the log of the min dispersion and $\alpha_g - \delta$ where $\delta$ is the current mesh size of the grid and $\alpha_g$ is the current dispersion estimate. This value will be used as a lower bound for the next grid search for the dispersion parameter. | Server | Center |
| | MAP_dispersions | nparray | $(G_r, )$ | For each gene $g$, the current estimate of the MAP dispersion parameter $\alpha_g$. This estimate is computed by first computing the global nll (summing all local nlls) as well as the global Cox-Reid regularization term, which is half the log determinant of the sum of the local Cox-Reid matrices, and the regularization coming from the prior on the log dispersions around the trend curve. The Cox-Reid, prior regularized nll per gene and per dispersion in the grid is obtained by summing the regularization terms and the nll. Finally, for every gene, the MAP dispersion parameter is estimated by taking the minizer of this regularized nll on the grid of size $N_{gs}$. | Server | Center |
| 109 | n_non_zero_genes | int | | The number of genes with non-zero counts in at least one center. | Each center | Server |
| | global_gram_matrix | nparray | $(p, p)$ | The sum of the local Gramm matrix across all centers $G = \sum_k G^{(k)}$. | Each center | Server |
| | local_log_features | nparray | $(G_r, p)$ | $\Phi_k = (X^{(k)})^\top \log(Z^{(k)} + 0.1)$. | Each center | Server |
| | gram_full_rank | bool | () | A boolean indicating whether the global Gramm matrix $G$ is full rank. | Each center | Server |
| 110 | global_nll | nparray | $(G_r, )$ | For each gene $g$, the negative log likelihood of the negative binomial GLM, initialized at 1000 for all genes. | Server | Center |

| | | | | | | |
|---|---|---|---|---|---|---|
| | irls_mask | nparray | $(G_{\mathtt{r}},)$ | A boolean array indicating which non zero genes are currently being optimized by the IRLS algorithm. Initialized to `True`. | Server | Center |
| | irls_diverged_mask | nparray | $(G_{\mathtt{r}},)$ | A boolean array indicating which non zero genes have caused the IRLS algorithm to diverge. Initialized to `False` | Server | Center |
| | beta | nparray | $(G_{\mathtt{r}}, p)$ | The initial value of the $\beta$ parameter. If the Gramm matrix is full rank, is set to $\beta_g = G^{-1}(\sum_k \Phi_g^{(k)})$ for al genes $g$. Otherwise, it is set as a weighed average of the normed log means, i.e., $\beta_g = \sum_k \frac{n_k}{n} \overline{\log(Z)}_g^{(k)}$, where $\overline{\log(Z)}_g^{(k)} = \frac{1}{n_k} \sum_{i=1}^{n_k} \log(Z_{ig}^{(k)})$ is computed locally. | Server | Center |
| | round_number_irls | int | | The current round number of the IRLS algorithm. Initialized at 0. | Server | Center |
| 111 | local_hat_matrix | nparray | $(G_{\mathtt{act}}, p, p)$ | For each gene $g$, the hat matrix $H_g^{(k)} = (X^{(k)})^\top W_g^{(k)} X^{(k)}$ where $W_g^{(k)} \in \mathbb{R}^{n_k \times n_k}$ is the diagonal matrix with diagonal entries $\frac{\mu_{ig}^{(k)}}{1+\mu_{ig}^{(k)}\alpha_g}$ for $1 \le i \le n_k$. $\alpha_g$ is the dispersion estimate of the gene and $\mu_{ig}^{(k)}$ is the expected value of the gene for sample for parameter $\beta$, that is $\gamma_g^{(k)} \exp(X_i^{(k)} \cdot \beta_g)$. | Each center | Server |
| | beta | nparray | $(G_{\mathtt{r}}, p)$ | The log fold change $\beta$ parameter. Not modified. | Each center | Server |
| | local_nll | nparray | $(G_{\mathtt{act}},)$ | For each gene $g$, the negative log likelihood of the negative binomial GLM with fixed dispersion at the $\beta_g$ parameter, computed on the local data. | Each center | Server |
| | round_number_irls | int | | The current round number of the IRLS algorithm. Simply passed on. | Each center | Server |
| | global_nll | nparray | $(G_{\mathtt{r}},)$ | For each gene $g$, the negative log likelihood at the $\beta_g$ parameter. Simply passed on. | Each center | Server |
| | irls_mask | nparray | $(G_{\mathtt{r}},)$ | A boolean array indicating which non zero genes are currently being optimized by the IRLS algorithm. Simply passed on. | Each center | Server |
| | irls_diverged_mask | nparray | $(G_{\mathtt{r}},)$ | A boolean array indicating which non zero genes have caused the IRLS algorithm to diverge. Simply passed on | Each center | Server |
| | irls_gene_names | Index | $(G_{\mathtt{act}},)$ | The gene names of the active genes. | Each center | Server |
| | local_features | nparray | $(G_{\mathtt{act}}, p)$ | $\Phi^{(k)} := (X^{(k)})^\top Z^{(k)}$, where $Z^{(k)}$ is the normalized counts in the center $k$ on the set of genes to replace, where the value of Cook's outliers have been replaced using a trimmed mean. | Each center | Server |
| 112 | round_number_irls | int | | The current round number of the IRLS algorithm, incremented by 1 during the local step. | Server | Center |
| | irls_diverged_mask | nparray | $(G_{\mathtt{r}},)$ | A boolean array indicating which non zero genes have caused the IRLS algorithm to diverge. A gene $g$ is considered to have diverged if any component of $\beta_g \in \mathbb{R}^p$ has absolute value above the threshold `max_beta`. | Server | Center |
| | beta | nparray | $(G_{\mathtt{r}}, p)$ | The updated value of the $\beta$ parameter after applying the IRLS update for the active genes. At the last step, is not updated. | Server | Center |
| | irls_mask | nparray | $(G_{\mathtt{r}},)$ | A boolean array indicating which non zero genes are currently being optimized by the IRLS algorithm. Genes whose $\beta$ parameter has diverged are removed from the optimization. Moreover, the deviance ratio is computed for each active gene $g$, as $\frac{\lvert 2\,\mathcal{L}(\beta_g) - 2\cdot\mathcal{L}(\beta_{g,\text{prev}})\rvert}{\lvert 2\,\mathcal{L}(\beta)\rvert + 0.1}$ where $\beta_{g,\text{prev}}$ is the value of $\beta_g$ at the previous iteration. If this deviance is smaller than the user inputed `beta_tol`, the gene is considered to have converged and is set to `False` in the `irls_mask`, meaning it will not be optimized further. | Server | Center |
| | global_nll | nparray | $(G_{\mathtt{r}},)$ | For each gene $g$, the negative log-likelihood at the $\beta_g$ parameter, built by summing the local nlls from the centers | Server | Center |
| 113 | beta | nparray | $(G_{\mathtt{r}}, p)$ | For evert gene $g$, the inital value of the $\beta_g$ parameter for the ProxQuasiNewton algorithm (PQN), which is either i) the value computed by IRLS if IRLS has converged on that gene (in that case, the gene will not be optimized) or ii) the initial value of the $\beta_g$ parameter for the IRLS algorithm, if IRLS has not converged on that gene. | Each center | Server |
| | ascent_direction_on_mask | NoneType | | The ascent direction on the active genes, which is set to `None` at the beginning. | Each center | Server |
| | newton_decrement_on_mask | NoneType | | The newton decrement on the active genes, which is set to `None` at the beginning. | Each center | Server |
| | global_reg_nll | nparray | $(G_{\mathtt{r}},)$ | The regularized negative log likelihood of the negative binomial GLM with fixed dispersion at the $\beta_g$ parameter intialized at `np.nan` since no nll has been computed at this stage. | Each center | Server |
| | PQN_mask | nparray | $(G_{\mathtt{r}},)$ | A boolean array indicating which non zero genes are currently being optimized by the Proximal Quasi-Newton algorithm (active genes). The genes which have diverged during the IRLS algorithm and those which have not converged at the end of the prescribed iterations are set to `True`. | Each center | Server |
| | PQN_diverged_mask | nparray | $(G_{\mathtt{r}},)$ | A boolean array indicating which non zero genes have caused the Proximal Quasi-Newton algorithm to diverge. Initialized to `False`. | Each center | Server |
| | local_gradient | nparray | $(1, G_{\mathtt{act}}, p)$ | For each gene $g$, the gradient of the negative log likelihood of the negative binomial GLM with fixed dispersion at the $\beta_g$ parameter, computed on the local data, i.e., $\nabla_\beta \mathcal{L}^{(k)}(\beta_g) = -(X^{(k)})^\top Y_g^{(k)} + (X^{(k)})^\top \left(\frac{1}{\alpha_g} + Y_g^{(k)}\right) \frac{\mu_g^{(k)}}{\frac{1}{\alpha_g} + \mu_g^{(k)}}$, where $\mu_g^{(k)}$ and $Y_g^{(k)}$ are both vectors in $\mathbb{R}^{n_k}$ and $\alpha_g$ is a scalar. TODO ref equation. | Each center | Server |
| | local_fisher | nparray | $(1, G_{\mathtt{act}}, p, p)$ | For each gene $g$, the Fisher information matrix of the negative binomial GLM with fixed dispersion, $n_k \mathcal{I}_g^{(k)} = (X^{(k)})^\top W_g^{(k)} X^{(k)}$ where $W_g^{(k)} \in \mathbb{R}^{n_k \times n_k}$ is the diagonal matrix with diagonal entries $W_{gii}^{(k)} = \frac{\mu_{ig}^{(k)}}{1+\mu_{ig}^{(k)}\alpha_g}$ for $1 \le i \le n_k$. $\alpha_g$ is the dispersion estimate of the gene and $\mu_{ig}^{(k)}$ is the expected value of the gene $g$ for sample for parameter $\beta_g$, that is $\gamma_g^{(k)} \exp(X_i^{(k)} \cdot \beta_g)$. | Each center | Server |
| | local_nll | nparray | $(1, G_{\mathtt{act}})$ | The nll of the negative binomial GLM with fixed dispersion at the $\beta_g$ parameter, computed on the local data, for each gene $g$. | Each center | Server |
| | irls_diverged_mask | nparray | $(G_{\mathtt{r}},)$ | A boolean array indicating which non zero genes have caused the IRLS algorithm to diverge. Simply passed on during the Fed Proximal Quasi-Newton algorithm, since the server is stateless. | Each center | Server |
| | round_number_PQN | int | | The current round number of the Proximal Quasi-Newton algorithm, initialized at 0. | Each center | Server |

| | | | | | | |
|---|---|---|---|---|---|---|
| 114 | beta | nparray | $(G_{\texttt{r}}, p)$ | For each gene $g$, the log fold change $\beta_g$, which remains unchanged during the first iteration of the Proximal Quasi-Newton algorithm. | Server | Center |
| | global_reg_nll | nparray | $(G_{\texttt{r}}, )$ | The regularized negative log likelihood of the negative binomial GLM with fixed dispersion at the $\beta_g$ parameter computed as the sum of the local negative log likelihoods with $L^2$ regularization with $\lambda = 10^{-6}$. | Server | Center |
| | round_number_PQN | int | | The current round number of the Proximal Quasi-Newton algorithm, incremented by 1. | Server | Center |
| | newton_decrement_on_mask | nparray | $(G_{\texttt{act}}, )$ | The newton decrement on the active genes. It is computed as $\nu_g = \Delta_g \cdot \nabla_\beta \mathcal{L}^\lambda(\beta_g)$ where $\Delta_g$ is the ascent direction on the active genes and $\mathcal{L}^\lambda$ is the regularized log likelihood. | Server | Center |
| | ascent_direction_on_mask | nparray | $(G_{\texttt{act}}, p)$ | The ascent direction on the active genes. To compute the ascent direction, we first compute the global Fisher information matrix for each gene $g$, $n\mathcal{I}_g = \sum_k n_k \mathcal{I}_g^{(k)}$ and the global gradient $\nabla_\beta \mathcal{L}(\beta_g) = \sum_k \nabla_\beta \mathcal{L}^{(k)}(\beta_g)$. From these two quantities, we build the Fisher information and gradients of the regularized negative log likelihood, with $L^2$ regularization $\lambda = 10^{-6}$. We add another regularization term to the Fisher matrix which depends on the iteration number. The goal is for the ascent direction to be close to the gradient for small iterations and to be close to the real natural gradient descent update near the end of the optimization. The ascent direction is then computed as from the regularized Fisher and gradient (see main paper). Roughly, it is the solution of the linear system $n\, \mathcal{I}_g^\lambda \Delta_g = \nabla_\beta \mathcal{L}^\lambda(\beta_g)$, where certain components are dropped in to handle boundary conditions as the optimization is constrained to values of $\beta_g$ in $[-\texttt{max\_beta}, \texttt{max\_beta}]$. | Server | Center |
| | PQN_diverged_mask | nparray | $(G_{\texttt{r}}, )$ | A boolean array indicating which non zero genes have caused the Proximal Quasi-Newton algorithm to diverge. Passed on without modification. | Server | Center |
| | PQN_mask | nparray | $(G_{\texttt{r}}, )$ | A boolean array indicating which non zero genes are currently being optimized by the Proximal Quasi-Newton algorithm (active genes), passed on without modification. | Server | Center |
| | irls_diverged_mask | nparray | $(G_{\texttt{r}}, )$ | A boolean array indicating which non zero genes have caused the IRLS algorithm to diverge. Passed on without modification. | Server | Center |
| 115 | local_gradient | nparray | $(N_{\texttt{ls}}, G_{\texttt{act}}, p)$ | For each potential step size $\delta$ and each gene $g$, the gradient of the negative log likelihood of the negative binomial GLM with fixed dispersion at the $\beta_g - \delta\Delta_g$ parameter computed on the local data. | Each center | Server |
| | local_fisher | nparray | $(N_{\texttt{ls}}, G_{\texttt{act}}, p, p)$ | For each potential step size $\delta \in \{1, 1/2, 1/4, ..., 1/2^{N_{\texttt{ls}}-1}\}$, and each gene $g$, the Fisher information matrix of the negative binomial GLM with fixed dispersion computed at $\beta_g - \delta\Delta_g$. Formally, $n_k \mathcal{I}_{\delta,g}^{(k)} = (X^{(k)})^\top W_{\delta,g}^{(k)} X^{(k)}$ where $W_{\delta,g}^{(k)} \in \mathbb{R}^{n_k \times n_k}$ is the diagonal matrix with diagonal entries $W_{\delta,gii}^{(k)} = \frac{\mu_{\delta,ig}^{(k)}}{1+\mu_{\delta,ig}^{(k)}\alpha_g}$ for $1 \leq i \leq n_k$. $\alpha_g$ is the dispersion estimate of the gene and $\mu_{\delta,ig}^{(k)}$ is the expected value of the gene $g$ for sample $i$ for parameter $\beta_g - \delta\Delta_g$, that is $\gamma_g^{(k)} \exp(X_i^{(k)} \cdot (\beta_g - \delta\Delta_g))$. | Each center | Server |
| | local_nll | nparray | $(N_{\texttt{ls}}, G_{\texttt{act}})$ | The nll of the negative binomial GLM with fixed dispersion at the $\beta_g$ parameter, computed on the local data, for each gene $g$ and for each potential next step $\beta_g - \delta\Delta_g$ for $\delta \in \{1, 1/2, 1/4, ..., 1/2^{N_{\texttt{ls}}-1}\}$. | Each center | Server |
| | beta | nparray | $(G_{\texttt{r}}, p)$ | For each gene $g$, the log fold change $\beta_g$, which is simply passed on. | Each center | Server |
| | ascent_direction_on_mask | nparray | $(G_{\texttt{act}}, p)$ | The ascent direction on the active genes, passed on without modification. | Each center | Server |
| | irls_diverged_mask | nparray | $(G_{\texttt{r}}, )$ | A boolean array indicating which non zero genes have caused the IRLS algorithm to diverge. Passed on without modification. | Each center | Server |
| | round_number_PQN | int | | The current round number of the Proximal Quasi-Newton algorithm, passed on without modification. | Each center | Server |
| | newton_decrement_on_mask | nparray | $(G_{\texttt{act}}, )$ | The newton decrement on the active genes, passed on without modification. | Each center | Server |
| | global_reg_nll | nparray | $(G_{\texttt{r}}, )$ | The regularized negative log likelihood of the negative binomial GLM with fixed dispersion at the $\beta_g$ parameter simply passed on. | Each center | Server |
| | PQN_mask | nparray | $(G_{\texttt{r}}, )$ | A boolean array indicating which non zero genes are currently being optimized by the Proximal Quasi-Newton algorithm (active genes), passed on without modification. | Each center | Server |
| | PQN_diverged_mask | nparray | $(G_{\texttt{r}}, )$ | A boolean array indicating which non zero genes have caused the Proximal Quasi-Newton algorithm to diverge. Passed on without modification. | Each center | Server |
| 116 | ascent_direction_on_mask | nparray | $(G_{\texttt{act}}, p)$ | The ascent direction on the new active genes $g$, computed at the new $\beta_g$ iterate. | Server | Center |
| | PQN_mask | nparray | $(G_{\texttt{r}}, )$ | A boolean array indicating which non zero genes are currently being optimized by the Proximal Quasi-Newton algorithm (active genes). Genes that had converged with the relative error criterion or diverged (no stepsize satisfying the Armijo condition) have set to `False`. | Server | Center |
| | round_number_PQN | int | | The current round number of the Proximal Quasi-Newton algorithm, incremented by 1. | Server | Center |
| | global_reg_nll | nparray | $(G_{\texttt{r}}, )$ | The regularized negative log likelihood of the negative binomial GLM with fixed dispersion at the $\beta_g$ parameter computed as the sum of the local negative log likelihoods with $L^2$ regularization with $\lambda = 10^{-6}$. | Server | Center |
| | irls_diverged_mask | nparray | $(G_{\texttt{r}}, )$ | A boolean array indicating which non zero genes have caused the IRLS algorithm to diverge. Passed on without modification. | Server | Center |
| | beta | nparray | $(G_{\texttt{r}}, p)$ | For each gene $g$, the log fold change $\beta_g$. This value has been updated by the Proximal Quasi-Newton algorithm for all active genes where an admissible step size was found using the Armijo condition (see equation 2.5 of [1]). | Server | Center |
| | PQN_diverged_mask | nparray | $(G_{\texttt{r}}, )$ | A boolean array indicating which non zero genes have caused the Proximal Quasi-Newton algorithm to diverge. A gene $g$ is considered to have diverged if for all step sizes $\delta \in \{1, 1/2, 1/4, ..., 1/2^{N_{\texttt{ls}}-1}\}$, and for the computed ascent direction $\Delta_g$, the Armijo condition is not satisfied, i.e., if $\mathcal{L}^\lambda(\beta_g) - \mathcal{L}^\lambda(\beta_g - \delta\Delta_g) \leq \delta\ \texttt{PQN\_c1}\ \nabla_\beta \mathcal{L}^\lambda(\beta_g) \cdot \Delta_g$ where $\mathcal{L}^\lambda$ is the regularized negative log likelihood, `PQN_c1` is the Armijo parameter which can be set by the user (default is $10^{-4}$), and $\nabla_\beta \mathcal{L}^\lambda(\beta_g)$ is the gradient of the regularized negative log likelihood. | Server | Center |
| | newton_decrement_on_mask | nparray | $(G_{\texttt{act}}, )$ | The newton decrement on the new active genes $g$, computed at the new $\beta_g$ iterate. For more details, see description of 19. | Server | Center |

| | | | | | | |
|---|---|---|---|---|---|---|
| 117 | newton_decrement_on_mask | nparray | $(G_{\texttt{act}},)$ | The newton decrement on the active genes, passed or without modification. | Each center | Server |
| | local_fisher | nparray | $(N_{\texttt{ls}}, G_{\texttt{act}}, p, p)$ | For each potential step size $\delta \in \{1, 1/2, 1/4, ..., 1/2^{N_{\texttt{ls}}-1}\}$, and each gene $g$, the Fisher information matrix of the negative binomia GLM with fixed dispersion computed at $\beta_g - \delta\Delta_g$ Formally, $n_k\mathcal{I}^{(k)}_{\delta,g} = (X^{(k)})^\top W^{(k)}_{\delta,g} X^{(k)}$ where $W^{(k)}_{\delta,g} \in \mathbb{R}^{n_k \times n_k}$ is the diagonal matrix with diag- onal entries $W^{(k)}_{\delta,gii} = \frac{\mu^{(k)}_{\delta,ig}}{1+\mu^{(k)}_{\delta,ig}\alpha_g}$ for $1 \leq i \leq n_k$. $\alpha_g$ is the dispersion estimate of the gene and $\mu^{(k)}_{\delta,ig}$ is the expected value of the gene $g$ for sample $i$ for parameter $\beta_g - \delta\Delta_g$, that is $\gamma^{(k)}_g \exp(X^{(k)}_i \cdot (\beta_g - \delta\Delta_g))$. | Each center | Server |
| | beta | nparray | $(G_{\texttt{r}}, p)$ | For each gene $g$, the log fold change $\beta_g$, which is simply passed on. | Each center | Server |
| | local_nll | nparray | $(N_{\texttt{ls}}, G_{\texttt{act}})$ | The nll of the negative binomial GLM with fixed disper- sion at the $\beta_g$ parameter, computed on the local data, for each gene $g$ and for each potential next step $\beta_g - \delta\Delta_g$ for $\delta \in \{1, 1/2, 1/4, ..., 1/2^{N_{\texttt{ls}}-1}\}$. | Each center | Server |
| | local_gradient | nparray | $(N_{\texttt{ls}}, G_{\texttt{act}}, p)$ | For each potential step size $\delta$ and each gene $g$, the gradi- ent of the negative log likelihood of the negative binomia GLM with fixed dispersion at the $\beta_g - \delta\Delta_g$ parameter computed on the local data. | Each center | Server |
| | PQN_diverged_mask | nparray | $(G_{\texttt{r}},)$ | A boolean array indicating which non zero genes have caused the Proximal Quasi-Newton algorithm to diverge Passed on without modification. | Each center | Server |
| | PQN_mask | nparray | $(G_{\texttt{r}},)$ | A boolean array indicating which non zero genes are cur- rently being optimized by the Proximal Quasi-Newton al- gorithm (active genes), passed on without modification. | Each center | Server |
| | global_reg_nll | nparray | $(G_{\texttt{r}},)$ | The regularized negative log likelihood of the negative binomial GLM with fixed dispersion at the $\beta_g$ parameter simply passed on. | Each center | Server |
| | round_number_PQN | int | | The current round number of the Proximal Quasi-Newtor algorithm, passed on without modification. | Each center | Server |
| | irls_diverged_mask | nparray | $(G_{\texttt{r}},)$ | A boolean array indicating which non zero genes have caused the IRLS algorithm to diverge. Passed on with- out modification. | Each center | Server |
| | ascent_direction_on_mask | nparray | $(G_{\texttt{act}}, p)$ | The ascent direction on the active genes, passed on with- out modification. | Each center | Server |
| 118 | beta | nparray | $(G_{\texttt{r}}, p)$ | For each gene $g$, the log fold change $\beta_g$. This value has been updated by the Proximal Quasi-Newton algorithm for all active genes where an admissible step size was found using the Armijo condition (see equation 2.5 of [1]) | Server | Center |
| | irls_diverged_mask | nparray | $(G_{\texttt{r}},)$ | A boolean array indicating which non zero genes have caused the IRLS algorithm to diverge. Passed on with- out modification. | Server | Center |
| | PQN_diverged_mask | nparray | $(G_{\texttt{r}},)$ | A boolean array indicating which non zero genes have caused the Proximal Quasi-Newton algorithm to diverge As this is the last step of the optimization, a gene $g$ is con- sidered to have diverged not only if the Armijo conditior is never satisfied, but also if the convergence criterion is not met. A gene is said to converge if the relative dif- ference between two successive iterates is smaller than a threshold value `PQN_ftol`, i.e., $\lvert \mathcal{L}^\lambda(\beta_g) - \mathcal{L}^\lambda(\beta_{g,\text{prev}})\rvert \leq$ `PQN_ftol` $\max(\mathcal{L}^\lambda(\beta_g), \mathcal{L}^\lambda(\beta_{g,\text{prev}}), 1)$. | Server | Center |
| 121 | n_samples | int | | The number of samples in a center $n_k$ for each center $k$. | Each center | Server |
| | mean_normed_counts | nparray | $(G,)$ | For each gene, the mean of the local normed counts, i.e. $\overline{Z}^{(k)}_g = \frac{1}{n_k}\sum_{i=1}^{n_k} Z^{(k)}_{ig}$. | Each center | Server |
| | local_hat_matrix | nparray | $(G_{\texttt{nz}}, p, p)$ | For each gene $g$, the hat matrix $H^{(k)}_g = (X^{(k)})^\top W^{(k)}_g X^{(k)}$ where $W^{(k)}_g \in \mathbb{R}^{n_k\times n_k}$ is the diagonal matrix with diagonal entries $\frac{\mu^{(k)}_{ig}}{1+\mu^{(k)}_{ig}\alpha_g}$ for $1 \leq i \leq n_k$. $\alpha_g$ is the dispersion estimate of the gene and $\mu^{(k)}_{ig}$ is the expected value of the gene for sample $i$ for parameter $\beta$, that is $\gamma^{(k)}_g \exp(X^{(k)}_i \cdot \beta_g)$. | Each center | Server |
| | contrast_vector | nparray | $(p,)$ | The contrast vector to apply to the LFC to get the desirec log fold change corresponding to the input contrast of the form Of the form `[factor, level1, level2]`. | Each center | Server |
| | LFC | DataFrame | $(G, p)$ | The log fold changes of the gene expression. This dataframe is indexed by genes on one hand, and by desigr column names on the other. | Each center | Server |
| | local_H_matrix | nparray | $(G, p, p)$ | The hat matrix with `nan` entries for zero genes, used to compute Wald statistics. Should be merged with hat ma- trix. | Each center | Server |
| | varEst | nparray | $(G,)$ | For each gene $g$, the trimmed variance estimate of the normed counts, denoted with $V^{\texttt{trim}}_g$. This quantitiy is retrieved from the current local state. | Each center | Server |
| | _skip_cooks | bool | | A boolean indicating whether to skip the computation of the intermediate quantities to compute the of the Cook's distance. This is set to `True` if the Cook's distance is storec in the local state (which is not the case by default due to memory issues). Otherwise, it is set to `False` (default behaviour). | Each center | Server |
| 122 | p_values | nparray | $(G,)$ | For each gene, the p-value of the Wald statistic, computed from the survival function of the normal distribution ap- plied to the wald statistic. | Server | Center |
| | wald_statistics | nparray | $(G,)$ | For each gene, the Wald statistic of the gene expression This statistics depends on the `lfc_null` parameter which sets the null hypothesis on the log fold change (set to 0 by default), and the `alt_hypothesis` parameter, which defines the alternative hypothesis on the log fold change (set to `None` by default, can be `greater`, `greaterAbs`, `less`, `lessAbs`). If the alternative hypothesis is `None`, then the Wald statistic is computed as the centered normalized log fold change. | Server | Center |
| | wald_se | nparray | $(G,)$ | For each gene, the standard error on the log fold change value for the given contrast given by the GLM. | Server | Center |
| | cooks_dispersions | nparray | $(G,)$ | For each gene $g$, a robust estimate of the dispersion pa- rameter $\alpha^{\texttt{cooks}}_g$ computed from the trimmed variance es- timate and the global mean of the normed counts. We compute this estimate as $\max((V^{\texttt{trim}}_g - \overline{Y}_g)/\overline{Y}^2_g, 0.04)$. | Server | Center |
| | global_hat_matrix_inv | nparray | $(G_{\texttt{nz}}, p, p)$ | For each gene $g$, we compute the global hat matrix as the sum of the local hat matrices, and its inverse. | Server | Center |
| 123 | _skip_cooks | bool | | A boolean indicating whether to skip the computation of the intermediate quantities to compute the of the Cook's distance. This is set to `True` if the Cook's distance is storec in the local state (which is not the case by default due to memory issues). Otherwise, it is set to `False` (default behaviour). | Each center | Server |
| | n_samples | int | | The number of samples in a center $n_k$ for each center $k$. | Each center | Server |

| | | | | | | |
|---|---|---|---|---|---|---|
| | local_cooks_outliers | nparray | $(G,)$ | A boolean array which for each gene $g$ indicates if the Cook's distance is above the cutoff value for any sample in the center which i) has not been replaced by an imputation value if Cook's outliers have been refitted and ii) has at least three replicates across all centers. | Each center | Server |
| | cooks_cutoff | float | () | The cutoff value for the Cook's distance, set to the 0.99-th quantile of the F-distribution with $p$ and $n-p$ degrees of freedom. | Each center | Server |
| | local_hat_matrix | nparray | $(G_{\texttt{nz}}, p, p)$ | For each gene $g$, the hat matrix $H_g^{(k)} = (X^{(k)})^\top W_g^{(k)} X^{(k)}$ where $W_g^{(k)} \in \mathbb{R}^{n_k \times n_k}$ is the diagonal matrix with diagonal entries $\frac{\mu_{ig}^{(k)}}{1+\mu_{ig}^{(k)}\alpha_g}$ for $1 \leq i \leq n_k$. $\alpha_g$ is the dispersion estimate of the gene and $\mu_{ig}^{(k)}$ is the expected value of the gene for sample $i$ for parameter $\beta$, that is $\gamma_g^{(k)} \exp(X_i^{(k)} \cdot \beta_g)$. | Each center | Server |
| | mean_normed_counts | nparray | $(G,)$ | For each gene, the mean of the local normed counts, i.e. $\overline{Z}_g^{(k)} = \frac{1}{n_k}\sum_{i=1}^{n_k} Z_{ig}^{(k)}$. | Each center | Server |
| | varEst | nparray | $(G,)$ | For each gene $g$, the trimmed variance estimate of the normed counts, denoted with $V_g^{\texttt{trim}}$. This quantitiy is retrieved from the current local state. | Each center | Server |
| 124 | cooks_outliers | nparray | $(G,)$ | A boolean array which for each gene $g$ indicates if the gene is a local Cook's outlier in any center. Computed from the local cooks outliers across all centers. | Server | Center |
| | cooks_dispersions | nparray | $(G,)$ | For each gene $g$, a robust estimate of the dispersion parameter $\alpha_g^{\texttt{cooks}}$ computed from the trimmed variance estimate and the global mean of the normed counts. We compute this estimate as $\max((V_g^{\texttt{trim}} - \overline{Y}_g)/\overline{Y}_g^2, 0.04)$. | Server | Center |
| | global_hat_matrix_inv | nparray | $(G_{\texttt{nz}}, p, p)$ | For each gene $g$, we compute the global hat matrix as the sum of the local hat matrices, and its inverse. | Server | Center |
| | cooks_cutoff | float | () | The cutoff value for the Cook's distance, set to the 0.99-th quantile of the F-distribution with $p$ and $n-p$ degrees of freedom. | Server | Center |
| 125 | cooks_outliers | nparray | $(G,)$ | A boolean array which for each gene $g$ indicates if the gene is a Cook's outlier in any center. Passed on without modification. | Each center | Server |
| | local_max_cooks | nparray | $(G_{\texttt{act}},)$ | For each gene $g$ which has a Cook's outlier, the maximum Cook's distance across all samples in the center, if there is one above the cutoff value. Otherwise, 0. | Each center | Server |
| 126 | cooks_outliers | nparray | $(G,)$ | A boolean array which for each gene $g$ indicates if the gene is a Cook's outlier in any center. Passed on without modification. | Server | Center |
| | max_cooks | nparray | $(G_{\texttt{act}},)$ | For each gene $g$ which has a Cook's outlier, the maximum Cook's distance across all samples in all centers, if there is one above the cutoff value. Otherwise, 0. Computed from the local max cooks across all centers. | Server | Center |
| 127 | cooks_outliers | nparray | $(G,)$ | A boolean array which for each gene $g$ indicates if the gene is a Cook's outlier in any center. Passed on without modification. | Each center | Server |
| | local_max_cooks_gene_counts | MaskedArray | $(G_{\texttt{act}},)$ | A masked array where for each gene $g$ with a Cook's outliers, the gene is masked if the sample maximizing the Cook's distance is not in the center, and the count value of the sample maximizing the Cook's distance otherwise. | Each center | Server |
| 128 | max_cooks_gene_counts | MaskedArray | $(G_{\texttt{act}},)$ | A masked array where for each gene $g$ with a Cook's outliers, we have the minimum of the counts maximizing the Cook's distance across all centers. Should not be masked. | Server | Center |
| | cooks_outliers | nparray | $(G,)$ | A boolean array which for each gene $g$ indicates if the gene is a Cook's outlier in any center. Passed on without modification. | Server | Center |
| 129 | wald_se | nparray | $(G,)$ | For each gene, the standard error on the log fold change value for the given contrast given by the GLM. Stored in the local state. | Each center | Server |
| | wald_statistics | nparray | $(G,)$ | For each gene, the Wald statistic of the gene expression. Stored in the local state. | Each center | Server |
| | p_values | nparray | $(G,)$ | For each gene, the p-value of the Wald statistic, computed from the survival function of the normal distribution applied to the wald statistic. Stored in the local state. | Each center | Server |
| | cooks_outliers | nparray | $(G,)$ | A boolean array which for each gene $g$ indicates if the gene is a Cook's outlier in any center. Passed on without modification. | Each center | Server |
| | local_num_samples_above | MaskedArray | $(G_{\texttt{act}},)$ | For each gene $g$ which has a Cook's outlier, the number of samples in the center whose gene count for gene $g$ is above the gene count of the sample maximizing the Cook's distance across all centers for that gene. | Each center | Server |
| 130 | wald_se | nparray | $(G,)$ | For each gene, the standard error on the log fold change value for the given contrast given by the GLM. Passed on without modification. | Server | Center |
| | wald_statistics | nparray | $(G,)$ | For each gene, the Wald statistic of the gene expression. Passed on without modification. | Server | Center |
| | p_values | nparray | $(G,)$ | If Cook's filtering is enabled (which is the case by default with the `cooks_filter` parameter), then the p-values of genes which have $\leq 2$ samples above the gene count of the sample maximizing the Cook's distance across all centers are set to `nan`. Otherwise, passed on without modification. | Server | Center |
| 132 | contrast | list | | A list of three strings representing the contrast of interest, in case it is not specified by the user. Of the form `[factor, level1, level2]`. For example `[stage, Advanced, Non-advanced]`. | Each center | Server |
| | _squared_logres | float | () | The squared mean absolute deviation of the difference between the log of the genewise dispersions and the log of the fitted dispersions, restricted to the non-zero genes whose gene-wise dispersions are above $100 \times$ `min_disp`. The mean absolute deviation estimate is defined as the median of the absolute difference between the log residual and its median, scaled by the percent point function of the normal distribution at 0.75. | Each center | Server |
| | prior_disp_var | float | () | A prior on the variance of the log dispersions around the log trend curve $\sigma^2_{\texttt{trend}}$, estimated as the maximum between 0.25 and `_squared_log_res` $- \psi_1((n-p)/2)$, where $\psi_1(f/2)$ is the variance of the log of a $\chi^2_f$ distribution. For more details, see [2]. | Each center | Server |
| | wald_se | nparray | $(G,)$ | For each gene, the standard error on the log fold change value for the given contrast given by the GLM. | Each center | Server |
| | replaced | nparray | $(G,)$ | A boolean array marking genes $g$ for which the Cook's distance is above the cutoff value for any sample in any center (the union of the local genes to replace across all centers). | Each center | Server |
| | wald_statistics | nparray | $(G,)$ | For each gene, the Wald statistic of the gene expression. | Each center | Server |
| | p_values | nparray | $(G,)$ | For each gene, the p-value of the Wald statistic, computed from the survival function of the normal distribution applied to the wald statistic, and set to `nan` if the gene is a Cook's outlier if `cooks_filter` is enabled. | Each center | Server |
